# Expanding CyanoHAB Monitoring: New Micropeptins and Generalizable MS/MS Workflows for the Annotation of Cyanopeptide Classes

**DOI:** 10.64898/2026.02.07.704577

**Authors:** Runjie Xia, Lindsey Ahn, Michaela Burkhauser, Ross Youngs, Matthew J. Bertin

**Affiliations:** Department of Chemistry, Case Western Reserve University, Cleveland, OH 44106, United States; Biosortia, Inc., 2545 Farmers Dr., Suite 370, Columbus, OH 43235, United States

**Keywords:** cyanoHABs, cyanopeptides, micropeptins, LC-MS/MS, protease inhibition

## Abstract

Cyanobacterial harmful algal blooms (cyanoHABs) are a major ecological and public health concern, commonly monitored for hepatotoxic microcystins and cylindrospermopsins and neurotoxic anatoxins and saxitoxins. However, the broader suite of bioactive metabolites produced during blooms remains under characterized. Here, we interrogated a chromatography fraction library generated from a cyanoHAB in Muskegon, Michigan. From this library, we isolated two new micropeptins (**1** and **2**), including an analog bearing a bishomologated tyrosine residue, and we confirmed the structure of ferintoic acid C (**3**). Structures were established using complementary spectrometric and spectroscopic methods. To expand chemical space coverage beyond isolated compounds, we analyzed LC-MS/MS data using the GNPS2 Analysis Hub query language for product ion searching, enabling annotation of cyanopeptide classes and class-specific modifications across the fraction set, which provided a practical and user-friendly approach for identifying cyanopeptide classes. One of the new micropeptins (**1**) exhibited moderate inhibition of neutrophil elastase, consistent with roles in ecological interactions and potential relevance to human exposure. Analysis of field samples from ongoing Lake Erie blooms showed recurring micropeptins but no evidence of microcystins. Together, these results challenge microcystin-centric assessments of bloom hazard and support expanded monitoring of non-microcystin cyanopeptides.

**SYNOPSIS:** Routine cyanoHAB monitoring targets few regulated toxins; we reveal abundant, under characterized cyanopeptides and enable rapid class-level annotation across datasets with a new LC-MS/MS analysis pipeline.

**GRAPHICAL ABSTRACT:** 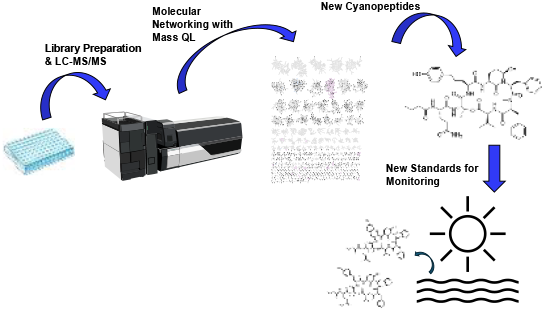

## INTRODUCTION

Cyanobacterial harmful algal blooms (cyanoHABs) are expanding globally and represent a persistent threat to water quality and human health through the production of toxic secondary metabolites.^1,2^ In most freshwater systems, monitoring and risk management are centered on microcystins, a structurally diverse family of hepatotoxins that is the basis for international guidance values and U.S. drinking-water health advisories.^3,4^ However, the current monitoring approach does not capture the broader metabolite mixtures produced during blooms, which is increasingly being shown in more and more studies.^5-7^

Cyanobacteria generate extensive suites of “cyanometabolites”, including numerous cyanopeptides assembled by nonribosomal peptide synthetases and other biosynthetic pathways, such as micropeptins/cyanopeptolins, anabaenopeptins, microviridins, microginins, and others in addition to the large class of microcystins. These compounds can exhibit potent bioactivities and may influence ecological interactions, food-web transfer, and human exposure profiles.^8^ A growing body of evidence indicates that non-microcystin cyanopeptides can occur as frequently as microcystins and sometimes at similar concentrations in surface waters, yet their environmental distributions, toxicity, and risk relevance remain less defined.^7-9^ This imbalance reflects practical barriers: high structural diversity, limited availability of reference standards, and analytical complexity that challenges routine targeted monitoring. Consequently, there is a need for workflows that both enable structure-level characterization of newly observed congeners and expand class-level screening across complex bloom events.

High-resolution LC–MS/MS paired with open computational resources offers a scalable approach to address these challenges. GNPS provides widely used infrastructure for organizing and annotating untargeted MS/MS data, including molecular networking and repository-scale spectral interpretation.^10^ GNPS2 extends these capabilities through a web-based analysis hub and tools designed for accessible interrogation of MS/MS datasets including a newly available query language to assess a multitude of molecular features such as halogenation and product ion composition.^11,12^ Such platforms can facilitate class-level annotation via diagnostic fragments and product ion patterns, complementing targeted quantification and isolation-based structure elucidation.

Here, we apply an integrated discovery and annotation workflow to a cyanoHAB from Muskegon, Michigan. We interrogated a chromatography fraction library, enabling both targeted isolation for rigorous structural characterization and untargeted LC–MS/MS analysis for broader metabolite annotation. Using complementary spectrometric and spectroscopic methods, we report two new micropeptins (**1** and **2**) and confirm the structure of ferintoic acid C (**3**), a metabolite previously assigned via MS/MS spectrum interpretation.^13^ We then leveraged GNPS2 product ion searching to annotate cyanopeptide classes (e.g., microcystins, anabaenopeptins, microviridins, micropeptins, microginins etc.) and modifications across the fraction set, which allowed for additional mass spectrometry based characterizations. Finally, we assessed protease inhibitory activities and examined field relevance by surveying Lake Erie bloom samples to evaluate whether micropeptins represent an abundant, recurring component of Great Lakes cyanoHAB chemical profiles. This work ultimately provides an integrated platform of mass spectrometry-driven class-level and new metabolite annotation in field samples and unequivocal identification of emerging cyanometabolites by chromatographic comparison to a standard library of rare cyanopeptides.

## MATERIALS AND METHODS

### LC-MS/MS analysis and molecular networking

We purchased a library of preparative chromatography fractions from Biosortia Microbiomics (20 fractions). The library was derived from a cyanobacterial bloom originally collected from Muskegon, MI. This library resulted from extraction and fractionation procedures that have been described previously,^9^ and we have explored it previously using bioassay-guidance to characterize new microcystins and new micropeptins and other cyanopeptides.^5,14^ The individual fractions were reconstituted in LC-MS grade CH_3_OH and each fraction (10 µL injection volume) was subjected to LC-MS/MS analysis on an Agilent Revident QToF mass spectrometer with a 1290 Infinity II Bio LC and column for analytical separations. Analyte separation was performed using an Eclipse Plus C18 column (1.8 μm, 50 mm × 2.1 mm) with a flow rate of 0.4 mL/min. Mobile phases consisted of H_2_O with 0.1% formic acid (A) and CH_3_CN with 0.1% formic acid (B). Untargeted HRMS and data-dependent MS/MS acquisition were conducted in positive ion mode. The gradient method was as follows: 80% A and 20% B held for 5 min, followed by a linear increase of B to 80% over 20 min (5–25 min), then to 90% B over the next 5 min, with a return to initial conditions at 30.01 min. Full-scan MS data were acquired over an *m/z* range of 200–1700, and MS/MS data were acquired over an *m/z* range of 100–1700. MS/MS spectra were collected in Auto MS/MS mode using a collision energy of 40 V at an acquisition rate of 3 spectra s^-1^, with up to four precursor ions selected per cycle and an isolation width of 1–3 *m/z*. Source parameters were as follows: gas temperature, 350°C; capillary temperature, 300°C; spray voltage, 3.5 kV; sheath gas flow rate, 11 L min^−1^. Targeted LC-MS/MS analysis of micropeptins used a modified gradient and the same solvents as above consisting of a 5 min isocratic hold at 95% A, followed by a 15 min linear gradient to 100% B, a 5 min hold at 100% B, and re-equilibration starting at 25.01 min. Full-scan MS and MS/MS data were acquired over an *m/z* range of 20–2000. Raw data files were exported as mzXML files and uploaded to the GNPS2 website using the classical molecular networking workflow and the default parameters, except that precursor *m/z* tolerance and fragment *m/z* tolerance were both set to 0.02 Da. Next after the network was generated, we used the Mass QL^12^ (query language) function available at GNPS2 to search for product ions associated with different cyanopeptide classes (e.g., *m/z* 282 and *m/z* 404 for micropeptins/cyanopeptolins) and product ions for other reported cyanopeptide groups such as the microcystins. Several new micropeptin structures were proposed based on the analysis of MS/MS spectra, details of which are described in the Results and Discussion section below. Following this first network generation via a small subset of chromatography fractions, we acquired a larger number of chromatography fractions from the Muskegon, MI cyanoHAB collection from Biosortia Microbiomics, which covered a much larger polarity range and contained a more complex portfolio of cyanometabolites. These fractions were subjected to the identical tandem mass spectrometry workflow and product ion searches were conducted to identify different classes of cyanopeptides.

### Isolation of 1-3 and [Leu^1^]MC-LR

Preparative amounts of certain chromatography fractions were purchased from Biosortia Microbiomics, which corresponded to the samples analyzed by LC-MS/MS. Following the molecular networking procedures, these fractions were profiled by LC-MS to demonstrate that certain analytes of interest (e.g., *m/z* 993 [M+H-H_2_O]^+^, micropeptin 1010) from the molecular networks were present. Micropeptin 1010 (**1**) was isolated from a chromatography fraction using a Kinetex C18 column (5 *μ*m, 250 mm × 10 mm) with H_2_O modified with 0.1% formic acid (solvent A) and CH_3_CN modified with 0.1% formic acid (solvent B). Separation was performed at a flow rate of 3.00 mL/min under isocratic conditions of 65% A and 35% B. A single peak was collected, yielding 4.7 mg of compound **1** (t_R_ 3.4 min).

A second chromatography fraction was reconstituted in CH_3_OH at a concentration of 10 mg/mL and subjected to separation using a YMC ODS column (5 *μ*m, 250 mm × 10 mm). Isocratic elution was performed with 55% A and 45% B at a flow rate of 3.00 mL/min. Seven peaks were collected and analyzed by LC-MS. LC-MS analysis indicated that the first peak consisted of two coeluting components. Repurification of this fraction was performed on a Kinetex C18 column (2.6 μm, 150 mm × 4.6 mm) using an isocratic method consisting of 65% A and 35% B. A subfraction was collected, yielding 0.8 mg of micropeptin 966 (**2**) (t_R_ 9.5 min). The remaining 6 peaks contained known micropeptins and previously characterized compounds (determined via ^1^H NMR, HRMS, and t_R_ values): 0.6 mg of micropeptin 982 (t_R_ 6.3 min, peak 2), 1.0 mg ferintoic acid A (t_R_ 7.4 min, peak 3), 1.0 mg of ferintoic acid C (t_R_ 8.5 min, peak 4, **3**), and 0.9 mg of micropeptin 982 (L-Ser) (t_R_ 9.0 min, peak 5). Peaks 6 and 7 from the initial separation were combined and further purified using the same YMC ODS column under isocratic conditions with 60% A and 40% B. This final purification yielded an 8.4 mg mixture of micropeptin 957 and micropeptin 982 (D-Gln) and a 6.4 mg mixture of micropeptin 950 and micropeptin 996. While ferintoic acid C (**3**) has been characterized via MS/MS,^13^ it has not been unequivocally characterized using NMR, which we accomplished in this report.

A third chromatography fraction with an abundant *m/z* 1037 ion signal was subjected to reversed-phase semi-preparative HPLC using a Kinetex C18 column (5 μm, 250 mm × 10 mm) with the same A and B solvents as detailed above. Separations were performed at a flow rate of 3.00 mL/min with starting conditions of 68% A and 32% B and a gradient as follows: 32% B to 42% B from 0-30 min and 42% B to 100% B from 35 to 40 min, returning to initial conditions at 41 min. A single peak was collected, yielding 6 mg of [Leu^1^]MC-LR (t_R_ 20.76 min).

*Micropeptin 1010* (**1**): colorless oil; [*α*]^21^_D_ = −27.4 (*c* 0.05, MeOH); UV *λ*_max_: 223, 280 nm (from HPLC-DAD); ^1^H NMR (500 MHz, DMSO-*d*_6_) and ^13^C NMR (125 MHz, DMSO-*d*_6_) see Table S1, HRMS (ESI-QTOF) *m/z* calcd for C_53_H_70_N_8_O_12_Na^+^: 1033.5005 [M+Na]^+^; found 1033.5009 (error 0.39 ppm).

*Micropeptin 966 (D-Gln)* (**2**): colorless oil; UV *λ*_max_: 280 nm (from HPLC-DAD); ^1^H NMR (500 MHz, DMSO-*d*_6_) and ^13^C NMR (125 MHz, DMSO-*d*_6_) see Table S2, HRMS (ESI-QTOF) *m/z* calcd for C_51_H_66_N_8_O_11_Na^+^: 989.4743 [M+Na]^+^; found 989.4771 (error 2.83 ppm).

*Ferintoic acid C* (**3**): colorless oil; UV *λ*_max_: 220, 280 nm (from HPLC-DAD); ^1^H NMR (500 MHz, DMSO-*d*_6_) *δ* Trp^1^: 4.00 (CH, m), 3.15, 2.99 (CH_2_, m), 7.48 (CH, d, *J*=7.9 Hz), 6.90 (CH, t, *J*=7.4 Hz), 7.00 (CH, ovlp), 7.28 (CH, d, *J*=8.1 Hz), 7.03 (CH, ovlp), 6.04 (NH, m), 10.66 (NH, s); Lys^2^: 4.01 (CH, m), 1.55 (CH_2_, m), 1.28, 1.15 (CH_2_, m), 1.42 (CH_2_, m), 3.53, 2.77 (CH_2_, m), 6.50 (NH, m); Phe^3^: 4.36 (CH, m), 3.26, 2.74 (CH_2_, m), 7.05 (2CH, d, *J*=7.2 Hz), 7.20 (2CH, t, *J*=7.3 Hz), 7.16 (CH, d, *J*=7.2 Hz), 8.53 (NH, d, *J*=8.9 Hz); *N*-Me-Ala^4^: 4.77 (CH, m), 1.06 (CH_3_, d, *J*=6.6 Hz), 1.76 (*N*CH_3_, s); Htyr^5^: 4.75 (CH, m), 1.88, 1.73 (CH_2_, m); 2.63, 2.43 (CH_2_, m), 7.00 (2CH, d, *J*=8.5 Hz), 6.68 (2CH, d, *J*=8.0 Hz), 9.01 (NH, d, *J*=5.0 Hz); Met^6^: 4.20 (CH, m), 1.99 (CH_2_, m), 2.62 (CH_2_, m), 2.08 (CH_3_, s); ^13^C NMR (125 MHz, DMSO-*d*_6_) *δ* Trp^1^: 54.5 (CH), 27.5 (CH_2_), 118.5 (CH), 117.5 (CH), 120.0 (CH), 110.6 (CH), 123.0 (CH); Lys^2^: 54.5 (CH), 31.8 (CH_2_), 19.5 (CH_2_), 27.8 (CH_2_), 37.5 (CH_2_); Phe^3^: 54.6 (CH), 37.2 (CH_2_, m), 128.4 (2CH), 128.0 (2CH), 125.8 (CH); *N*-Me-Ala^4^: 54.0 (CH), 13.4 (CH_3_), 26.5 (NCH_3_); Htyr^5^: 48.2 (CH), 32.9 (CH_2_), 28.9 (CH_2_), 128.8 (2CH), 114.7 (2CH); Met^6^: 51.8 (CH), 31.1 (CH_2_), 28.9 (CH_2_), 14.2 (CH_3_). HRMS (ESI-QTOF) *m/z* calcd for C_46_H_59_N_8_O_9_S^+^: 899.4120 [M+H]^+^; found 899.4128 (error 0.89 ppm).

### Biological assays

Neutrophil elastase inhibition was evaluated using a Human Neutrophil Elastase Inhibitor Screening Kit (Abcam, Cambridge, UK) following the manufacturer’s protocol. Compounds **1** and **2** were tested over a serial dilution range from 10 μM to 1 nM in DMSO, with micropeptin 996 included as a positive control. Fluorescence was monitored at λ_ex/em_ = 400/505 nm at 10 min intervals for 30 min using a Tecan Spark multimode plate reader. IC_50_ values were calculated using Prism 9.5.1 (GraphPad, San Diego, CA, USA).

### Collection of cyanobacterial field samples, toxin analysis, and genomic analysis

Surface water samples were collected from five sites along western Lake Erie on July 23, 2025 (1. Huntington Beach Park, Bay Village, OH, USA, 41.48990°N, 81.93071°W; 2. Miller Road Park, Avon Lake, OH, USA; 41.50275°N, 82.06157°W; 3. Showse Park, Vermilion, OH, USA, 41.43014°N, 82.31420°W; 4. Nickel Plate Beach, Huron, OH, USA, 41.39612°N, 82.54391°W; 5. Huron Harbor North, Huron, OH, USA, 41.39154°N, 82.55421°W), and two samples from East Sandusky Bay (41.45929°N, 82.70725°W) on August 11, 2025, during a cyanoHAB event. At each site, approximately 1 L of surface water was collected into opaque amber HDPE bottles and transported to the laboratory on ice. Upon arrival, 60 mL aliquots of each sample were filtered through 47 mm, 0.2 μm Sterlitech PETE membrane filters. Four replicate filters were prepared per site: two replicates stored at –20°C for cyanotoxin analysis, and the remaining two replicates stored at – 80°C for DNA extraction and molecular analyses.

Laboratory cultures were established by inoculating 1 mL of field water into 9 mL of BG-11 medium, with two cultures per site (n = 14 total). Cultures were maintained at 22°C under a 12 h light/12 h dark photoperiod for three weeks. Biomass was collected by filtration through 47 mm, 0.2 μm membrane filters. Metabolites from both field samples and laboratory cultures were extracted directly from the filters by three sequential extractions with 2 mL of 75% CH_3_OH each. The combined extracts were concentrated under reduced pressure and reconstituted in 100 μL of CH_3_OH. Samples were then loaded onto preconditioned 100 mg C18 SPE cartridges and eluted with CH_3_OH. The final eluates were collected in LC-MS vials at a final volume of 1 mL before LC-MS/MS analysis. LC-MS/MS analysis was carried out on the Agilent Revident QTOF as described above with identical molecular networking workflows at GNPS2.

Genomic DNA was extracted from a 10 mL laboratory culture, which showed the presence of micropeptins using the Qiagen DNeasy Plant Mini Kit with minor modifications. Cells were pelleted by centrifugation at 4000 × g for 10 min, resuspended in 400 μL of AP1 buffer with 4 μL RNase A, and disrupted by bead beating for 1 min. Following lysis, DNA was purified and eluted in 200 μL of Buffer AE according to the manufacturer’s protocol. Concentration and purity were assessed using a NanoDrop 2000 spectrophotometer and a Qubit fluorometer prior to sequencing. Purified DNA was sequenced by Plasmidsaurus using the Oxford Nanopore Technologies platform.

## RESULTS AND DISCUSSION

### LC-MS/MS-based molecular networking: Mass QL and product ion analysis for cyanopeptide annotation and MS/MS characterization of micropeptins

Tandem mass spectrometry-based molecular networking identified 1,063 molecular features in the initial group of fractions from the Muskegon, MI cyanoHAB chromatography fraction library. We interrogated the network using product ion queries that have been reported for different cyanopeptide groups.^15-17^ Intriguingly, this approach was not only able to clearly identify cyanopeptide classes but could also distinguish analog groups within classes in certain cases (Figure 1A-D). Using a Mass QL search of product ions for proposed fragments of [Ahp-Phe-*N*-Me-Phe+H–H_2_O]^+^ (*m/z* 404), [Ahp-Phe+H–H_2_O]^+^ (*m/z* 243), and the *N*-methyl phenylalanine immonium ion *(m/z* 134), we were able to highlight a cluster of molecules in the full network (Figure 1B). Analyzing the MS/MS fragmentation patterns and retention times of these analytes, identified *m/z* 979 [M+H–H_2_O]^+^ as micropeptin 996 (L-Gln), a previously characterized compound.^18^ We were able to isolate two of the metabolites in this cluster: *m/z* 993 [M+H–H_2_O]^+^ (**1**) and *m/z* 949 [M+H–H_2_O]^+^ (**2**) and characterize them as new micropeptins based on MS/MS and NMR data (described below) (Figure 1C).

**Figure 1.**
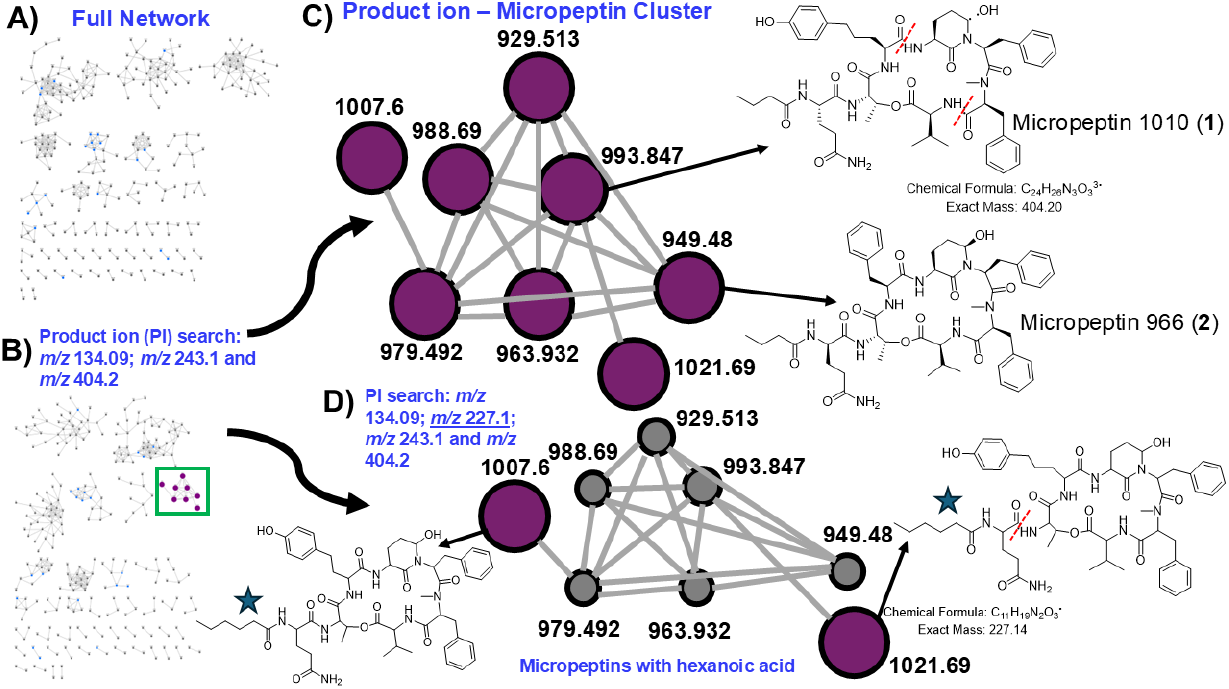
(A) Molecular network derived from analysis of the chromatography fraction library. (B) Product ion search (*m/z* 134.09, *m/z* 243.1, *m/z* 404.2), which highlighted a cluster of micropeptins in the network (green box). (C) Enhanced view of the micropeptin cluster with new structures shown at right (micropeptin 1010 (**1**) and micropeptin 966 (**2**). Red dashed lines show the fragmentation that results in *m/z* 404 in **1**. (D) A second product ion search adding in *m/z* 227.1, which annotated two putative new micropeptins (node 1021.69: micropeptin 1038 and node 1007.6: micropeptin 1024) with hexanoic acid units (blue stars). The red dashed line shows the fragmentation of micropeptin 1038 (*m/z* 1021.69) key in determining the putative structure. All nodes in the networks are designated by an [M+H– H_2_O]^+^ precursor ion.

The ability to identify and annotate new micropeptins via MS/MS, but difficulty in separation and purification, highlights a limitation of HPLC isolation and NMR-based characterization. We therefore proposed that more structurally related micropeptins co-exist within the fractions yet remain overlooked as evidenced by the number of networks nodes corresponding to analytes that we could not purify and characterize by NMR (Figure 1C), and we conducted thorough HRMS and MS/MS analysis to characterize several new micropeptins (Table S3,S4). During repurification of the second chromatographic fraction from which several micropeptins were isolated including **2**, a minor compound was consistently observed co-eluting with micropeptin 996 (L-Gln). This compound showed ions of *m/z* 973 and *m/z* 933, corresponding to [M+Na]^+^ and [M+H–H_2_O]^+^ ions, respectively. Different methods and stationary phases were used to separate the two compounds, but these attempts were unsuccessful. MS/MS comparison of the two network nodes (*m/z* 933 and *m/z* 979) showed extensive overlapping product ions in a mirror plot (Figure S1). Diagnostic ions at *m/z* 560 [butyric acid (BTA)-Gln-Thr-Val-*N*-MePhe+H]^+^ and *m/z* 404 [*N*-MePhe-Phe-Ahp+H–H_2_O]^+^ established the conserved scaffold and side chain, leaving one unresolved residue at position 5. The presence of methionine was supported by the immonium ion at *m/z* 104, while product ions at *m/z* 535 [Met-Ahp-Phe-*N*-MePhe+H–H_2_O]^+^ and *m/z* 374 [Met-Ahp-Phe+H−H_2_O]^+^ further confirmed the position of Met (Figure S2,S3). The compound was designated micropeptin 950, with the proposed sequence BTA-Gln-Thr-Met-Ahp-Phe-*N*MePhe-Val. Delving further into the micropeptin network (Figure 1C), two compounds were rapidly characterized by MS/MS and named as micropeptin 1005 and micropeptin 980, with proposed sequences BTA-Gln-Thr-Trp-Ahp-Phe-*N*-MePhe-Val and BTA-Gln-Thr-Hphe-Ahp-Phe-*N*-MePhe-Val, respectively following careful product ion annotation (Figure S4,S5). We propose Hphe due to the dearth of *N*-methylated residues in micropeptins at position 5 and the common presence of homologated amino acid residues. Another node, characterized by a dehydrated ion at *m/z* 929.5, was named as micropeptin 946 and assigned the sequence BTA-Gln-Thr-Hleu-Ahp-Phe-*N*-MePhe-Val or BTA-Gln-Thr-Hile-Ahp-Phe-*N*-MePhe-Val, as MS/MS fragmentation could not distinguish between leucine and isoleucine (Figure S6). Structural elucidation followed the same workflow applied to micropeptin 950: conserved fragments at *m/z* 560 and *m/z* 404 defined most of the residues in the cyclic scaffold and the side chain, while the residue at position 5 was resolved using HRMS data. Consistent fragmentation patterns across all annotated micropeptins, including ions corresponding to [BTA-Gln-Thr-X+H–H_2_O]^+^ and [X-Ahp-Phe-*N*-MePhe+H–H_2_O]^+^, further supported these assignments. Additional product ion searching incorporating *m/z* 227 annotated two molecules in the cluster as putative new micropeptins each proposed to contain a hexanoic acid (HA) starting unit based on key product ions at *m/z* 227 [HA-Gln]^+^ and *m/z* 310 [HA-Gln-Thr+H–H_2_O]^+^ (Figure 1D) (28 Da higher mass shifts than *m/z* 199 and *m/z* 282, respectively), which we assigned the names micropeptin 1038 and micropeptin 1024 (Figure S7,S8).

We next searched for product ions associated with microcystins (*m/z* 105.05, *m/z* 135.08, and *m/z* 213.11).^15^ This highlighted a small cluster in the full network (Figure 2A). We isolated the analyte with *m/z* 1037 and characterized it via HRMS (found, 1037.6030; calculated, 1037.6030 for C_52_H_81_N_10_O_12_^+^) and ^1^H NMR as [Leu^1^]MC-LR (Figure S9,S10) (Figure 2B),^19^ and the MS/MS pattern comparisons between *m/z* 1037 and *m/z* 1051 strongly suggested that the *m/z* 1051 compound was [Leu^1^,Glu(OCH_3_)^6^]MC-LR (Figure 2C).^15^

**Figure 2.**
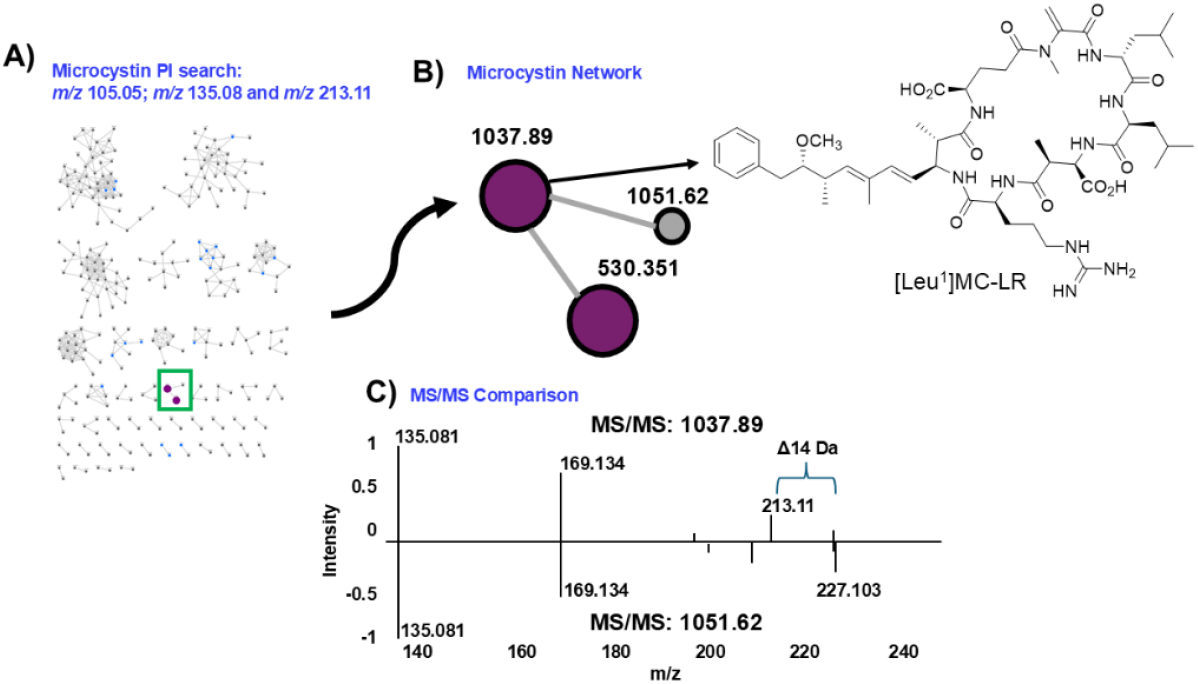
(A) Molecular network of the chromatography fraction library with product ion search for microcystins (*m/z* 105.05, *m/z* 135.08, and *m/z* 213.11). (B) Microcystin cluster with *m/z* 1037.89, which was later identified as [Leu^1^]MC-LR. (C) Comparison of the MS/MS fragmentation patterns of *m/z* 1037.89 and *m/z* 1051.62 in a mirror plot showing a difference of 14 mass units and a *m/z* 227.103 fragment in *m/z* 1051.62 consistent with the fragmentation pattern of [Leu^1^,Glu(OCH_3_)^6^]MC-LR.^15^

### Structure characterization of 1-3

Following the MS/MS network annotation, we identified high abundance ions *m/z* 1033 [M+Na]^+^ (**1**) and *m/z* 989 [M+Na]^+^ (**2**) as putative new micropeptins and targets for isolation and structure elucidation. Compound **1** had a precursor *m/z* of 1033.5009 [M+Na]^+^ suggesting a molecular formula of C_53_H_70_N_8_O_12_. A thorough analysis of product ions supported fragments for Ahp-Phe-*N*-Me-Phe–H_2_O (*m/z* 404), Ahp-Phe–H_2_O (*m/z* 243) and *m/z* 282 likely corresponding to a BTA-Gln-Thr–H_2_O fragment (Figure S11), and these fragment data were in harmony with the 1D and 2D NMR data (Figure S12-S17). This left two amino acid positions unassigned, which NMR was able to unequivocally address. A ^1^H-^1^H TOCSY spin system *δ*_H_ 4.73 (CH), 2.07 (CH), and two methyl signals at *δ*_H_ 0.87 and 0.73 strongly supported a valine residue which NOE correlations positioned adjacent to the *N*-methyl-Phe. The final amino acid was assigned as bis-homologated tyrosine (bHtyr) based on the TOCSY correlations between three CH_2_ signals in the side chain of a modified Tyr residue (*δ*_H_ 1.83, 1.41 and 2.40 and 2.35) (Table S1). NOE correlations positioned this residue between the Ahp residue and the threonine. The bHtyr residue was also supported by product ions of *m/z* 595 (bHtyr-Ahp-Phe-*N*-Me-Phe– H_2_O) and *m/z* 434 (bHtyr-Ahp-Phe-*N*-Me-Phe–H_2_O) (Figure S11). This is the first time the bHtyr has been reported in a micropeptin and we named this molecule micropeptin 1010. Compound **2** gave a high-resolution precursor *m/z* value of 989.4737 [M+Na]^+^ supporting a formula of C_51_H_66_N_8_O_11_. NMR analysis assigned a structure nearly identical to **1**, but with a Phe residue in place of the bHtyr residue (cf. Table S2 and Figures S18-S21). Again, this structure was also supported by a product ion at *m/z* 551 in the MS/MS spectrum (Phe-Ahp-Phe-*N*-Me-Phe–H_2_O) (Figure S22). We named this compound micropeptin 966. A third molecule was isolated, although it was not present in the first molecular network (**3**). The precursor *m/z* of 899.4128 supported a molecular formula of C_46_H_59_N_8_O_9_S (Figure S23) and database searching identified ferintoic acid C as a possible identity. This molecule has been characterized via MS/MS.^12^ The 1D and 2D NMR data were fully supportive of the previously proposed structure including the methionine residue as supported by the correlations observed in the multiplicity-edited HSQC spectrum for *δ*_H_ 2.08 (CH_3_) and *δ*_C_ 14.2 (CH_3_) (Figure S24-S27). The amino acid configurations of **1** were all of the L-configuration as determined by Marfey’s analysis (Table S5, Figure S28), while micropeptin 966 (**2**) possessed a D-glutamine (Figure S29) with the remaining amino acids in the L-configuration (Table S5). Ferintoic acid C (**3**) had a D-lysine, which is the convention in this group of molecules, with the remaining amino acids in the L-configuration (Table S5).

### Protease inhibition activity

Micropeptins are well-established inhibitors of serine proteases, and our previous work showed that many micropeptins show potent activity against human neutrophil elastase.^5^ To continue this structure-activity relationship, compounds **1** and **2** were evaluated for their inhibitory activity against human neutrophil elastase with micropeptin 996 (L-Gln) used as a positive control, which showed an IC_50_ of 0.52 µM. Micropeptin 1010 (**1**) showed moderate inhibitory activity, with an IC_50_ of 3.4 µM (Figure S30). In contrast, **2** was not active at concentrations up to 10 µM. Micropeptins 996 (L-Gln), **1**, and **2** are closely related analogues that differ only at position 5, which is occupied by homologated tyrosine, bishomologated tyrosine, and phenylalanine, respectively. The observed differences in inhibitory activity suggest that the presence of a phenolic hydroxyl group at this position plays a critical role in protease inhibition, likely by stabilizing key interactions within the protease active site, and its absence results in markedly reduced inhibition.

### Additional metabolite annotation workflows of cyanopeptides using product ion searching

After the initial chromatography fraction molecular networking and micropeptin and microcystin annotation, we analyzed a larger chromatography fraction data set of 99 samples derived from the same Muskegon, MI cyanoHAB material to access more chemical space and additional cyanopeptide groups. We also had a standard library of cyanopeptides from our previous work consisting of microcystins, micropeptins/cyanopeptolins, anabaenopeptins/ferintoic acids, microginins, and microviridins for authentication of network hits.^5,14^ The new network had 5,998 features with over 70 nodes in a putative micropeptin/cyanopeptolin network (Figure 3A). Product ion searching via Mass QL could decipher two subgroups in the micropeptin/cyanopeptolin network. Product ion searching using *m/z* 209.128 and *m/z* 370.21 showed a group of molecules containing either leucine or isoleucine,^17^ while searching for *m/z* 243.114 and *m/z* 404.198 showed a group with phenylalanine (Figure 3B). We were also able to designate different groups of microcystins one group with [Glu(OCH_3_)^6^] modifications (product ions: *m/z* 135.08 and *m/z* 389.2),^15,16^ validated by previously characterized microcystins in our own laboratory standard library (Figure S31).^14^ An anabaenopeptin/ferintoic acid cluster was highlighted by searching product ions *m/z* 114.055 and *m/z* 405.2,12 which included ferintoic acids A and C (**3**) (Figure S32).^13^ We searched the network for analytes with product ions of *m/z* 116.07 and *m/z* 159.09 to annotate a cluster of microviridins (Figure S33),^20^ and product ions of *m/z* 147, *m/z* 207, *m/z* 221, *m/z* 281, *m/z* 342, *m/z* 403 to annotate microginins (Figure S34), again annotating specific library molecules in these clusters.

**Figure 3.**
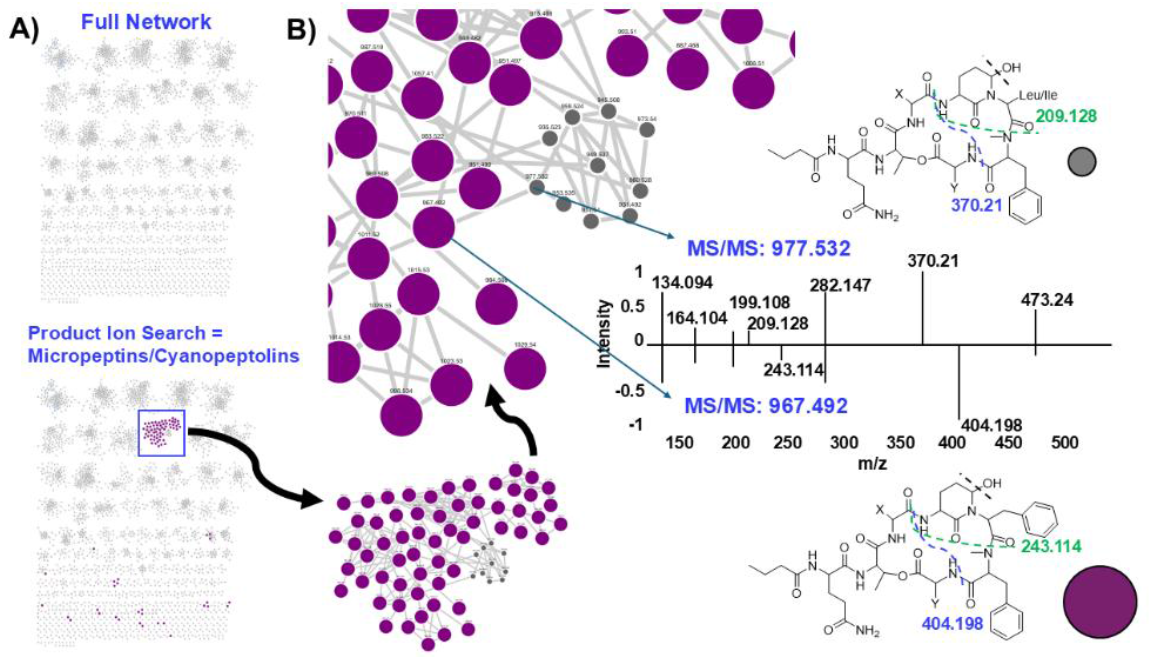
A) Full network of analytes from the full Muskegon, MI cyanoHAB chromatography library (top network) with product ion searching for cyanopeptolins/micropeptins in the bottom network. B) Enhanced view of the cyanopeptolin/micropeptin network showing cluster members with product ion *m/z* 243 and *m/z* 404 (purple). The remaining small gray nodes had a different product ion profile (e.g., *m/z* 209 and *m/z* 370) supporting the presence of a leucine or isoleucine residue versus the phenylalanine residue present in molecules with *m/z* 243 and *m/z* 404, illustrated with the mirror plot of MS/MS spectra and putative fragmentations shown with dashed lines.

### Cyanopeptides in cultures established from Lake Erie and analysis of a metagenome

We screened both initial field collections and starter cultures collected from several Lake Erie sites using two pipelines. The first examined HRMS MS1 data against the CyanoMetDB^21^ using the compound identification function available in Agilent’s Mass Explorer software. We assigned a putative ID to analytes showing a scoring mark of 80 or greater and a mass accuracy within 5 ppm. There were 13 putative compound identifications that met these criteria: carmaphycins A & B, [D-Asp]MC-RF, fisherellin B, lyngbyapeptin D, malyngamide H microcolins A & F, micropeptins 996 and 982, neo-debromoaplysiatoxin H, nostocyclyne A, ribocyclophane D. The second pipeline utilized the LC-MS/MS workflow with product ion searching and comparison of MS/MS data to literature and the GNPS library. We were able to identify both micropeptin 982 and micropeptin 996 in the molecular network created from the field samples (Figure 4A), and as these standards were both in our library,^22^ we were also able to use these authentic standards for verification (Figure 4B), show that they were in all field samples (Figure 4C), and validate them using the standards that we have accumulated over the years from previous research (Table S6, Figure 4D).^22^ However, we could not unequivocally annotate any other cyanopeptides in the field samples that satisfied MS/MS fragmentation data validation other than these two micropeptins. Metagenomic sequencing and analysis of the sample from Miller Road Park identified a 2215 bp section that BLAST searching matched to the *ociB* gene in cyanopeptolin biosynthesis from *Plantothrix agardhii* (Figure S35),^23^ supporting a likely connection between the chemistry found from the field sample and the genetic architecture that produces cyanopeptolins/micropeptins.

**Figure 4.**
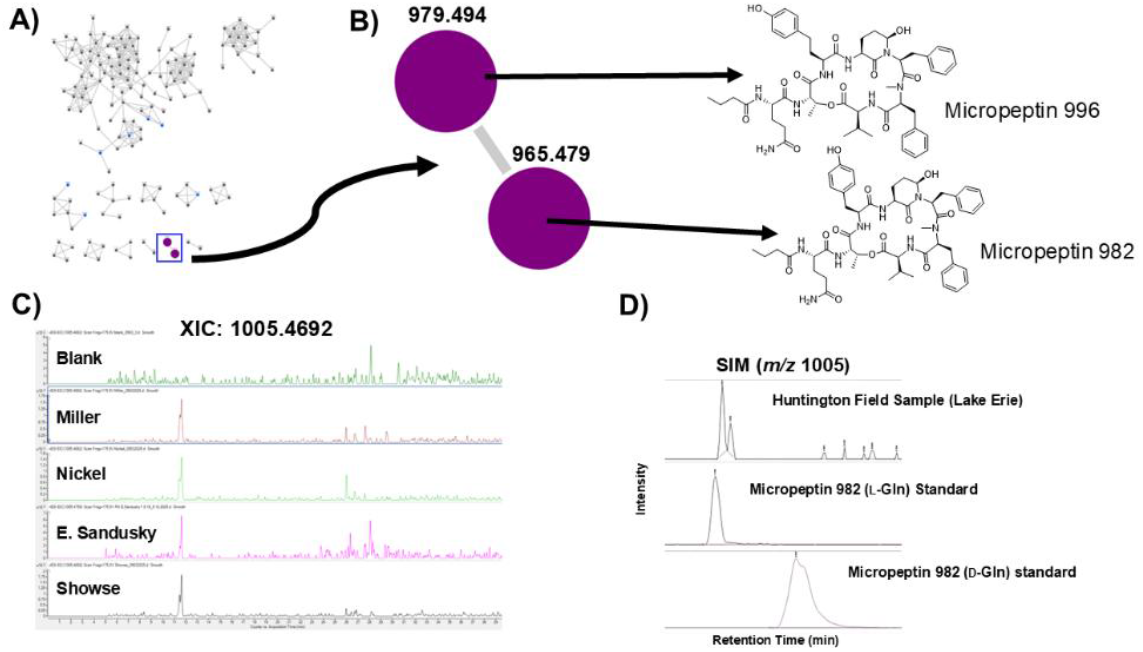
A) LC-MS/MS-based molecular network of field samples from Lake Erie samples with micropeptin cluster highlighted in the blue box. B) Network cluster showing micropeptin nodes [M+H–H_2_O]^+^ and identified micropeptin compounds in our compound library (micropeptin 996 and micropeptin 982). C) Extracted ion chromatograms of selected field samples for micropeptin 982 (*m/z* 1005.4692, [M+Na]^+^). D) Comparison of Huntington field sample with two authentic standards of micropeptin 982 (L-Gln) and micropeptin 982 (D-Gln) monitoring *m/z* 1005.

These results indicate that Great Lakes cyanoHABs and cyanoHABs in other water bodies (e.g., Muskegon, MI) contain substantial, recurrent non-microcystin cyanopeptide chemistry that is largely invisible to routine targeted monitoring. Product ion–driven GNPS2 annotation provides a scalable path to class-level screening of these metabolites in complex bloom samples, enabling mixture assessment, and prioritization for toxicological and treatment studies.

Together, the chemistry, bioactivity, and genetic evidence we provide in this report support expanded surveillance beyond canonical toxins to better represent real-world bloom exposure profiles. Cyanopeptolins/micropeptins have shown aquatic toxicity,^24^ but their environmental impact and toxicological profiles are unknown and our SAR approach and large library generation can prioritize metabolites for further *in vitro* and *in vivo* studies. The molecular networks showed multiple cyanopeptide classes (micropeptins, microcystins, anabaenopeptins/ferintoic acids, microginins, microviridins) and dozens of individual metabolites in each cluster reinforcing that blooms are chemical mixtures, and the ecological/toxic effects from these events may reflect additivity/synergy/antagonism rather than one compound-one effect paradigms (e.g., microcystin-LR). This can be supported by mixture-aware monitoring and bioassays (effect-directed analysis). While there are thousands of nodes in the network, it is important to note that the network does overrepresent analyte space due to in-source fragmentation and adduct profiles.^25^ Some of these issues can be abrogated with additional post acquisition workflows.^26^ While molecular networking has been used previously in cyanopeptide annotation,^12,22,27^ most of these studies focus on class-level identification whereas we use the Mass QL tool to probe within classes for unprecedented levels of annotation. The GNPS2-based product ion annotation will also be valuable to resource managers and public health officials as it is broadly generalizable across mass spectrometry platforms and reduces the need for standards for every congener providing both untargeted data for continued mining, but also an element of targeted analysis with respect to cyanometabolite classes and annotation within classes. While the product ion search platform provides excellent large dataset annotation (class assignment and subfamily discrimination), there are clear limitations as to what product ion searching cannot provide (isomer resolution, stereochemistry, absolute quantitation). We abrogated these limitations by isolating and fully characterizing new molecules including, identifying for the first time, the incorporation of bHtyr in the micropeptins. Our standard library provided retention time isomer matching and we also provide orthogonal metagenomic support for cyanopeptolin/micropeptin biosynthesis. We only sampled over a short temporal time scale before the biomass maximum of the Lake Erie bloom,^28^ and thus we may not have annotated microcystins and other toxins in bloom samples, as we have shown previously that there is a correlation between biomass maxima and microcystin production.^14^ Enhancing spatial and temporal resolution of bloom monitoring can provide a richer data set and ultimately illustrate that monitoring programs may need site-specific and time-specific monitoring panels or adaptable suspect screening rather than a fixed toxin list. Finally, we were focused on qualitative annotation and not quantitative assessment in this study, but quantitation can be easily added to untargeted metabolomics workflows with internal standards and calibration curves or even ion count filtering in MS/MS networks. Furthermore, additional computational workflows can integrate peptide sequencing machine learning platforms, additional molecular feature finding, and genomics and transcriptomics pipelines to identify more cyanometabolites and connect taxa, genetic architecture, and chemistry to public health and environmental health outcomes annotating unprecedented levels of bloom chemical space.

## ASSOCIATED CONTENT

Supplementary data can be found with the online version of this article. It includes: Additional experimental details, NMR data, mass spectrometry data, bioassay data, and genomic data. (PDF)

NMR data for compounds **1-3** (NP0352196, NP0352197, NP0352198) and [Leu1]MC-LR (NP0352199) have been deposited at https://np-mrd.org/ and tandem mass spectrometry data have been deposited at massIVE under accession number MSV000100694.

## AUTHOR INFORMATION

**Runjie Xia**, Department of Chemistry, Case Western Reserve University, Cleveland, OH 44106, United States

**Lindsey Ahn**, Department of Chemistry, Case Western Reserve University, Cleveland, OH 44106, United States

**Michaela Burkhauser**, Department of Chemistry, Case Western Reserve University, Cleveland, OH 44106, United States

**Ross Youngs**, Biosortia, Inc., 2545 Farmers Dr., Suite 370, Columbus, OH 43235, United States.

## Author Contributions

M. J. B. conceived the study and performed analysis. R. X., L. A., and M. B. performed methodology, data acquisition, and analysis. R. Y. provided key materials and manuscript reviewing. The manuscript was written through contributions of all authors. All authors have given approval to the final version of the manuscript.

## Notes

R. Y. is the founder and CEO and has a financial interest in Biosortia Inc., which provided samples for this study. R.Y.’s role was limited to sample provision and manuscript review. All other authors declare no competing interests.

## ACKNOWLEDGMENTS

Research reported in this article was supported by the National Institute of Environmental Health Sciences of the National Institutes of Health under award number R21ES033758 (M. J. B.). The content is solely the responsibility of the authors and does not necessarily represent the official views of the National Institutes of Health. Additionally, L. A. was supported by a CWRU SOURCE Summer Fellowship, and M. B. was supported by an Expanding Horizons Initiative Award from CWRU.

## REFERENCES

1) Wang, Y.; Zhao, D.; Woolway, R. I.; Yan, H.; Paerl, H. W.; Zheng, Y.; Zheng, C.; Feng, L. Global Elevation of Algal Bloom Frequency in Large Lakes over the Past Two Decades. National Science Review 2025, 12 (3), nwaf011.

2) Merder, J.; Harris, T.; Zhao, G.; Stasinopoulos, D. M.; Rigby, R. A.; Michalak, A. M. Geographic Redistribution of Microcystin Hotspots in Response to Climate Warming. Nat. Water 2023, 1 (10), 844–854.

3) Environmental Protection Agency. Drinking Water Health Advisory for the Cyanobacterial Microcytins Toxins. 2015;EPA, Office of Water. EPA document number:820R15100.

4) Chorus, I.; Welker, M., Eds. Toxic Cyanobacteria in Water: A Guide to Their Public Health Consequences, Monitoring and Management, 2nd ed.; CRC Press: London, 2021.

5) Xia, R.; Xie, H.; Mahmud, A. M. S.; Bertin, M. J. Alteration in Amino Acid Composition and Configuration in Cyanobacterial Peptides Affects Biological Activity. J. Nat. Prod. 2025, 88 (8), 1950–1957.

6) Yancey, C. E.; Hart, L.; Hefferan, S.; Mohamed, O. G.; Newmister, S. A.; Tripathi, A.; Sherman, D. H.; Dick, G. J. Metabologenomics Reveals Strain-Level Genetic and Chemical Diversity of Microcystis Secondary Metabolism. mSystems 2024, 9 (7), e00334–24.

7) Stringer, B. B.; Szlag Silva, R. G.; Kodanko, J. J.; Westrick, J. A. Structure, Toxicity, Prevalence, and Degradation of Six Understudied Freshwater Cyanopeptides. Toxins 2025, 17 (5), 233.

8) Janssen, E. M.-L. Cyanobacterial Peptides beyond Microcystins – A Review on Co-Occurrence, Toxicity, and Challenges for Risk Assessment. Water Research 2019, 151, 488–499.

9) He, H.; Wahome, P. G.; Bertin, M. J.; Pedone, A. C.; Beauchesne, K. R.; Moeller, P. D. R.; Carter, G. T. Microcystins Containing Doubly Homologated Tyrosine Residues from a Microcystis Aeruginosa Bloom: Structures and Cytotoxicity. J. Nat. Prod. 2018, 81, 1368–1375.

10) Wang, M.; Carver, J. J.; Phelan, V. V.; Sanchez, L. M.; Garg, N.; Peng, Y.; Nguyen, D. D.; Watrous, J.; Kapono, C. A.; Luzzatto-Knaan, T.; Porto, C.; Bouslimani, A.; Melnik, A. V.; Meehan, M. J.; Liu, W.-T.; Crüsemann, M.; Boudreau, P. D.; Esquenazi, E.; Sandoval-Calderón, M.; Kersten, R. D.; Pace, L. A.; Quinn, R. A.; Duncan, K. R.; Hsu, C.-C.; Floros, D. J.; Gavilan, R. G.; Kleigrewe, K.; Northen, T.; Dutton, R. J.; Parrot, D.; Carlson, E. E.; Aigle, B.; Michelsen, C. F.; Jelsbak, L.; Sohlenkamp, C.; Pevzner, P.; Edlund, A.; McLean, J.; Piel, J.; Murphy, B. T.; Gerwick, L.; Liaw, C.-C.; Yang, Y.-L.; Humpf, H.-U.; Maansson, M.; Keyzers, R. A.; Sims, A. C.; Johnson, A. R.; Sidebottom, A. M.; Sedio, B. E.; Klitgaard, A.; Larson, C. B.; Boya P C.A.;, Torres-Mendoza, D.; Gonzalez, D. J.; Silva, D. B.; Marques, L. M.; Demarque, D. P.; Pociute, E.; O’Neill, E. C.; Briand, E.; Helfrich, E. J. N.; Granatosky, E. A.; Glukhov, E.; Ryffel, F.; Houson, H.; Mohimani, H.; Kharbush, J. J.; Zeng, Y.; Vorholt, J. A.; Kurita, K. L.; Charusanti, P.; McPhail, K. L.; Nielsen, K. F.; Vuong, L.; Elfeki, M.; Traxler, M. F.; Engene, N.; Koyama, N.; Vining, O. B.; Baric, R.; Silva, R. R.; Mascuch, S. J.; Tomasi, S.; Jenkins, S.; Macherla, V.; Hoffman, T.; Agarwal, V.; Williams, P. G.; Dai, J.; Neupane, R.; Gurr, J.; Rodríguez, A. M. C.; Lamsa, A.; Zhang, C.; Dorrestein, K.; Duggan, B. M.; Almaliti, J.; Allard, P.-M.; Phapale, P.; Nothias, L.-F.; Alexandrov, T.; Litaudon, M.; Wolfender, J.-L.; Kyle, J. E.; Metz, T. O.; Peryea, T.; Nguyen, D.-T.; VanLeer, D.; Shinn, P.; Jadhav, A.; Müller, R.; Waters, K. M.; Shi, W.; Liu, X.; Zhang, L.; Knight, R.; Jensen, P. R.; Palsson, B.Ø.; Pogliano, K.; Linington, R. G.; Gutiérrez, M.; Lopes, N. P.; Gerwick, W. H.; Moore, B. S.; Dorrestein, P. C.; Bandeira, N. Sharing and Community Curation of Mass Spectrometry Data with Global Natural Products Social Molecular Networking. Nat. Biotechnol. 2016, 34 (8), 828–837.

11) Petras, D.; Phelan, V. V.; Acharya, D.; Allen, A. E.; Aron, A. T.; Bandeira, N.; Bowen, B. P.; Belle-Oudry, D.; Boecker, S.; Cummings, D. A.; Deutsch, J. M.; Fahy, E.; Garg, N.; Gregor, R.; Handelsman, J.; Navarro-Hoyos, M.; Jarmusch, A. K.; Jarmusch, S. A.; Louie, K.; Maloney, K. N.; Marty, M. T.; Meijler, M. M.; Mizrahi, I.; Neve, R. L.; Northen, T. R.; Molina-Santiago, C.; Panitchpakdi, M.; Pullman, B.; Puri, A. W.; Schmid, R.; Subramaniam, S.; Thukral, M.; Vasquez-Castro, F.; Dorrestein, P. C.; Wang, M. GNPS Dashboard: Collaborative Exploration of Mass Spectrometry Data in the Web Browser. Nat. Methods 2022, 19 (2), 134–136.

12) Damiani, T.; Jarmusch, A. K.; Aron, A. T.; Petras, D.; Phelan, V. V.; Zhao, H. N.; Bittremieux, W.; Acharya, D. D.; Ahmed, M. M. A.; Bauermeister, A.; Bertin, M. J.; Boudreau, P. D.; Borges, R. M.; Bowen, B. P.; Brown, C. J.; Chagas, F. O.; Clevenger, K. D.; Correia, M. S. P.; Crandall, W. J.; Crüsemann, M.; Fahy, E.; Fiehn, O.; Garg, N.; Gerwick, W. H.; Gilbert, J. R.; Globisch, D.; Gomes, P. W. P.; Heuckeroth, S.; James, C. A.; Jarmusch, S. A.; Kakhkhorov, S. A.; Kang, K. B.; Kessler, N.; Kersten, R. D.; Kim, H.; Kirk, R. D.; Kohlbacher, O.; Kontou, E. E.; Liu, K.; Lizama-Chamu, I.; Luu, G. T.; Luzzatto Knaan, T.; Mannochio-Russo, H.; Marty, M. T.; Matsuzawa, Y.; McAvoy, A. C.; McCall, L.-I.; Mohamed, O. G.; Nahor, O.; Neuweger, H.; Niedermeyer, T. H. J.; Nishida, K.; Northen, T. R.; Overdahl, K. E.; Rainer, J.; Reher, R.; Rodriguez, E.; Sachsenberg, T. T.; Sanchez, L. M.; Schmid, R.; Stevens, C.; Subramaniam, S.; Tian, Z.; Tripathi, A.; Tsugawa, H.; van der Hooft, J. J. J.; Vicini, A.; Walter, A.; Weber, T.; Xiong, Q.; Xu, T.; Pluskal, T.; Dorrestein, P. C.; Wang, M. A Universal Language for Finding Mass Spectrometry Data Patterns. Nat. Methods 2025, 22 (6), 1247–1254.

13) Teta, R.; Della Sala, G.; Glukhov, E.; Gerwick, L.; Gerwick, W. H.; Mangoni, A.; Costantino, V. Combined LC–MS/MS and Molecular Networking Approach Reveals New Cyanotoxins from the 2014 Cyanobacterial Bloom in Green Lake, Seattle. Environ. Sci. Technol. 2015, 49 (24), 14301–14310.

14) Maurer, J. A.; Xia, R.; Kim, A. M.; Oblie, N.; Hefferan, S.; Xie, H.; Slitt, A.; Jenkins, B. D.; Bertin, M. J. Temporal Dynamics of Cyanobacterial Bloom Community Composition and Toxin Production from Urban Lakes. ACS ES&T Water 2024, 4(8), 3423-3432.

15) Cottrill, K. A.; Miles, C. O.; Krajewski, L. C.; Cunningham, B. R.; Bragg, W.; Boise, N. R.; Victry, K. D.; Wunschel, D. S.; Wahl, K. L.; Hamelin, E. I. Identification of Novel Microcystins in Algal Extracts by a Liquid Chromatography–High-Resolution Mass Spectrometry Data Analysis Pipeline. Harmful Algae 2024, 139, 102739.

16) Qi, Y.; Rosso, L.; Sedan, D.; Giannuzzi, L.; Andrinolo, D.; Volmer, D. A. Seven New Microcystin Variants Discovered from a Native Microcystis Aeruginosa Strain – Unambiguous Assignment of Product Ions by Tandem Mass Spectrometry. Rapid Comm. Mass Spectrometry 2015, 29 (2), 220–224.

17) McDonald, K.; Renaud, J. B.; Pick, F. R.; Miller, J. D.; Sumarah, M. W.; McMullin, D. R. Diagnostic Fragmentation Filtering for Cyanopeptolin Detection. Environ. Toxicol. Chem. 2020, 40 (4), 1087–1097.

18) Strangman, W. K.; Stewart, A. K.; Herring, M. C.; Wright, J. L. C. Identification of the new chymotrypsin inhibitor micropeptin 996 by metabolomics-guided analysis. Tetrahedron Lett. 2018, 59, 934–937.

19) Schripsema, J.; Dagnino, D. Complete Assignment of the NMR Spectra of [D -Leu^1^]- microcystin-LR and Analysis of Its Solution Structure. Magn. Reson. Chem. 2002, 40 (9), 614–617.

20) McDonald, K.; DesRochers, N.; Renaud, J. B.; Sumarah, M. W.; McMullin, D. R. Metabolomics Reveals Strain-Specific Cyanopeptide Profiles and Their Production Dynamics in Microcystis Aeruginosa and M. flos-aquae. Toxins 2023, 15 (4), 254.

21) Janssen, E. M.-L.; Jones, M. R.; Pinto, E.; Dörr, F.; Torres, M. A.; Rios Jacinavicius, F.; Mazur-Marzec, H.; Szubert, K.; Konkel, R.; Tartaglione, L.; Dell’Aversano, C.; Miglione, A.; McCarron, P.; Beach, D. G.; Miles, C. O.; Fewer, D. P.; Sivonen, K.; Jokela, J.; Wahlsten, M.; Niedermeyer, T. H. J.; Schanbacher, F.; Pedro, L.; Preto, M.; D’Agostino, P. M.; Baunach, M.; Dittmann, E.; Miguel-Gordo, M.; Reher, R.; Sieber, S. S75 | CyanoMetDB | Comprehensive database of secondary metabolites from cyanobacteria (NORMAN-SLE-S75.0.3.0) [Data set]. 2024, Zenodo. 10.5281/zenodo.13854577.

22) Kirk, R. D.; He, H.; Wahome, P. G.; Wu, S.; Carter, G. T.; Bertin, M. J. New Micropeptins with Anti-Neuroinflammatory Activity Isolated from a Cyanobacterial Bloom. ACS Omega 2021, 6 (23), 15472–15478.

23) Rounge, T. B.; Rohrlack, T.; Tooming-Klunderud, A.; Kristensen, T.; Jakobsen, K. S. Comparison of Cyanopeptolin Genes in Planktothrix, Microcystis, and Anabaena Strains: Evidence for Independent Evolution within Each Genus. Appl. Environ. Microbiol. 2007, 73 (22), 7322–7330.

24) de Almeida Torres, M.; Jones, M. R.; vom Berg, C.; Pinto, E.; Janssen, E. M. L. Lethal and Sublethal Effects towards Zebrafish Larvae of Microcystins and Other Cyanopeptides Produced by Cyanobacteria. Aquatic Toxicol. 2023, 263, 106689.

25) Xue, J.; Domingo-Almenara, X.; Guijas, C.; Palermo, A.; Rinschen, M. M.; Isbell, J.; Benton, H. P.; Siuzdak, G. Enhanced In-Source Fragmentation Annotation Enables Novel Data Independent Acquisition and Autonomous METLIN Molecular Identification. Anal. Chem. 2020, 92 (8), 6051–6059.

26) Schmid, R.; Petras, D.; Nothias, L.-F.; Wang, M.; Aron, A. T.; Jagels, A.; Tsugawa, H.; Rainer, J.; Garcia-Aloy, M.; Dührkop, K.; Korf, A.; Pluskal, T.; Kameník, Z.; Jarmusch, A. K.; Caraballo-Rodríguez, A. M.; Weldon, K. C.; Nothias-Esposito, M.; Aksenov, A. A.; Bauermeister, A.; Albarracin Orio, A.; Grundmann, C. O.; Vargas, F.; Koester, I.; Gauglitz, J. M.; Gentry, E. C.; Hövelmann, Y.; Kalinina, S. A.; Pendergraft, M. A.; Panitchpakdi, M.; Tehan, R.; Le Gouellec, A.; Aleti, G.; Mannochio Russo, H.; Arndt, B.; Hübner, F.; Hayen, H.; Zhi, H.; Raffatellu, M.; Prather, K. A.; Aluwihare, L. I.; Böcker, S.; McPhail, K. L.; Humpf, H.-U.; Karst, U.; Dorrestein, P. C. Ion Identity Molecular Networking for Mass Spectrometry-Based Metabolomics in the GNPS Environment. Nat. Commun. 2021, 12 (1), 3832.

27) Kust, A.; Řeháková, K.; Vrba, J.; Maicher, V.; Mareš, J.; Hrouzek, P.; Chiriac, M.-C.; Benedová, Z.; Tesařová, B.; Saurav, K. Insight into Unprecedented Diversity of Cyanopeptides in Eutrophic Ponds Using an MS/MS Networking Approach. Toxins 2020, 12 (9), 561.

28) National Centers for Coastal Ocean Science. 2025 Lake Erie Harmful Algal Bloom Seasonal Assessment. December 4, 2025.

