## Supporting Information for "Expanding CyanoHAB Monitoring: New Micropeptins and Generalizable MS/MS Workflows for the Annotation of Cyanopeptide Classes"

### Additional Experimental Procedures

**General Experimental Procedures.** Optical rotation values were measured using a Jasco P-2000 polarimeter. NMR spectra were recorded on a Bruker 500 MHz Ascend Advance III NMR instrument. Chemical shifts reported for **1-3** were referenced to the residual solvent peak of $(CD_3)_2SO$ ($\delta_H$ 2.50 and $\delta_C$ 39.5). LC-HRMS and LC-HRMS$^2$ data were collected on an Agilent Revident QTOF mass spectrometer equipped with a Jet Stream source and 1290 Infinity II Bio LC (with multisampler and multicolumn thermostat) and MassHunter Workstation software. Additional LC-MS analyses were conducted on an Agilent LC-MSD single-quadrupole mass spectrometer equipped with an Agilent 1260 HPLC system with autosampler. Semipreparative HPLC separations were carried out using an Agilent 1260 Infinity system equipped with a vacuum degasser, autosampler, and diode array detector.

**Configuration Analysis of 1-3.** To determine the absolute configuration of the α-amino acids in compounds **1–3**, 0.4 mg of each compound was hydrolyzed in 0.5 mL of 6 N HCl at 110°C for 16 h. After cooling to room temperature, the hydrolysates were dried under a stream of nitrogen, reconstituted in 100 μL of water and 100 μL of 1 M $NaHCO_3$. Derivatization was performed by the addition of 500 μL of a 1% (w/v) solution of *N*-α-(2,4-dinitro-5-fluorophenyl)-L-valinamide (L-FDVA) in acetone. The reaction mixtures were stirred and heated at 40°C for 1 h, then quenched by the addition of 50 μL of 2 N HCl. The derivatized hydrolysates were subsequently diluted 1:10 with a 1:1 mixture of $H_2O/CH_3CN$ to a final volume of 1 mL. Separations were performed on a Luna C18 column (5 μm, 150 × 2.0 mm) using a linear gradient of water (A) and acetonitrile (B), each modified with 0.1% formic acid, at a flow rate of 0.4 mL/min. The gradient was programmed from 20% to 50% B over 30 min, followed by a return to initial conditions from 31 to 36 min. Absolute configurations were assigned by comparison of retention times with those of L-FDVAderivatized authentic amino acid standards prepared under identical conditions. Using this approach, all proteinogenic amino acid residues were unambiguously assigned. For the bishomologated tyrosine (bHtyr) residue present in compound **1**, an authentic D-bHtyr standard was not available. To resolve its configuration, two aliquots of compound **1** (0.25 mg each) were independently hydrolyzed as described above and derivatized using 1% solutions of *N*-α-(2,4-dinitro-5-fluorophenyl)-L-leucinamide (L-FDLA) and *N*-α-(2,4-dinitro-5-fluorophenyl)-D-leucinamide (D-FDLA) in acetone, respectively. The resulting derivatives were analyzed by LC-MS using the same column, with a modified gradient elution from 20% to 80% B over 30 min, followed by re-equilibration to initial conditions from 31 to 36 min. Configuration was assigned based on the interaction of the derivatives with the reversed-phase column chemistry as has been previously described in cyanopeptide analysis.[1]

### Tables and Figures

**Table S1.** NMR Data for Micropeptin 1010 (**1**).

**Table S2.** NMR data for micropeptin 966 (D-Gln) (**2**).

**Table S3.** Molecular formulas, exact masses, and high-resolution mass spectrometric data for newly identified micropeptins and other cyanopeptides and confidence level of identification.

**Table S4.** Sequences of micropeptin variants isolated/identified in this study (BTA=butanoic acid, HA=hexanoic acid).

**Table S5.** Marfey's derivatization data and assignments for **1–3.**

**Table S6.** Retention times of micropeptin stereoisomers and field sample peaks under LC-MSD conditions.

**Figure S1.** Mirror plot of MS/MS spectra of micropeptin 950 and micropeptin 996.

**Figure S2.** HRMS measurement (*m/z* 973.4456) and annotated MS/MS fragmentation pattern of micropeptin 950.

**Figure S3.** Annotated MS/MS spectrum of micropeptin 950.

**Figure S4.** Annotated MS/MS spectrum of micropeptin 1005.

**Figure S5.** Annotated MS/MS spectrum of micropeptin 980.

**Figure S6.** Annotated MS/MS spectrum of micropeptin 946.

**Figure S7.** Annotated MS/MS spectrum of micropeptin 1038.

**Figure S8.** Annotated MS/MS spectrum of micropeptin 1024.

**Figure S9.** Isolation and characterization of [Leu$^{1}$]MC-LR.

**Figure S10.** $^{1}$H NMR (500 MHz, MeOH-$d_4$) of [Leu$^{1}$]MC-LR.

**Figure S11.** Mass spectrometry data of **1**.

**Figure S12.** $^{1}$H NMR (500 MHz, DMSO-$d_6$) of micropeptin 1010 (**1**).

**Figure S13.** $^{13}$C NMR (125 MHz, DMSO-$d_6$) of micropeptin 1010 (**1**).

**Figure S14.** Multiplicity-edited HSQC of micropeptin 1010 (**1**).

**Figure S15.** HMBC of micropeptin 1010 (**1**).

**Figure S16.** TOCSY of micropeptin 1010 (**1**).

**Figure S17.** NOESY of micropeptin 1010 (**1**).

**Figure S18.** $^{1}$H NMR (500 MHz, DMSO-$d_6$) of micropeptin 966 (D-Gln) (**2**).

**Figure S19.** Multiplicity-edited HSQC of micropeptin 966 (D-Gln) (**2**).

**Figure S20.** TOCSY of micropeptin 966 (D-Gln) (**2**).

**Figure S21.** NOESY of micropeptin 966 (D-Gln) (**2**).

**Figure S22.** Mass spectrometry data of **2.**

**Figure S23.** HRMS of ferintoic acid C *m/z* 899.4128 $[M+H]^{+}$.

**Figure S24.** $^{1}$H NMR (500 MHz, DMSO-$d_6$) of ferintoic acid C (**3**).

**Figure S25.** Multiplicity-edited HSQC of ferintoic acid C (**3**).

**Figure S26.** TOCSY of ferintoic acid C (**3**).

**Figure S27.** NOESY of ferintoic acid C (**3**).

**Figure S28.** LC-MS analysis of the hydrolysate of **1** reacted with L-FDLA (top panel) and a mixture of L- and D-FDLA (bottom panel) to determine the configuration of the bHtyr in **1**.

**Figure S29.** LC-MS analysis of the hydrolysate of **2**.

**Figure S30.** Activity of micropeptin 996 (L-Gln), micropeptin 1010 (**1**), and micropeptin 966 (D-Gln) (**2**) against human neutrophil elastase.

**Figure S31.** Microcystin cluster in MS/MS molecular network subjected to two different product ion searches.

**Figure S32.** MS/MS cluster of anabaenopeptins/ferintoic acids annotated via product ion searching.

**Figure S33.** Microviridin cluster annotated using product ion searching in MS/MS networks.

**Figure S34.** Microginin cluster annotated using product ion searching in MS/MS networks.

**Figure S35.** Blast hit to the *ociB* gene (cyanopeptolin biosynthetic pathway) in the metagenomic sequence data from Lake Erie (Miller Road Park).

**Table S1.** NMR Data for Micropeptin 1010 (**1**) (500 MHz for $^{1}H$ NMR, 125 MHz for $^{13}C$ NMR; DMSO-$d_6$)

| Position | *δ*C, mult | *δ*H, mult, *J* (Hz) | TOCSY | ROESY |
|---|---|---|---|---|
| **Val-1** | | | | |
| 2 | 55.3, CH | 4.73, ovlp | 3, NH | |
| 3 | 30.3, CH | 2.07, m | 2, 4, 5, NH | |
| 4 | 18.9, $CH_3$ | 0.87, ovlp | 3, 5, NH | |
| 5 | 16.8, $CH_3$ | 0.73, d (6.4) | 3, 4, NH | |
| NH | | 7.41, ovlp | 2, 3, 4, 5 | *N*-MePhe-2, *N*-Me |
| ***N*-MePhe-1** | | | | |
| 2 | 60.1, CH | 5.03, ovlp | 3a, 3b | Val-NH, Phe-2 |
| 3a | 33.3, $CH_2$ | 3.23, m | 2 | |
| 3b | | 2.84, m | 2 | |
| 4 | | | | |
| 5/9 | 129.1, CH | 7.25, d (7.3) | 6, 7, 8 | Phe-2 |
| 6/8 | 128.3, CH | 7.40, t (7.3) | 5, 7, 9 | |
| 7 | 126.3, CH | 7.31, t (7.2) | 5, 6, 8, 9 | |
| *N*-Me | 30.0, $CH_3$ | 2.79, s | | Val-NH |
| **Phe-1** | 170.8, C | | | |
| 2 | 49.7, CH | 4.73, ovlp | 3a, 3b | *N*-MePhe-2, 5, 9, Ahp-5 |
| 3a | 34.9, $CH_2$ | 2.83, m | 2, 3b | Ahp-5 |
| 3b | | 1.65, m | 2, 3a | Ahp-5 |
| 4 | | | | |
| 5/9 | 129.1, CH | 6.78, d (7.4) | 6, 7, 8 | Ahp-5 |
| 6/8 | 127.5, CH | 7.18, t (7.3) | 5, 7, 9 | |
| 7 | 125.9, CH | 7.13, d (7.4) | 5, 6, 8, 9 | |
| **Ahp-1** | | | | |
| 2 | 48.2, CH | 3.58, m | 3a, 3b, 4a, 4b, 5, NH | Ahp-2, 3b, 4b |
| 3a | 21.2, $CH_2$ | 2.38, m | 2, 3b, 4a, 4b, 5, NH | Ahp-NH |
| 3b | | 1.56, m | 2, 3a, 4b, NH | Ahp-2 |
| 4a | 28.7, $CH_2$ | 1.66, m | 2, 3a, 4b, 5, NH | Ahp-3a, 5 |
| 4b | | 1.48, m | 2, 3a, 4a, 5, NH | Ahp-2, 5 |
| 5 | 73.2, CH | 5.03, ovlp | 2, 3a, 4a, 4b, OH | Ahp-4a, 4b, Phe-2, 3a, 5, 9 |
| NH | | 7.10, ovlp | 2, 3a, 3b, 4a, 4b | Ahp-3a, bHtyr-2, bHtyr-NH |
| OH | | 6.05, br | 5 | |
| **bHtyr-1** | | | | |
| 2 | 51.3, CH | 4.22, m | 3, 4, 5a, 5b, NH | Ahp-NH |
| 3 | 29.1 $CH_2$ | 1.83, m | 2, 4, 5a, 5b, NH | |
| 4 | 27.1 $CH_2$ | 1.41, m | 2, 3, 5a, 5b, NH | |
| 5a | 33.4, $CH_2$ | 2.40, m | 2, 3, 4, 5b | |
| 5b | | 2.35, m | 2, 3, 4, 5a | |
| 6 | 155.8, C | | | |
| 7/9 | 128.9, CH | 6.91, d (8.2) | 7, 9 | |
| 8/10 | 114.7, CH | 6.64, d (8.2) | 6, 8 | |
| 11 | | | | |
| NH | | 8.42, d (8.7) | 2, 3, 4 | Ahp-NH, Thr-2, 3 |
| **Thr-1** | | | | |
| 2 | 54.2, CH | 4.62, m | NH | bHtyr-NH |
| 3 | 71.6, CH | 5.41, m | 4 | bHtyr-NH |
| 4 | 17.3, $CH_3$ | 1.17, d (6.4) | 3 | |
| NH | | 7.91, d (9.2) | 2 | Gln-2, 3a |
| **Gln-1** | | | | |
| 2 | 51.7, CH | 4.37, m | 3a, 3b, 4, NH | Thr-NH |
| 3a | 27.3, $CH_2$ | 1.90, m | 2, 3b, 4, NH | Thr-NH |
| 3b | | 1.71, m | 2, 3a, 4, NH | |
| 4 | 31.4, $CH_2$ | 2.17, m | 2, 3a, 3b, NH | |

| | | | | |
|---|---|---|---|---|
| 5 | | | | |
| NH | | 8.05, d (7.7) | 2, 3a, 3b, 4 | BTA-2 |
| $NH_2$ | | 7.27, br<br>6.75, br | | |
| **BTA-1** | | | | |
| 2 | 36.6, $CH_2$ | 2.12, m | 3, 4 | Gln-NH |
| 3 | 18.4. $CH_2$ | 1.53, m | 2, 4 | |
| 4 | 13.3, $CH_3$ | 0.88, ovlp | 2, 3 | |

**Table S2.** NMR data for micropeptin 966 (D-Gln) (**2**) (500 MHz for $^{1}H$ NMR, 125 MHz for $^{13}C$ NMR; DMSO-$d_6$).

| Position | δC, mult | δH, mult, *J* (Hz) | TOCSY | NOESY |
|---|---|---|---|---|
| **Val-1** | | | | |
| 2 | 54.2, CH | 4.74, m | 3, 4, 5, NH | |
| 3 | 30.3, CH | 2.07, m | 2, 4, 5, NH | |
| 4 | 18.7, $CH_3$ | 0.86, d (7.3) | 2, 3, 5, NH | |
| 5 | 16.8, $CH_3$ | 0.72, d (6.8) | 2, 3, 4, NH | |
| NH | | 7.44, ovlp[a] | 2, 3, 4, 5 | *N*-MePhe-2 |
| ***N*-MePhe-1** | | | | |
| 2 | 60.6, CH | 5.04, m | 3a, 3b | Val-NH, Phe-2, 3a |
| 3a | 33.6, $CH_2$ | 3.23, m | 2 | |
| 3b | | 2.87, m | 2 | |
| 4 | | | | |
| 5/9 | 129.2, CH | 7.25, d (7.5) | 6, 7, 8 | Phe-3a |
| 6/8 | 128.3, CH | 7.42, t (7.5) | 5, 7, 9 | |
| 7 | 125.7, CH | 7.32, d (7.4) | 5, 6, 8, 9 | |
| *N*-Me | 30.0, $CH_3$ | 2.79, s | | |
| **Phe[1]-1** | | | | |
| 2 | 49.5, CH | 4.75, ovlp | 3a, 3b | *N*-MePhe-2, Ahp-5, NH |
| 3a | 34.7, $CH_2$ | 2.83, m | 2 | *N*-MePhe-5, 9, Ahp-5 |
| 3b | | 1.67, m | 2 | Ahp-5 |
| 4 | | | | |
| 5/9 | 129.2, CH | 6.78, d (7.6) | 6, 7, 8 | |
| 6/8 | 127.5, CH | 7.17, t (7.3) | 5, 7, 9 | |
| 7 | 125.9, CH | 7.13, d (7.3) | 5, 6, 8, 9 | |
| **Ahp-1** | | | | |
| 2 | 48.4, CH | 3.60, m | 3a, 3b, 4a, NH | Ahp-3a, 3b, Ahp-NH |
| 3a | 20.9, $CH_2$ | 2.40, m | 2, 3b, 4a, 4b, 5, NH | Ahp-NH |
| 3b | | 1.59, m | 2, 3a | Ahp-2 |
| 4a | 29.2, $CH_2$ | 1.66, ovlp | 2, 3a, 4b, NH | Ahp-3a, 5, OH |
| 4b | | 1.51, m | 4a, 5, NH | Ahp-5 |
| 5 | 73.5, CH | 5.04, ovlp | 3a, 4a, 4b, OH | Ahp-4a, 4b, Phe[1]-2, 3a, 3b, 4a |
| NH | | 7.10, d (9.3) | 2, 3a, 4a, 4b | Ahp-2, 3a, Phe[2]-NH |
| OH | | 6.06, br | 5 | Ahp-4a |
| **Phe[2]-1** | | | | |
| 2 | 48.4, CH | 4.32, m | 3a, 3b, NH | |
| 3a | 36.9, $CH_2$ | 2.64, m | 2, 3b, NH | |
| 3b | | 1.79, m | 2, 3a, NH | |
| 4 | | | | |
| 5/9 | 123.8, CH | 7.11, ovlp | 6, 8 | |
| 6/8 | 122.6, CH | 7.08, ovlp | 5, 7 | |
| 7 | 122.2, CH | 6.91, m | | |
| NH | | 8.59, d (8.3) | 2, 3a, 3b | Ahp-NH, Thr-2, 3 |
| **Thr-1** | | | | |
| 2 | 54.6, CH | 4.53, m | NH | Phe-NH |
| 3 | 71.3, CH | 5.39, m | 4 | Phe-NH |
| 4 | 17.5, $CH_3$ | 1.18, d (6.5) | 3 | |
| NH | | 7.91, m | 2 | Gln-2 |
| **Gln-1** | | | | |
| 2 | 52.1, CH | 4.41, m | 3a, 3b, 4, NH | Thr-NH |
| 3a | 27.6, $CH_2$ | 1.89, m | 2, 3b, 4, NH | |
| 3b | | 1.75, m | 2, 3a, 4, NH | |
| 4 | 31.3, $CH_2$ | 2.11, m | 2, 3a, 3b, NH | |
| 5 | | | | |
| NH | | 8.10, d (8.0) | 2, 3a, 3b, 4 | BTA-2 |

| | | | | |
|---|---|---|---|---|
| $NH_2$ | | 7.28, br<br>6.78, br | | |
| **BTA-1** | | | | |
| 2 | 36.8, $CH_2$ | 2.13, m | 3, 4 | Gln-NH |
| 3 | 18.4. $CH_2$ | 1.54, m | 2, 4 | |
| 4 | 13.1, $CH_3$ | 0.89, d (7.3) | 2, 3 | |

[a]overlapping signals

**Table S3.** Molecular formulas, exact masses, and high-resolution mass spectrometric data for newly identified micropeptins and other cyanopeptides and confidence level of identification.

| Name | Molecular formula | Calcd $[M+H-H_2O]^+$ | Exact mass | calcd $[M+H]^+$ or $[M+Na]^+$ | found | Mass error (ppm) | Confidence Level |
|---|---|---|---|---|---|---|---|
| MP 1010 | $C_{53}H_{70}N_8O_{12}$ | 993.5080 | 1010.5113 | 1033.5005 | 1033.5009 | 0.39 | 1 |
| MP 966 (D-Gln) | $C_{51}H_{66}N_8O_{11}$ | 949.4818 | 966.4851 | 989.4743 | 989.4771 | 2.83 | 1 |
| MP 980 | $C_{52}H_{68}N_8O_{11}$ | 963.4975 | 980.5008 | 1003.4900 | 1003.4903 | 0.30 | 2 |
| MP 950 | $C_{47}H_{66}N_8O_{11}S$ | 933.4539 | 950.4572 | 973.4464 | 973.4456 | -0.82 | 2 |
| MP 1005 | $C_{53}H_{67}N_9O_{11}$ | 988.4927 | 1005.496 | 1028.4852 | 1028.4856 | 0.39 | 2 |
| MP 946 | $C_{49}H_{70}N_8O_{11}$ | 929.5131 | 946.5164 | 969.5056 | 969.5060 | 0.41 | 2 |
| MP 1038 | $C_{55}H_{74}N_8O_{12}$ | 1021.5393 | 1038.5426 | 1061.5318 | 1061.5318 | 0.00 | 2 |
| MP 1024 | $C_{54}H_{72}N_8O_{12}$ | 1007.5237 | 1024.5270 | 1047.5162 | 1047.5167 | 0.48 | 2 |
| [Leu$^1$[MC-LR] | $C_{52}H_{80}N_{10}O_{12}$ | | 1036.5957 | 1037.6030 | 1037.6030 | 0.00 | 1 |
| [Leu 1 ,Glu(OCH 3 ) 6 ] MC-LR | $C_{53}H_{82}N_{10}O_{12}$ | | 1050.6114 | 1051.6192 | 1051.6188 | -0.38 | 2 |
| Ferintoic acid C | $C_{46}H_{58}N_8O_9S$ | | 898.4047 | 899.4120 | 899.4128 | 0.89 | 1 |

**Table S4.** Sequences of micropeptin variants isolated/identified in this study (BTA=butanoic acid, HA=hexanoic acid).

| | Residue 1 | Residue 2 | Residue 3 | Residue 4 | Residue 5 | Residue 6 | Residue 7 | Side chain |
|---|---|---|---|---|---|---|---|---|
| MP 996 | Val | *N*Me-Phe | Phe | Ahp | Htyr | Thr | Gln | BTA |
| MP 982 | Val | *N*Me-Phe | Phe | Ahp | Tyr | Thr | Gln | BTA |
| MP 982 (L-Ser) | Val | *N*Me-Phe | Phe | Ahp | Htyr | **Ser** | Gln | BTA |
| MP 957 | Val | ***N*Me-Trp** | Phe | Ahp | **Val** | Thr | Gln | BTA |
| MP 1010 | Val | *N*Me-Phe | Phe | Ahp | **bHtyr** | Thr | Gln | BTA |
| MP 966 (D-Gln) | Val | *N*Me-Phe | Phe | Ahp | **Phe** | Thr | Gln | BTA |
| MP 980 | Val | *N*Me-Phe | Phe | Ahp | **Hphe** | Thr | Gln | BTA |
| MP 950 | Val | *N*Me-Phe | Phe | Ahp | **Met** | Thr | Gln | BTA |
| MP 1005 | Val | *N*Me-Phe | Phe | Ahp | **Trp** | Thr | Gln | BTA |
| MP 946 | Val | *N*Me-Phe | Phe | Ahp | **Hleu/ HIle** | Thr | Gln | BTA |
| MP 1038 | Val | *N*Me-Phe | Phe | Ahp | **bHtyr** | Thr | Gln | **HA** |
| MP 1024 | Val | *N*Me-Phe | Phe | Ahp | **Htyr** | Thr | Gln | **HA** |

**Table S5.** Marfey's derivatization data and assignments for **1–3.**

| **Amino Acid** | **$t_R$ (min)** | **Ferintoic acid C (3)** | **Micropeptin 966 (D-Gln) (2)** | **Micropeptin 1010 (1)** |
|---|---|---|---|---|
| L-Glutamine | 13.36 | | | |
| D-Glutamine | 16.99 | | | |
| L-Glutamic acid | 15.10 | | | 15.12 (L) |
| D-Glutamic acid | 16.61 | | 16.64 (D)<br>(from D-Gln) | |
| L-Serine | 13.09 | | | |
| D-Serine | 13.92 | | | |
| L-Tryptophan | 25.11 | 25.14 (L) | | |
| D-Tryptophan | 28.68 | | | |
| L-Valine | 20.86 | | 20.59 (L) | 20.67 (L) |
| D-Valine | 27.66 | | | |
| L-*N*-Me-Phenylalanine | 26.12 | | 25.74 (L) | 25.81 (L) |
| D-*N*-Me Phenylalanine | 27.48 | | | |
| L-Phenylalanine | 25.44 | 25.09 (L) | 25.00 (L) | 25.12 (L) |
| D-Phenylalanine | 30.95 | | | |
| L-Homotyrosine | 21.45 | 20.87 (L) | | |
| D-Homotyrosine | 24.37 | | | |
| L-Methionine | 20.52 | 20.96 (L) | | |
| D-Methionine | 26.22 | | | |
| L-Threonine | 12.86 | | 12.47 (L) | 12.54 (L) |
| L-allo-Threonine | 13.66 | | | |
| D-allo-Threonine | 15.73 | | | |
| D-Threonine | 17.66 | | | |
| L-*N*-Me-Alanine | 18.57 | 18.66 (L) | | |
| D-*N*-Me-Alanine | 18.98 | | | |
| L-Lysine | 6.20 | | | |
| D-Lysine | 5.56 | 5.58 (D) | | |
| Bis-homotyrosine (L-FDLA) | 17.61 | | | (L) |
| Bis-homotyrosine (D-FDLA) | 18.89 | | | |

**Table S6.** Detection of micropeptins in Lake Erie samples. Retention times of micropeptin stereoisomers and field sample peaks under LC-MSD conditions (Luna C18, 150 × 2 mm, 5 µm, 55% A / 45% B, 0.6 mL/min).

| **Sample / Standard** | **Name** | ***m/z*** | **$t_R$ (min)** |
|---|---|---|---|
| Standard | Micropeptin 982 (L-Gln) | 1005 | 1.80 |
| Standard | Micropeptin 982 (L-Ser) | 1005 | 2.56 |
| Standard | Micropeptin 982 (D-Gln) | 1005 | 2.72 |
| Standard | Micropeptin 982 (L-*allo*-Thr) | 1005 | 2.90 |
| Field Extract (Huntington) | Micropeptin 982 (L-Gln) | 1005 | 1.86 |
| Standard | Micropeptin 996 (L-Gln) | 1019 | 2.75 |
| Standard | Micropeptin 996 (D-Gln) | 1019 | 2.86 |
| Field Extract (Huntington) | Micropeptin 996 (L-Gln) | 1019 | 2.78 |

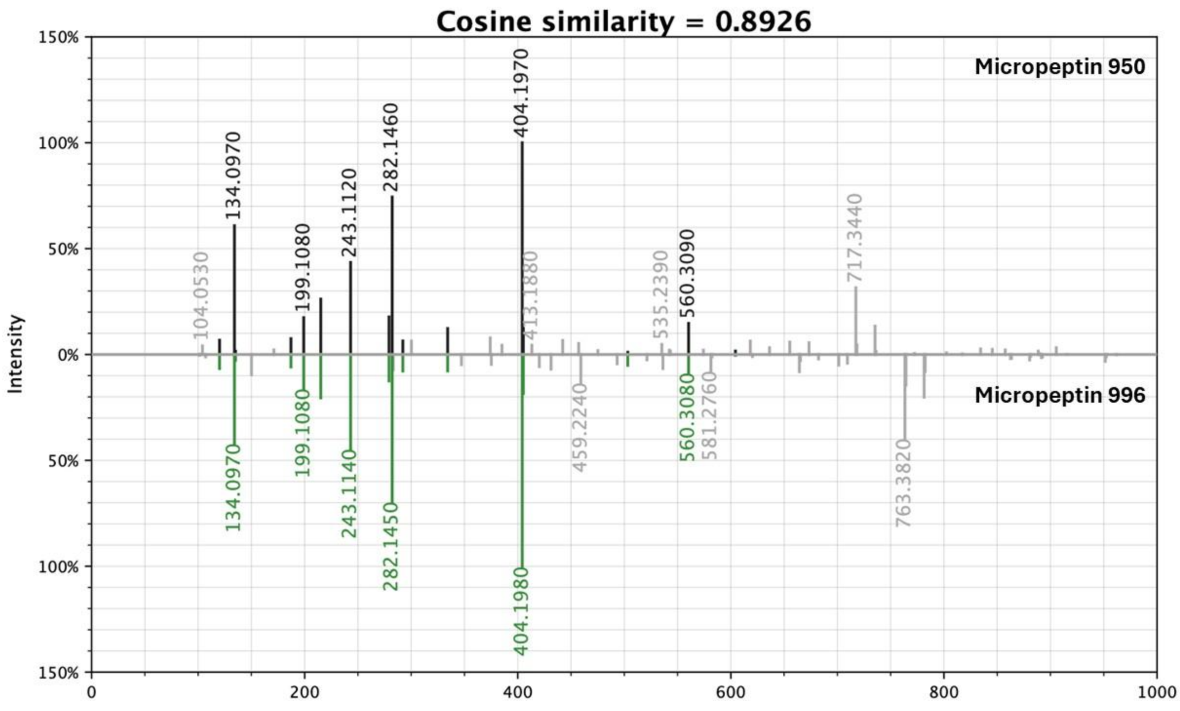


**Figure S1.** Mirror plot of MS/MS spectra of micropeptin 950 and micropeptin 996.

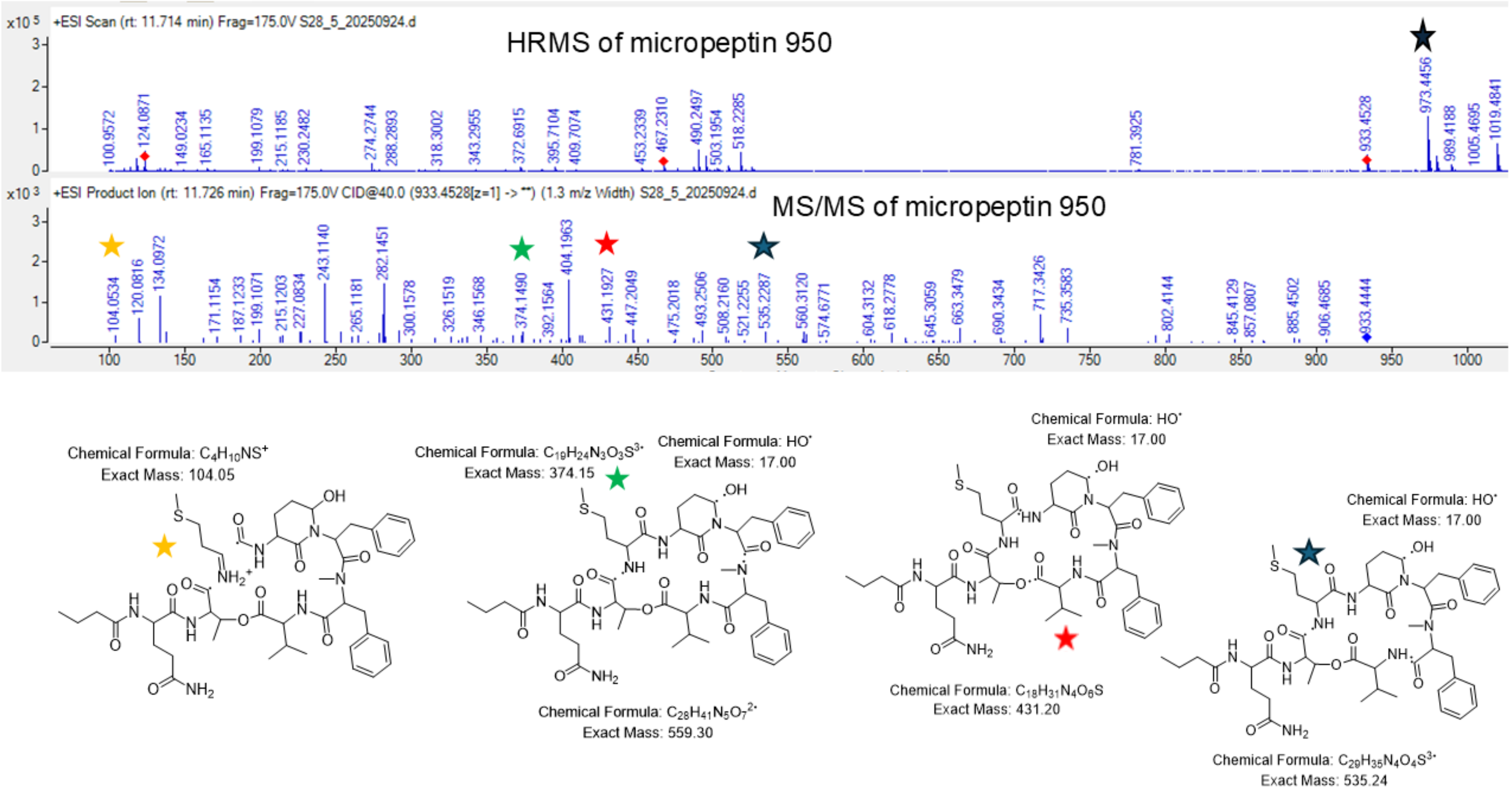


**Figure S2.** HRMS measurement (*m/z* 973.4456) and annotated MS/MS fragmentation pattern of micropeptin 950.

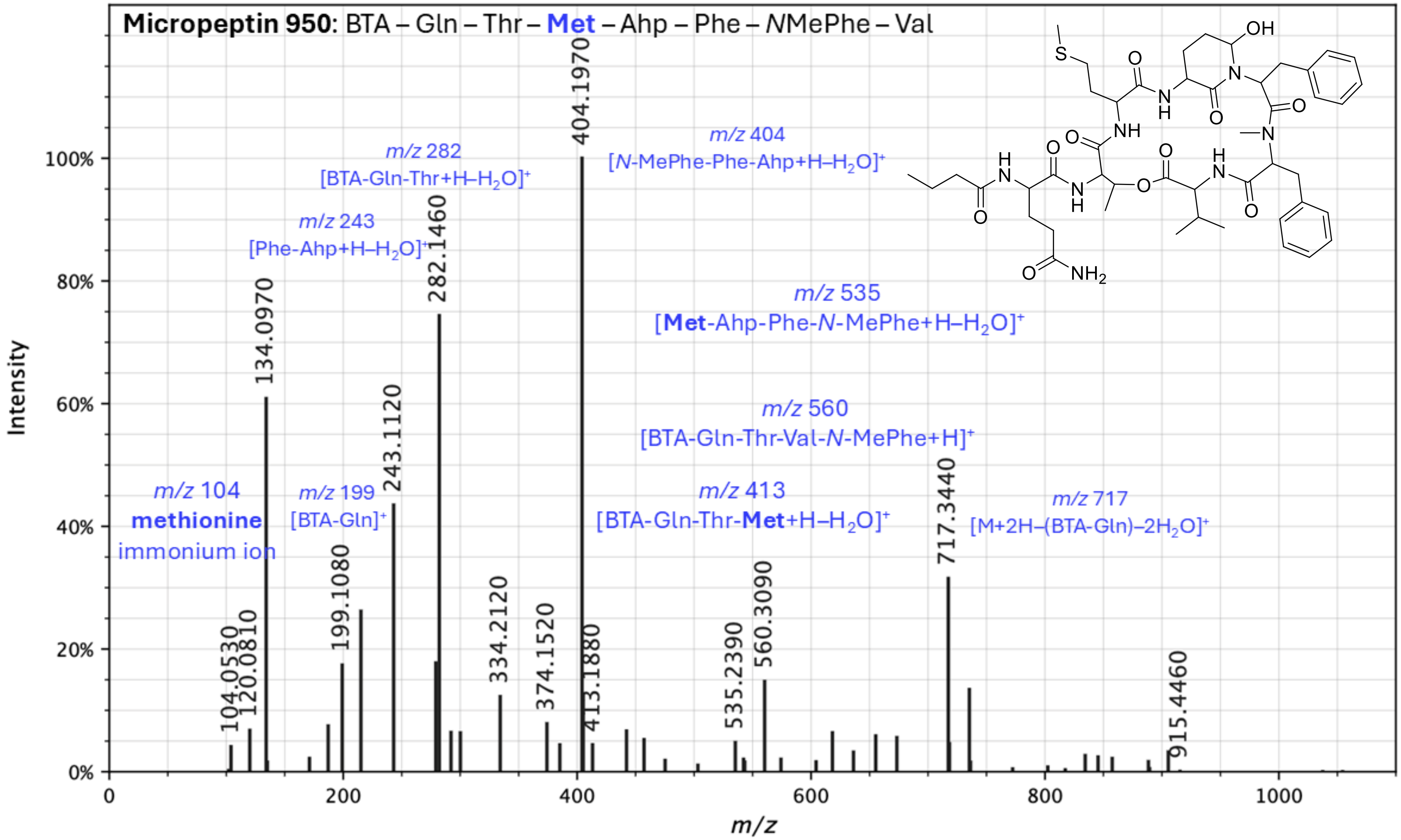


**Figure S3.** Annotated MS/MS spectrum of micropeptin 950.

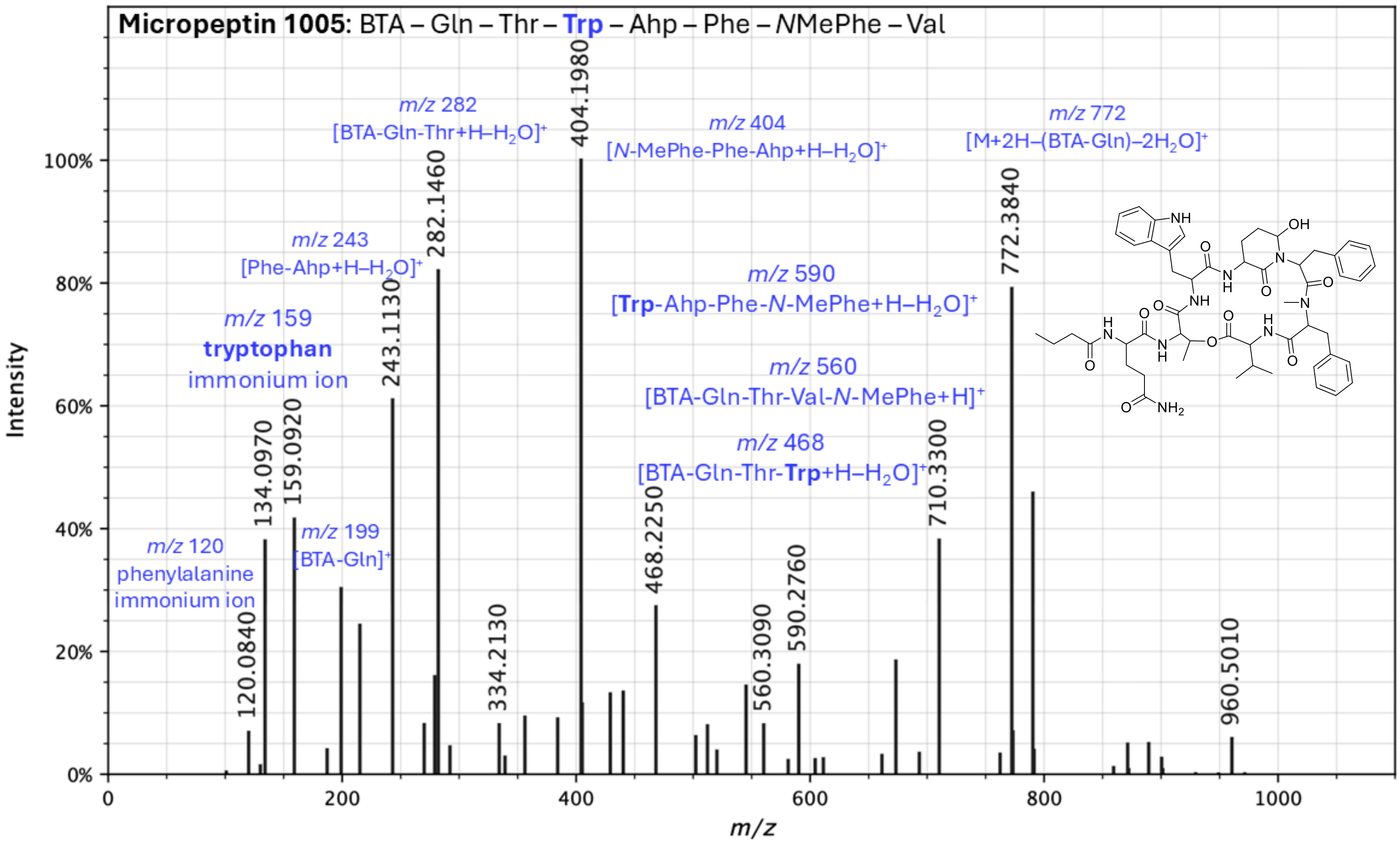


**Figure S4.** Annotated MS/MS spectrum of micropeptin 1005.

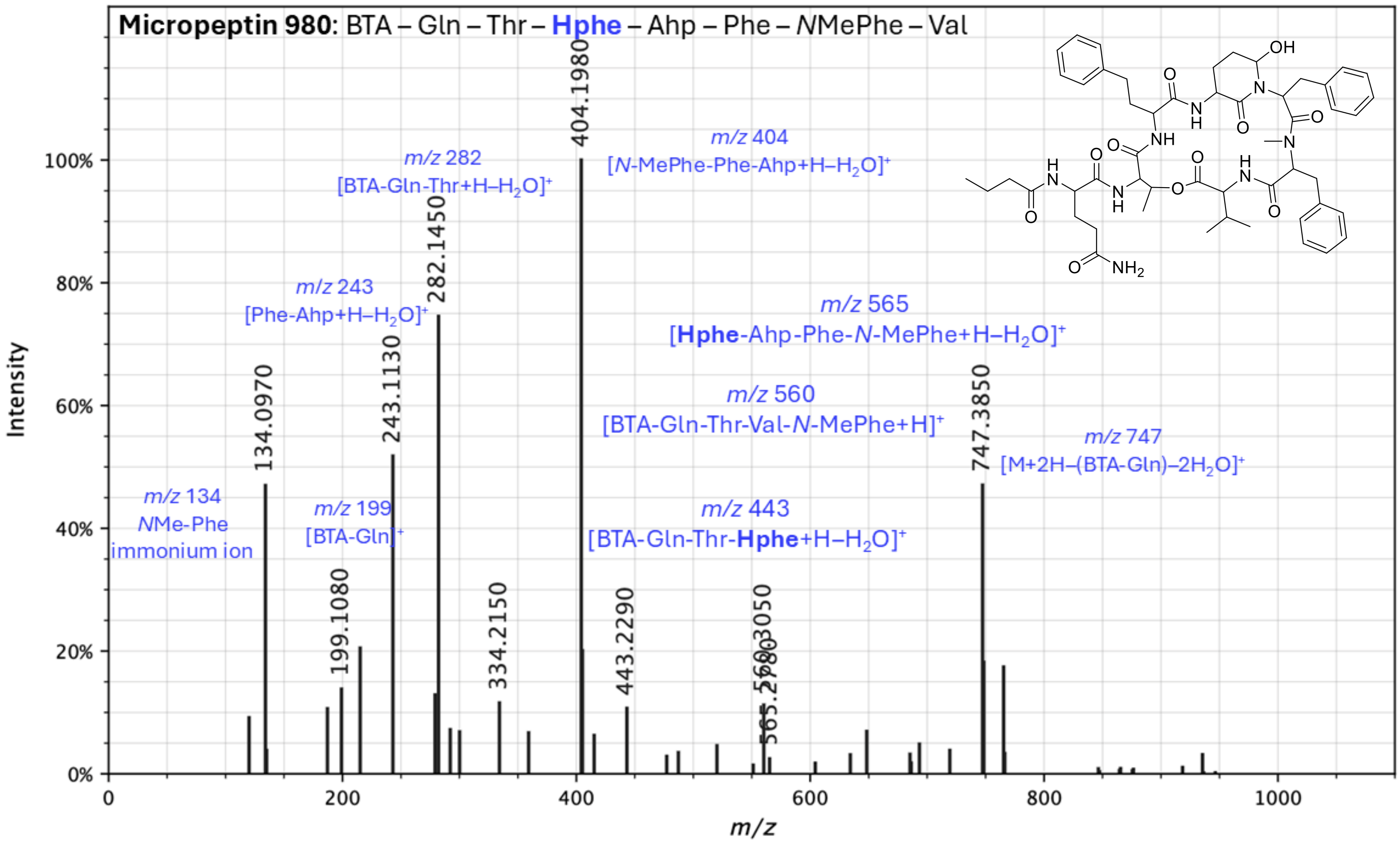


**Figure S5.** Annotated MS/MS spectrum of micropeptin 980.

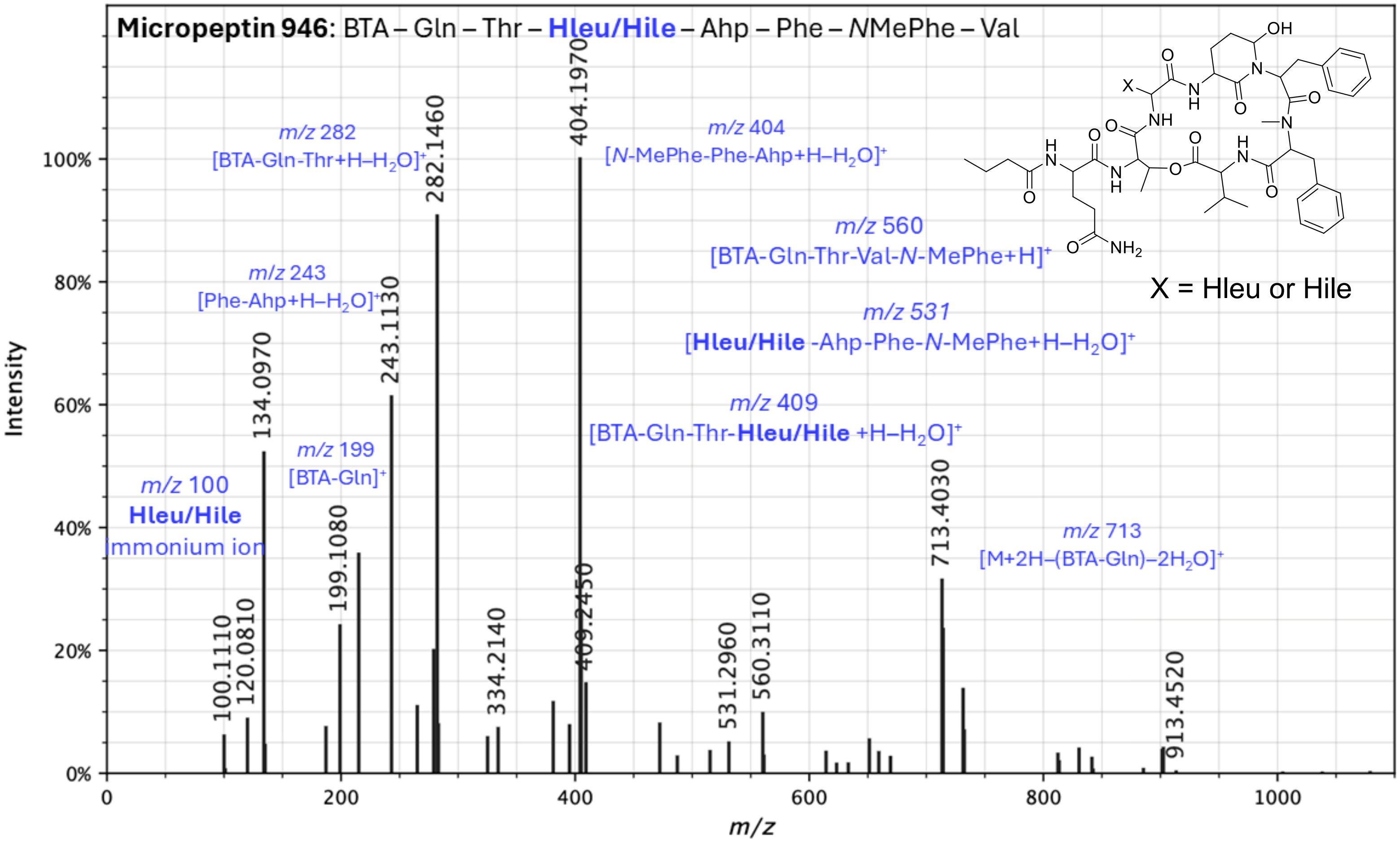


**Figure S6.** Annotated MS/MS spectrum of micropeptin 946.

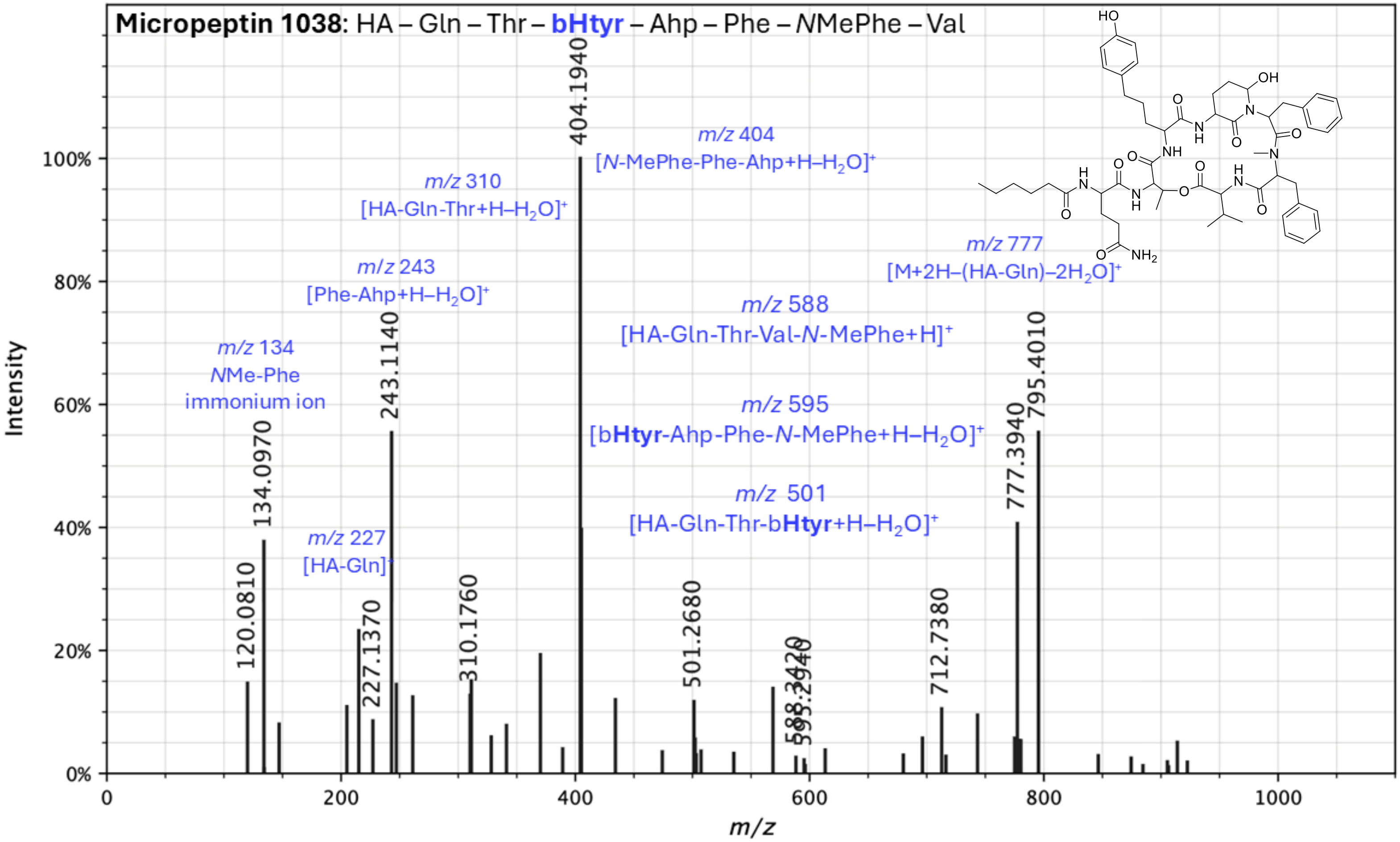


**Figure S7.** Annotated MS/MS spectrum of micropeptin 1038.

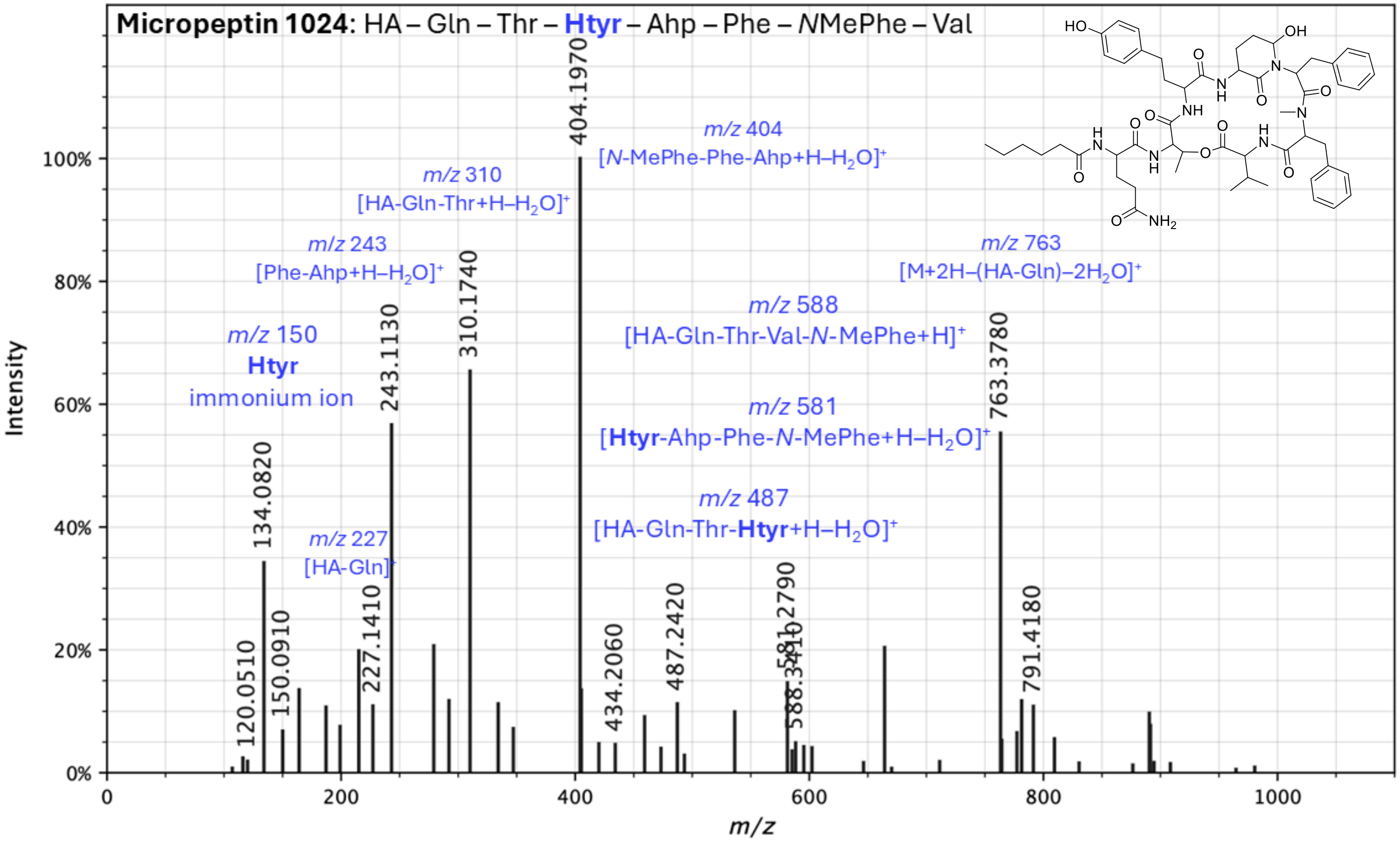


**Figure S8.** Annotated MS/MS spectrum of micropeptin 1024.

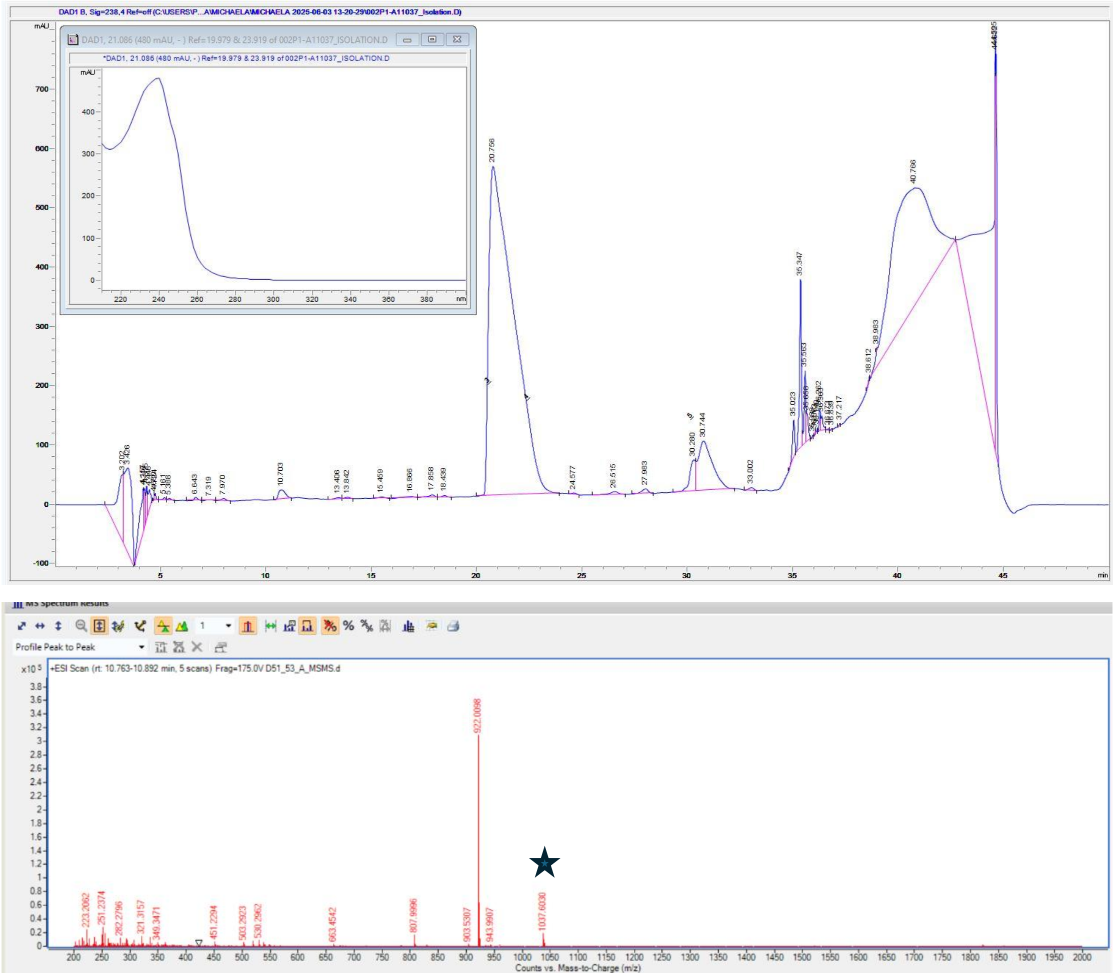


**Figure S9.** Isolation and characterization of [Leu[1]]MC-LR. Top panel: HPLC-DAD analysis with UV spectrum with $\lambda_{max}$ of 238 nm (inset) of [Leu[1]]MC-LR. Bottom panel: LC-HRMS analysis of [Leu[1]]MC-LR with *m/z* 1037.6030 (star).

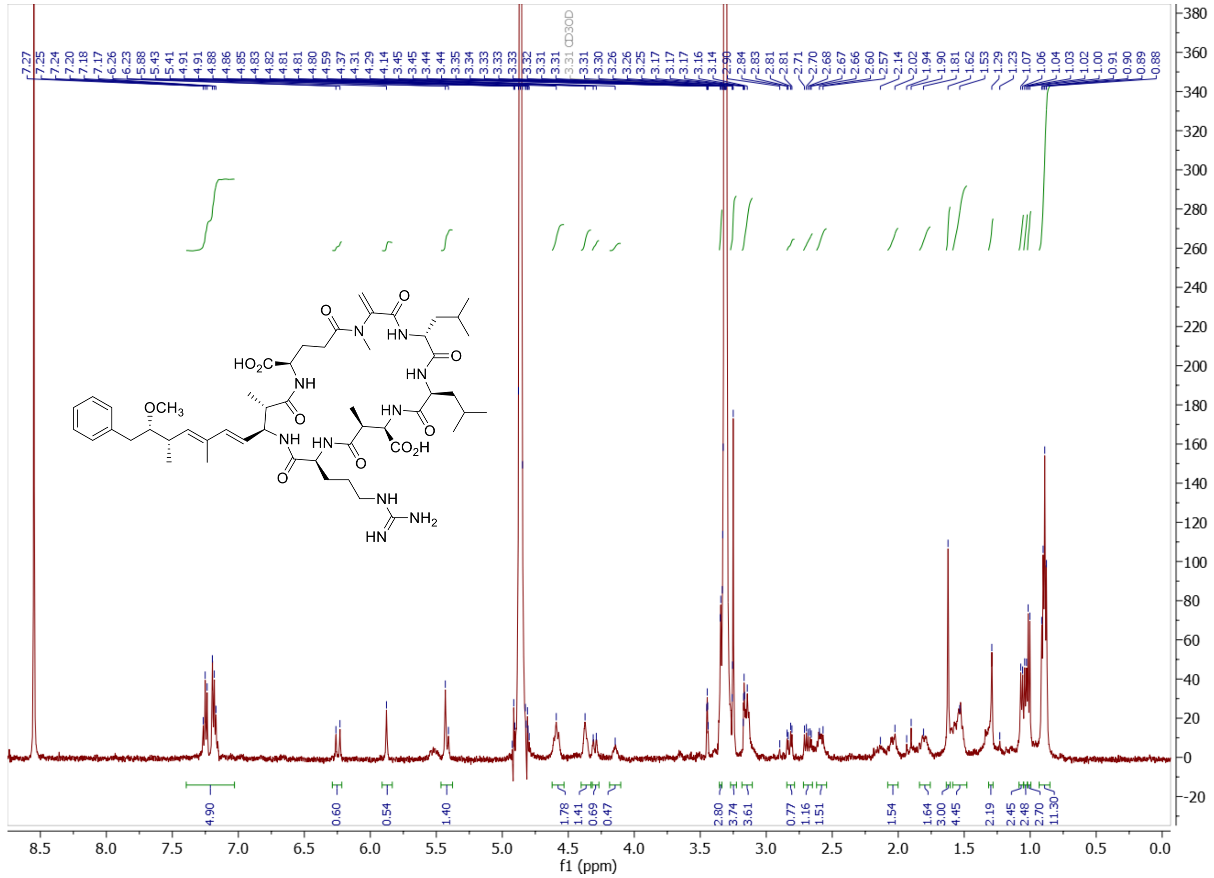

**Figure S10.** $^{1}H$ NMR (500 MHz, MeOH-$d_4$) of [Leu[1]]MC-LR.

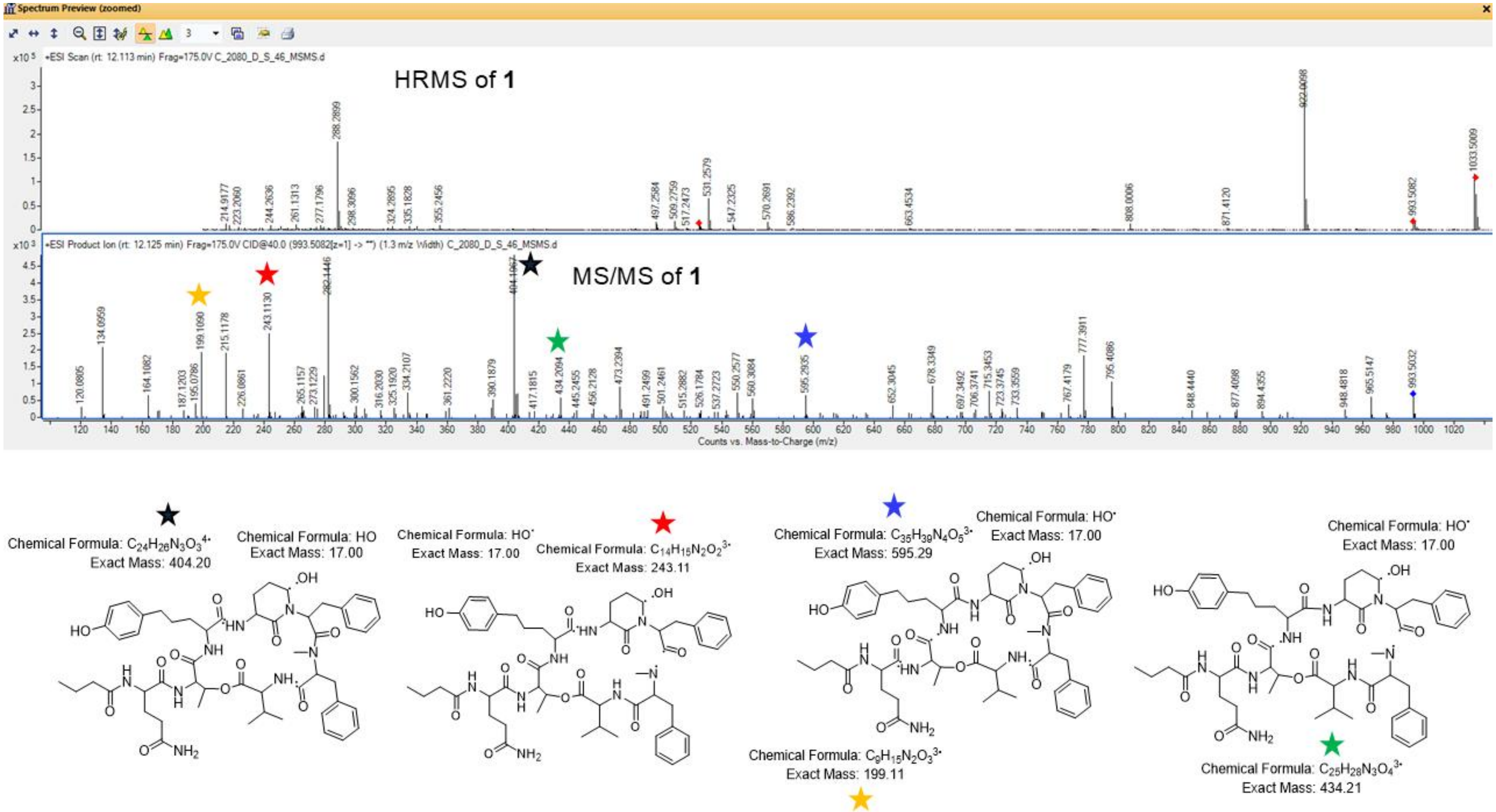


**Figure S11.** Mass spectrometry data of **1**. Top panel: HRMS of compound **1** *m/z* 1033.5009 $[M+Na]^+$. Middle panel: MS/MS of **1** with key fragmentation ions noted with stars, which correspond to the putative fragmentations illustrated in the bottom panel.

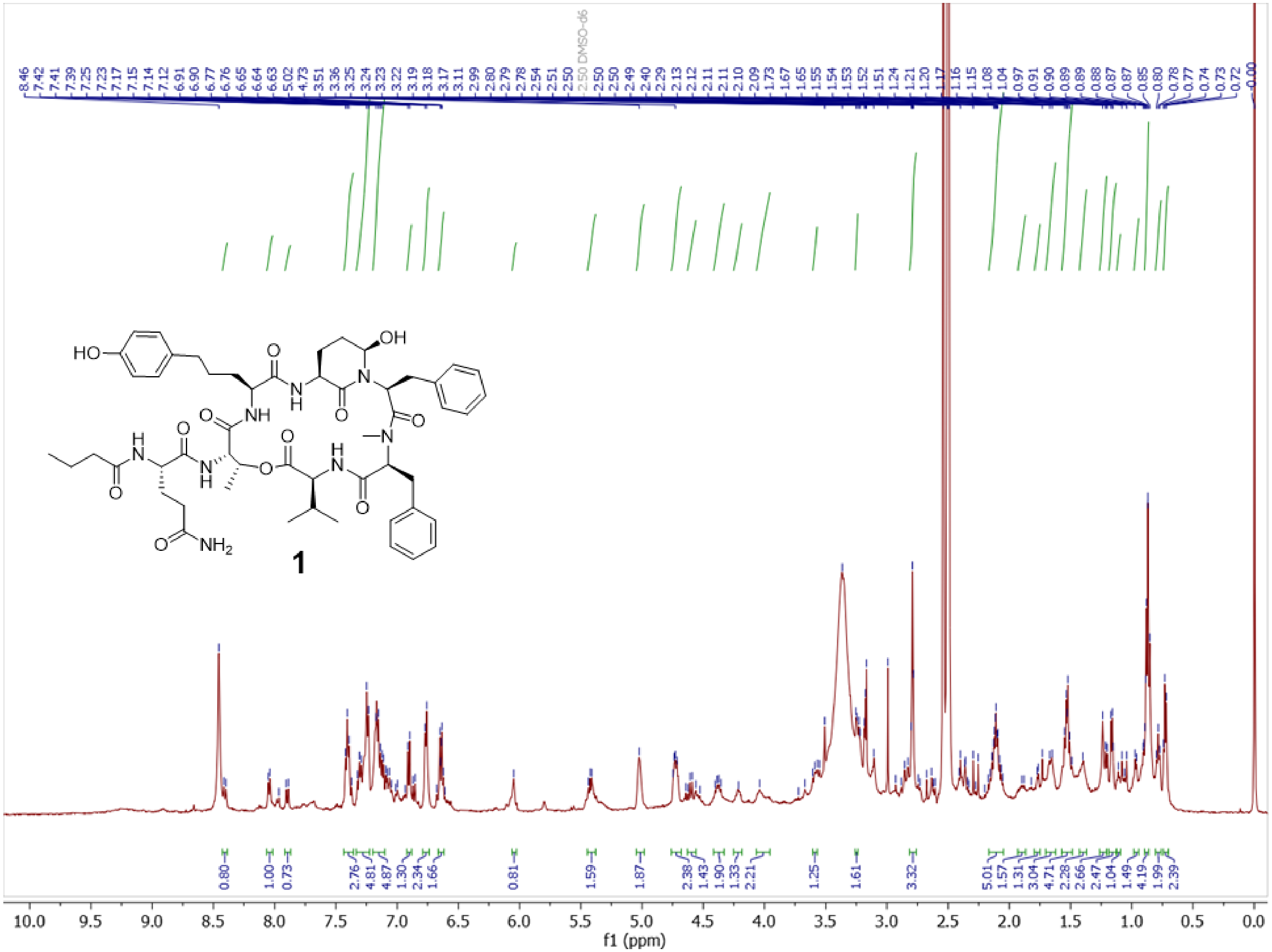

**Figure S12.** $^{1}$H NMR (500 MHz, DMSO-$d_6$) of micropeptin 1010 (**1**).

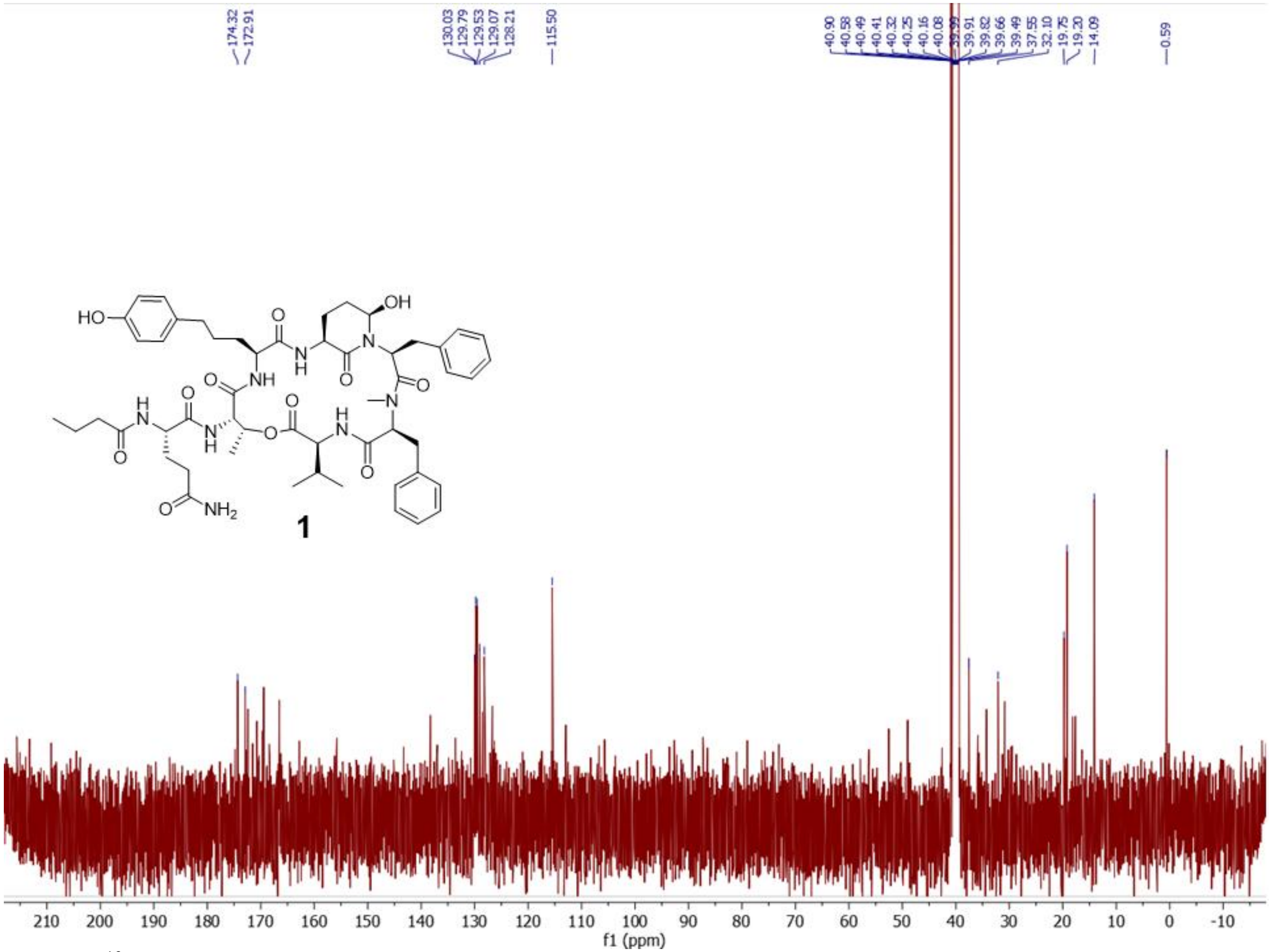

**Figure S13.** $^{13}C$ NMR (125 MHz, DMSO-$d_6$) of micropeptin 1010 (**1**).

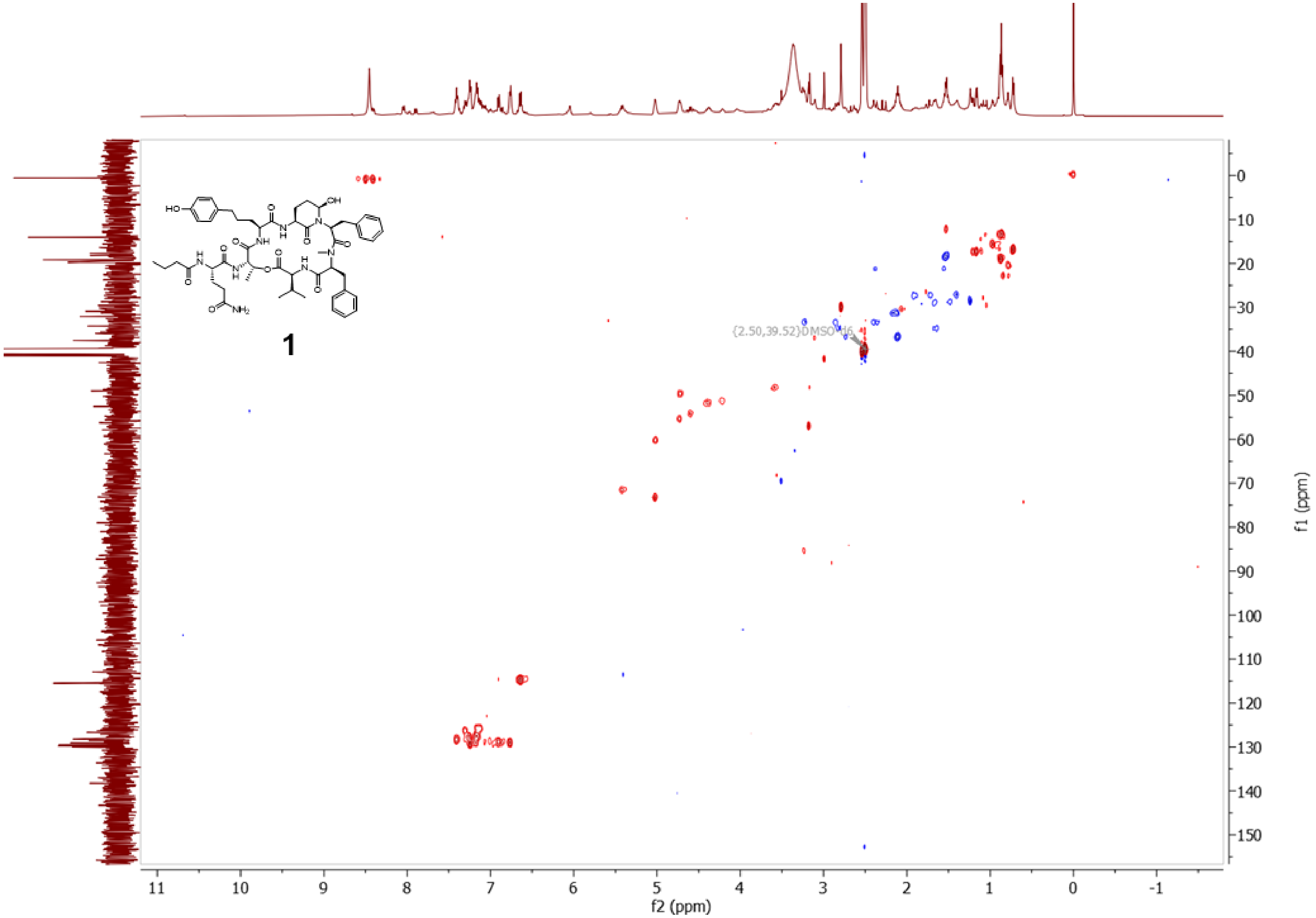


**Figure S14.** Multiplicity-edited HSQC of micropeptin 1010 (**1**).

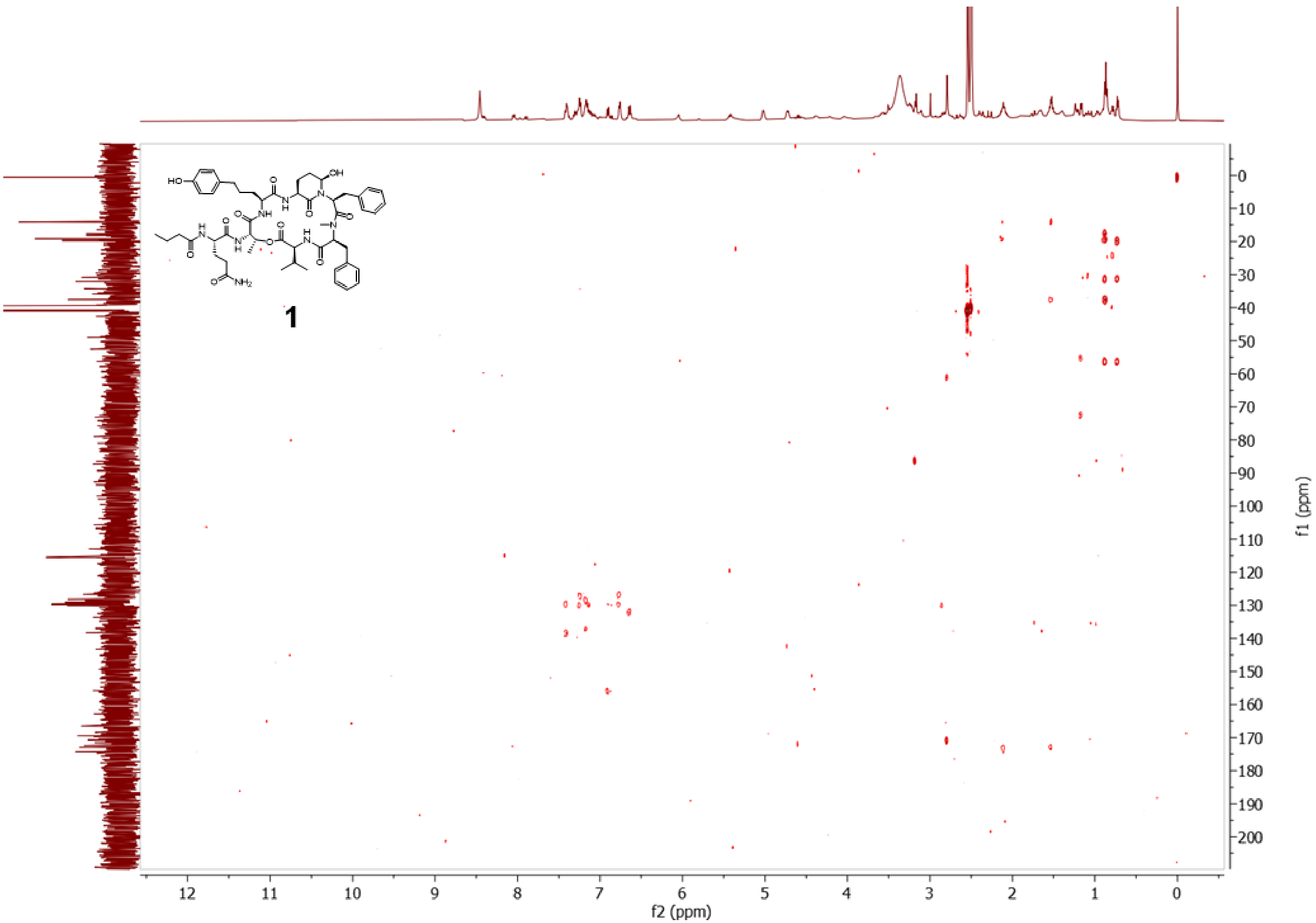


**Figure S15.** HMBC of micropeptin 1010 (**1**).

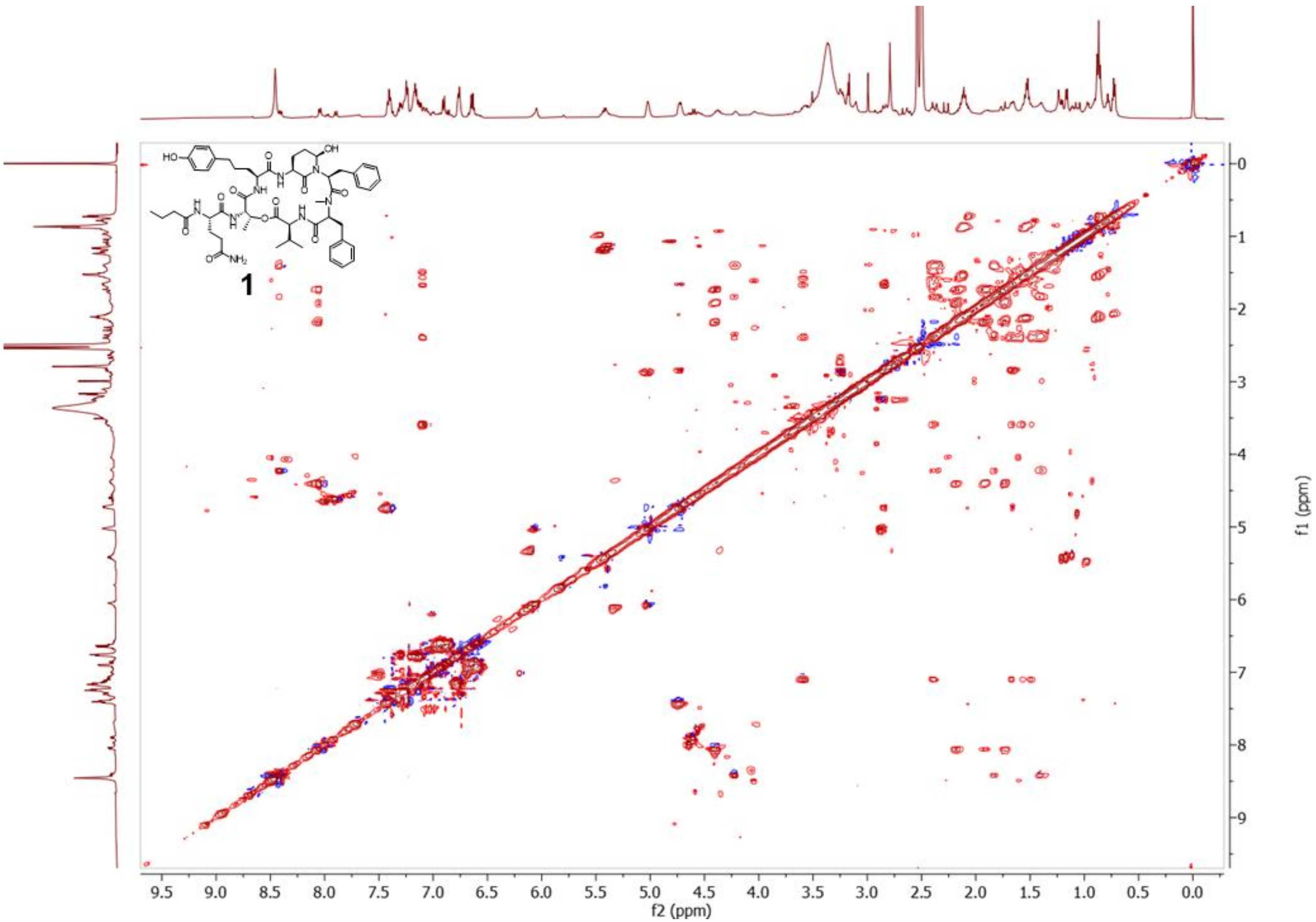


**Figure S16.** TOCSY of micropeptin 1010 (**1**).

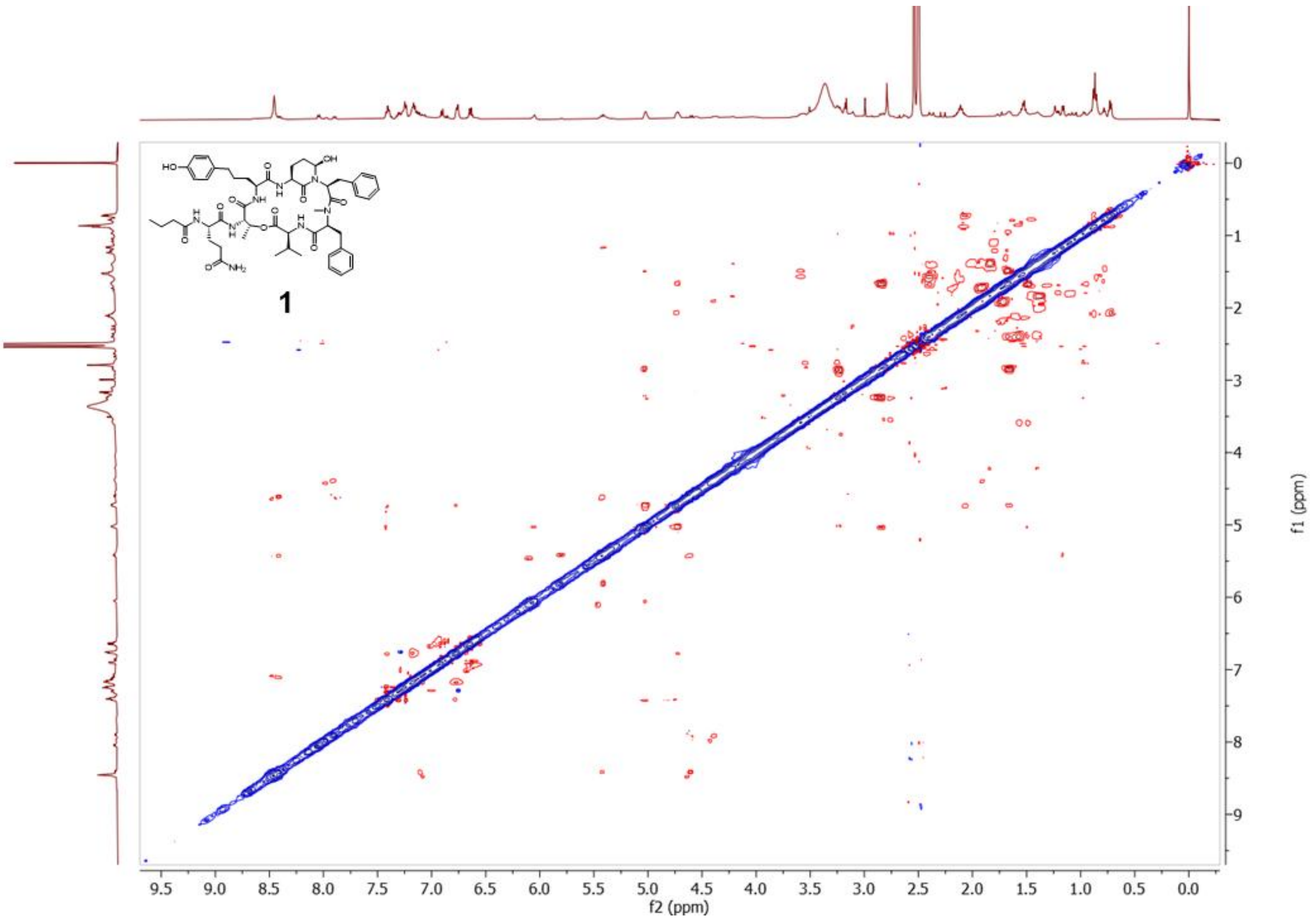


**Figure S17.** NOESY of micropeptin 1010 (**1**).

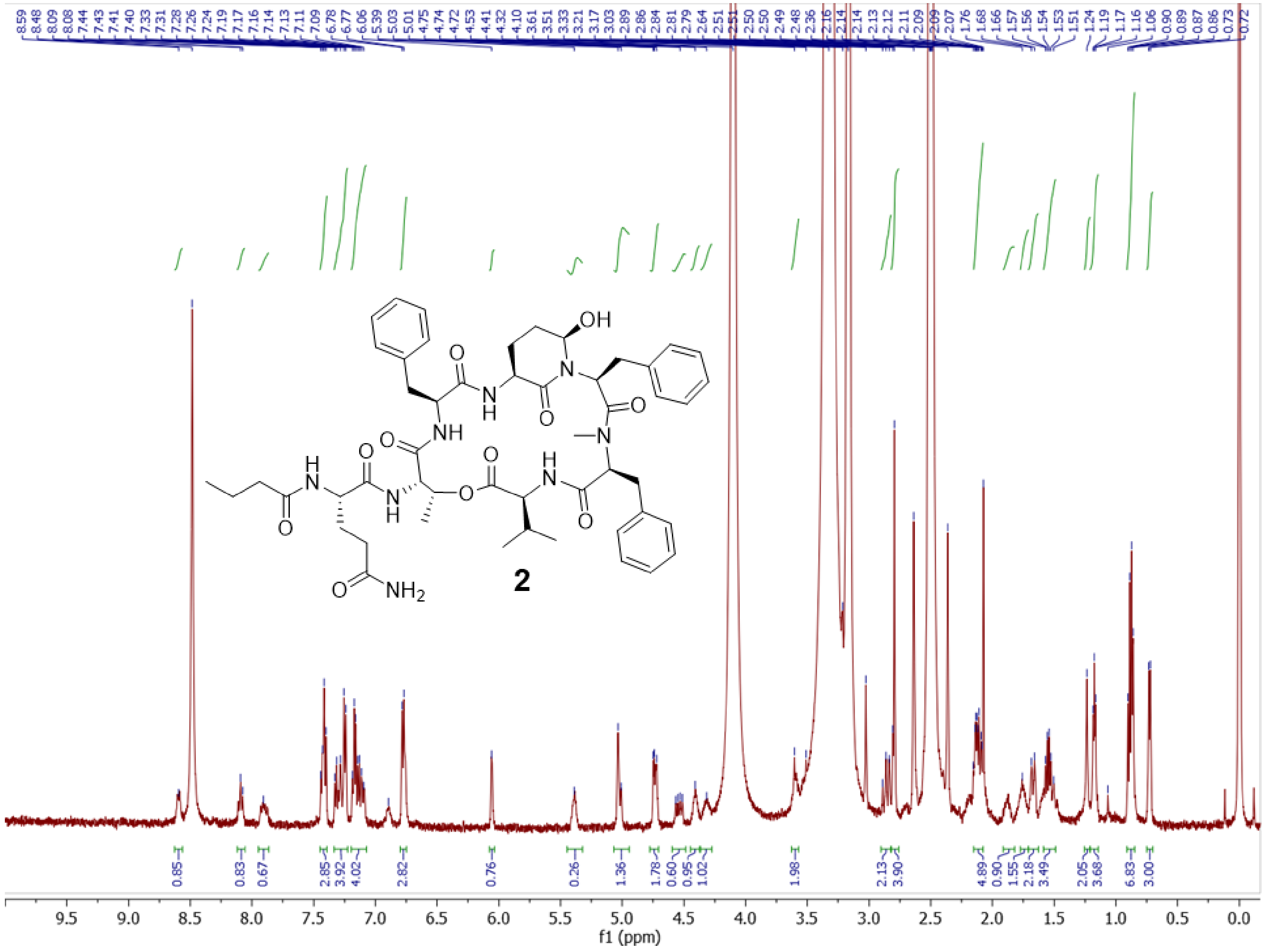

**Figure S18.** $^{1}$H NMR (500 MHz, DMSO-$d_6$) of micropeptin 966 (D-Gln) (**2**).

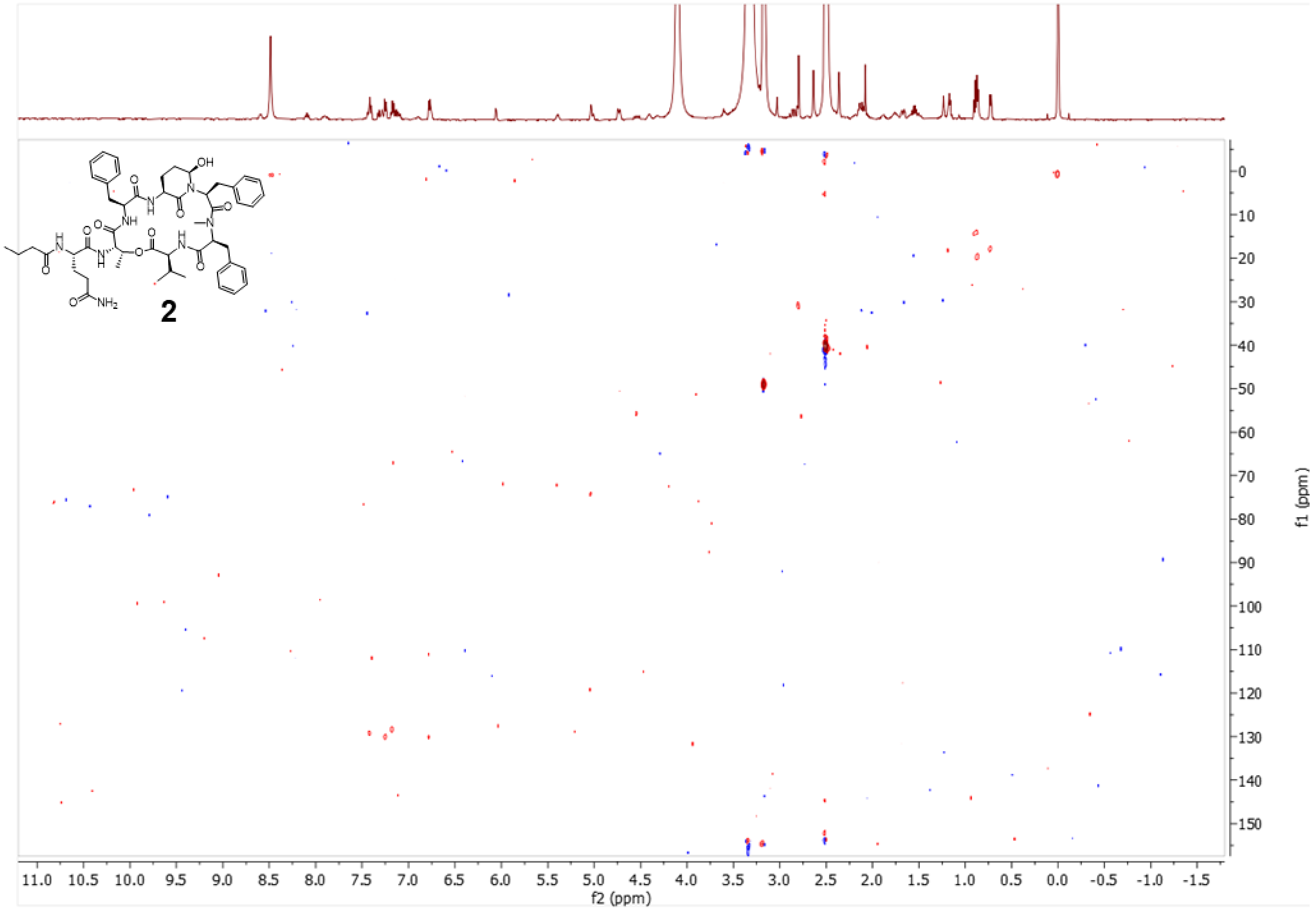

**Figure S19.** Multiplicity-edited HSQC of micropeptin 966 (D-Gln) (**2**).

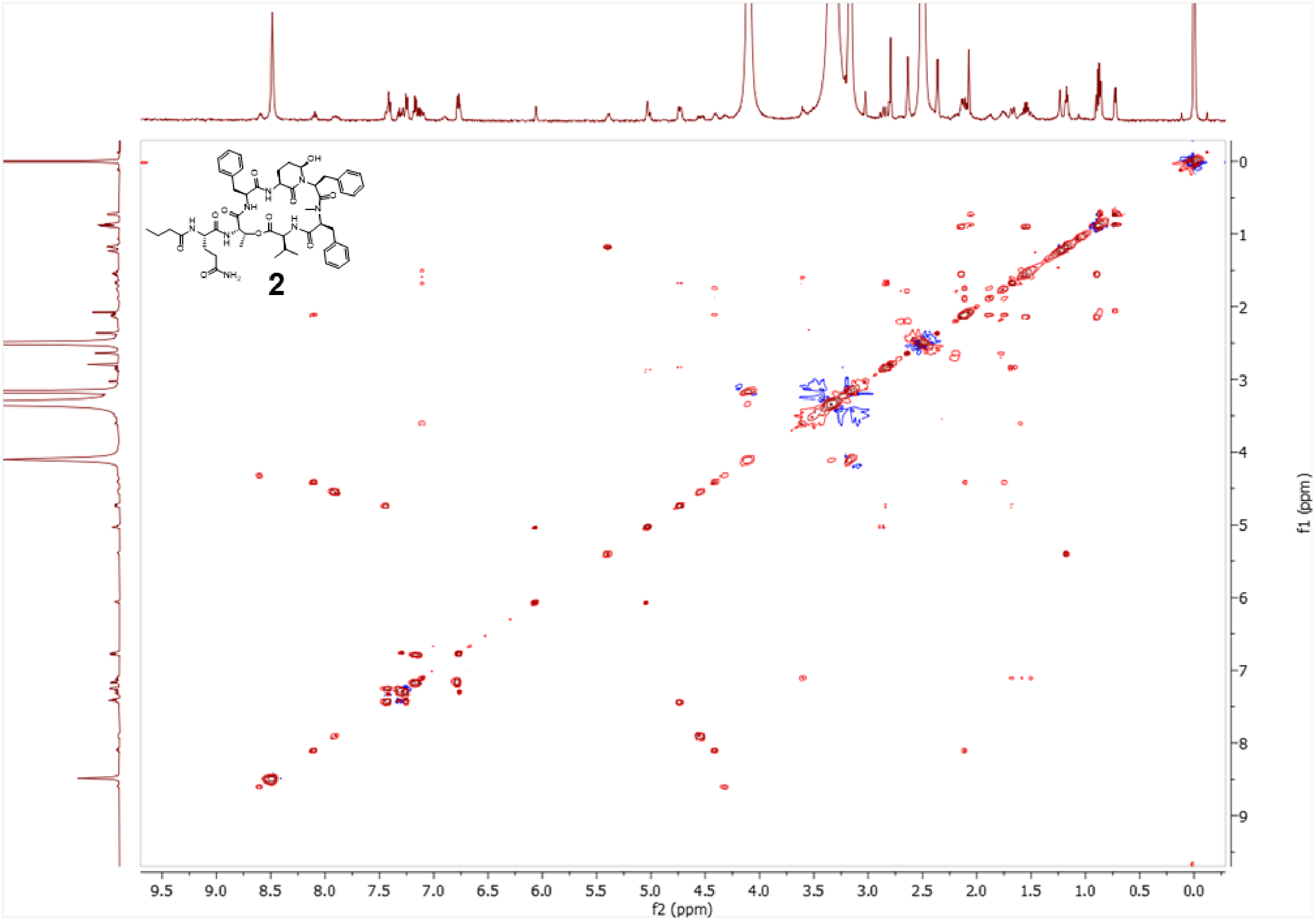


**Figure S20.** TOCSY of micropeptin 966 (D-Gln) (**2**).

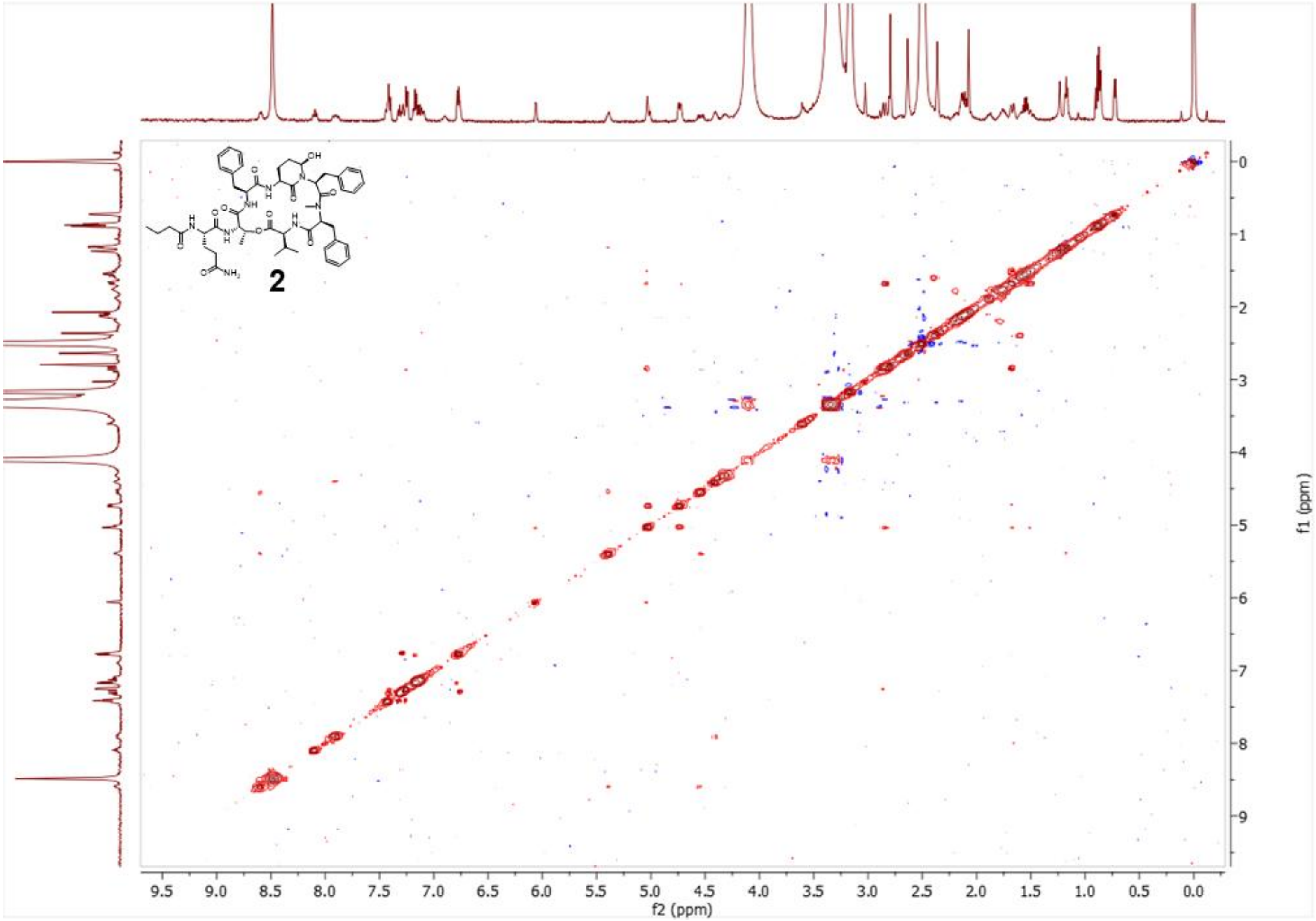


**Figure S21.** NOESY of micropeptin 966 (D-Gln) (**2**).

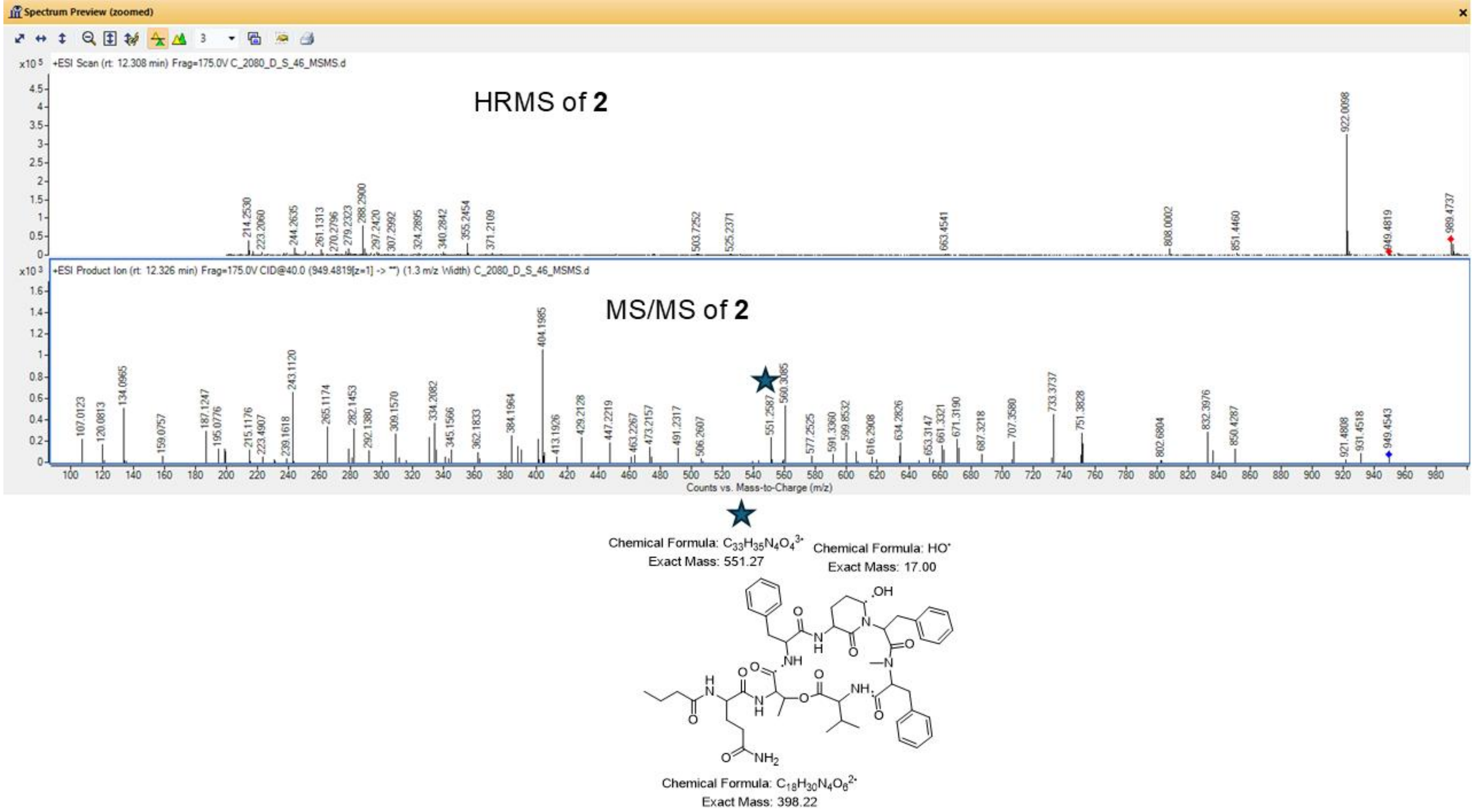


**Figure S22.** Mass spectrometry data of **2.** Top panel: HRMS of compound **2** *m/z* 989.4737 [M+Na]$^{+}$. Middle panel: MS/MS of **2** with a key fragmentation ion noted with a star, which corresponds to the putative fragmentation illustrated in the bottom panel.

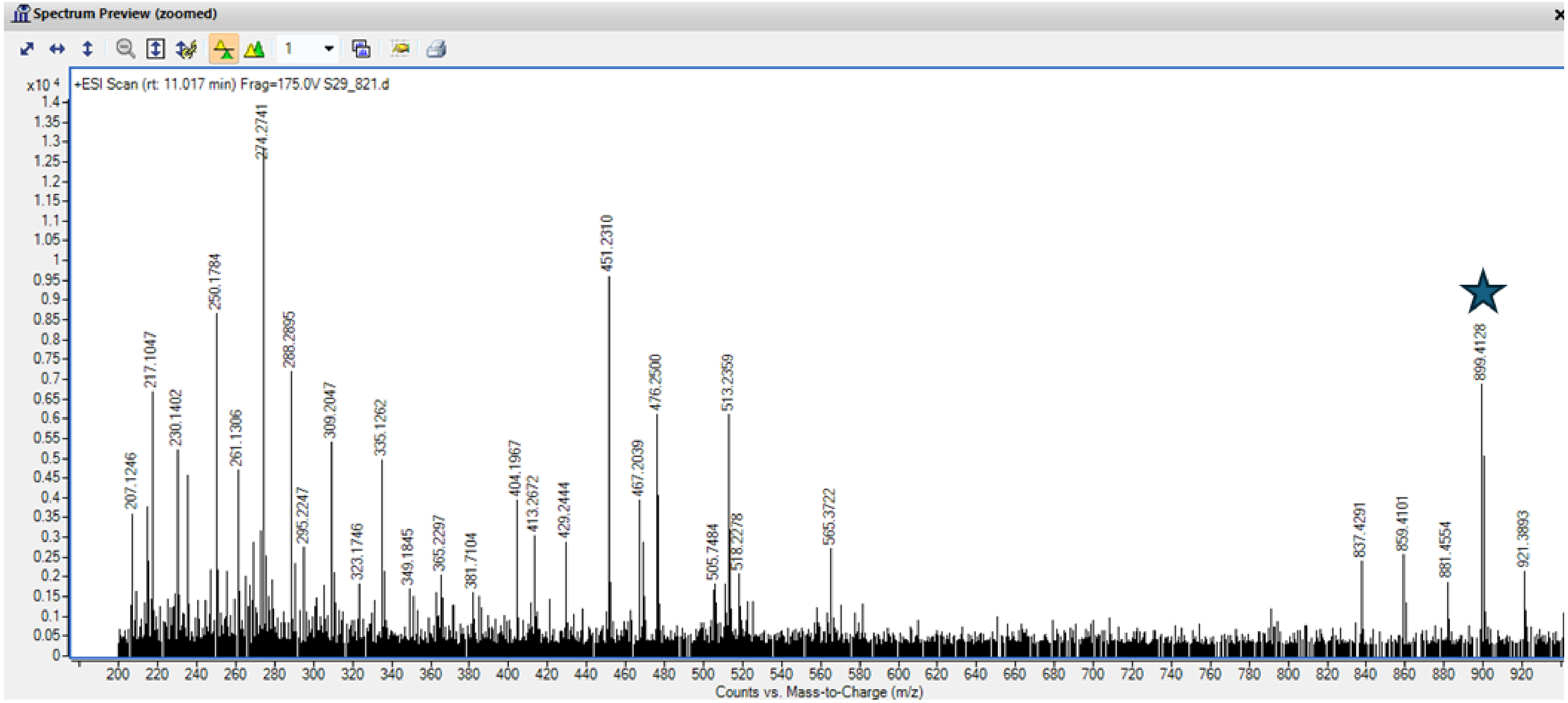


**Figure S23.** HRMS of ferintoic acid C *m/z* 899.4128 $[M+H]^+$.

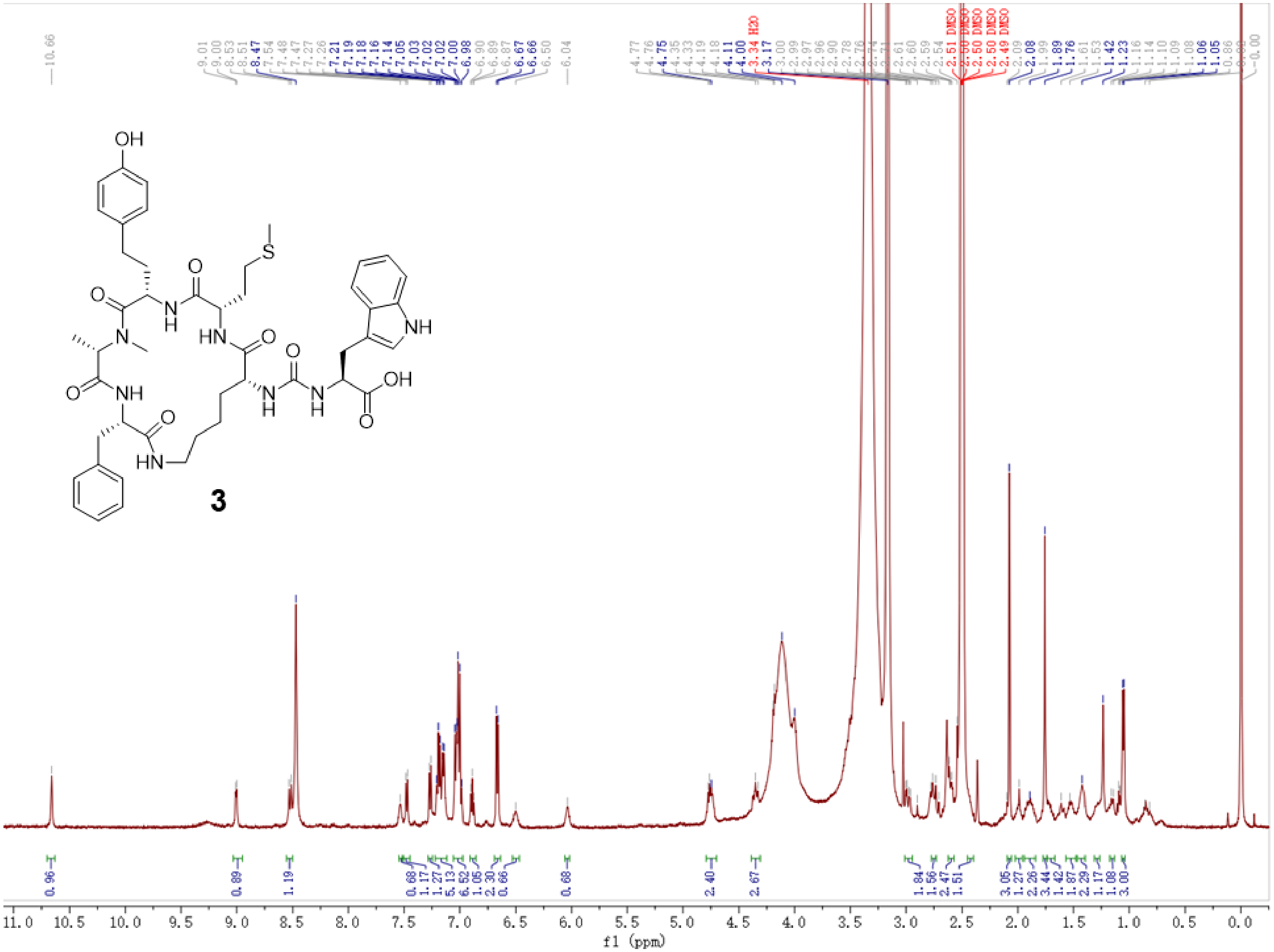


**Figure S24.** $^1$H NMR (500 MHz, DMSO-$d_6$) of ferintoic acid C (**3**).

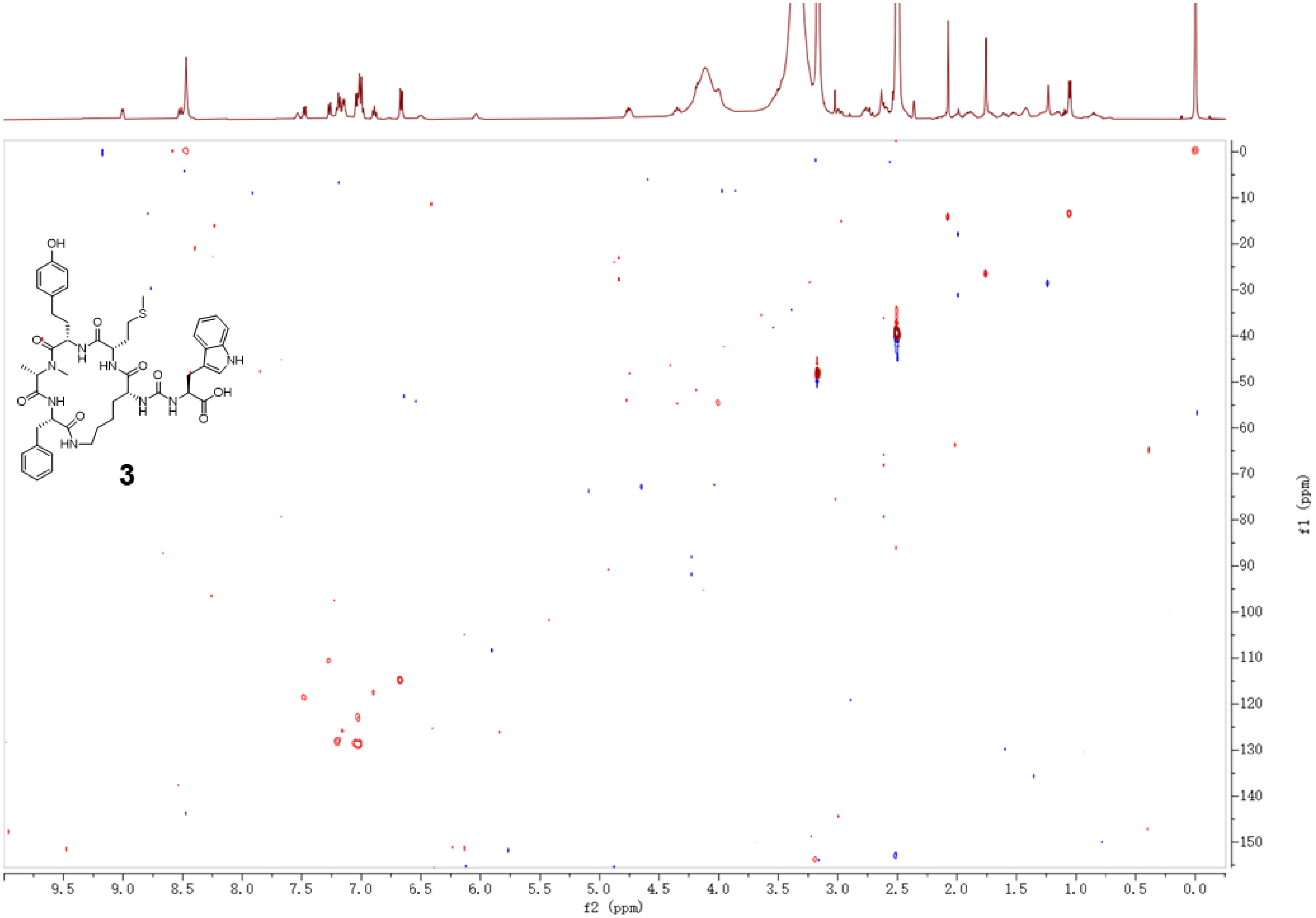


**Figure S25.** Multiplicity-edited HSQC of ferintoic acid C (**3**).

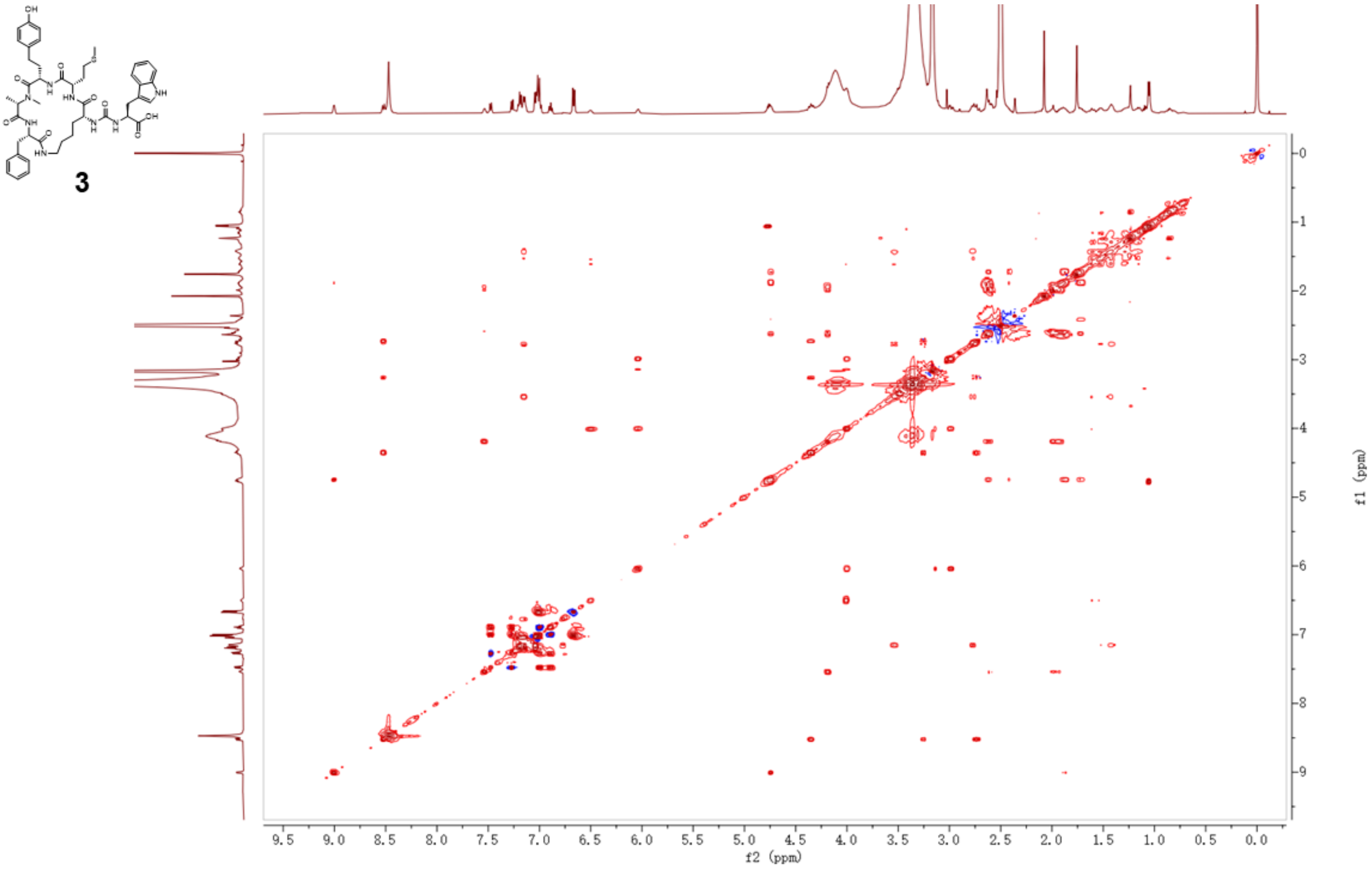


**Figure S26.** TOCSY of ferintoic acid C (**3**).

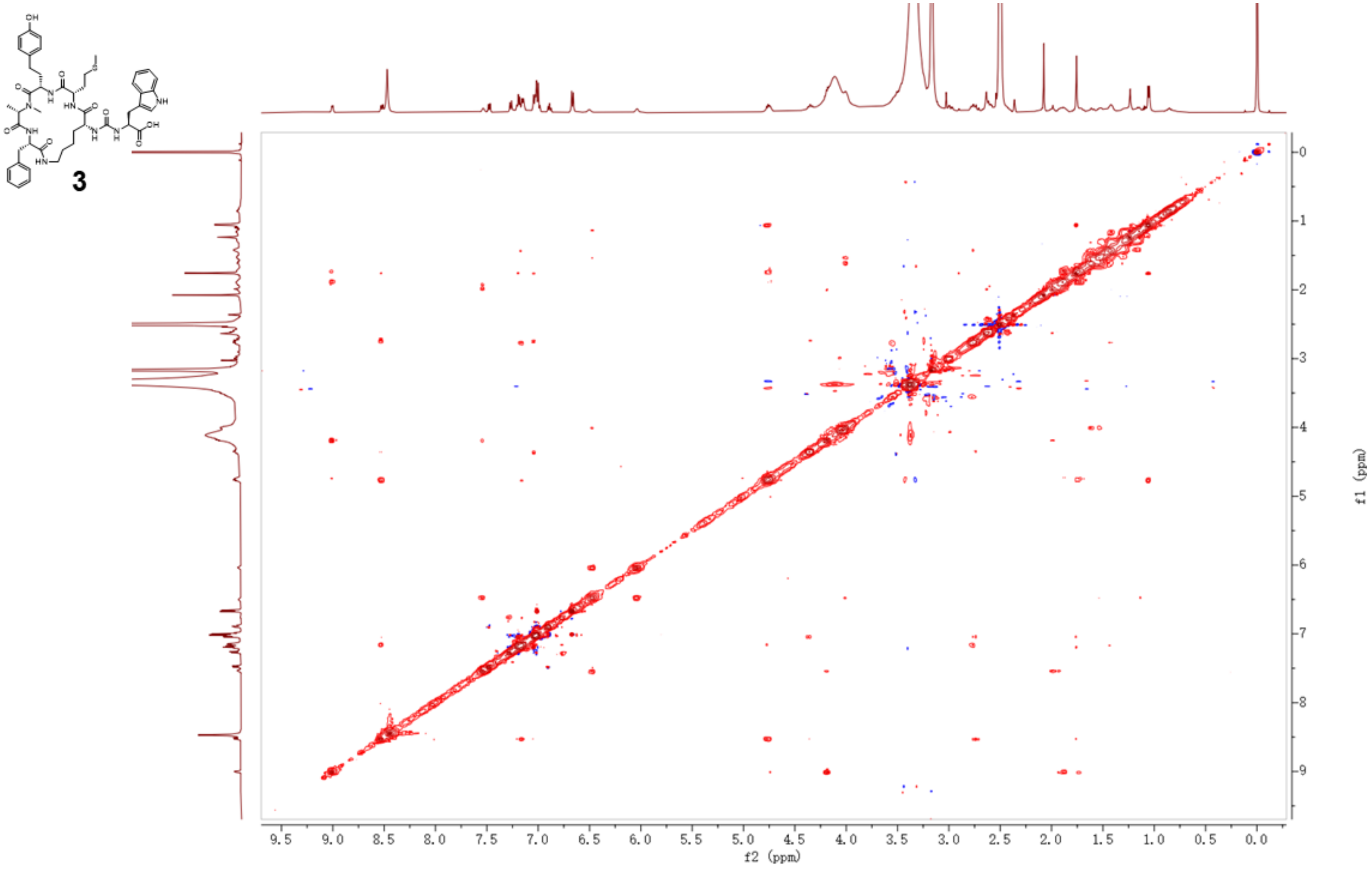


**Figure S27.** NOESY of ferintoic acid C (**3**).

Micropeptin 1010

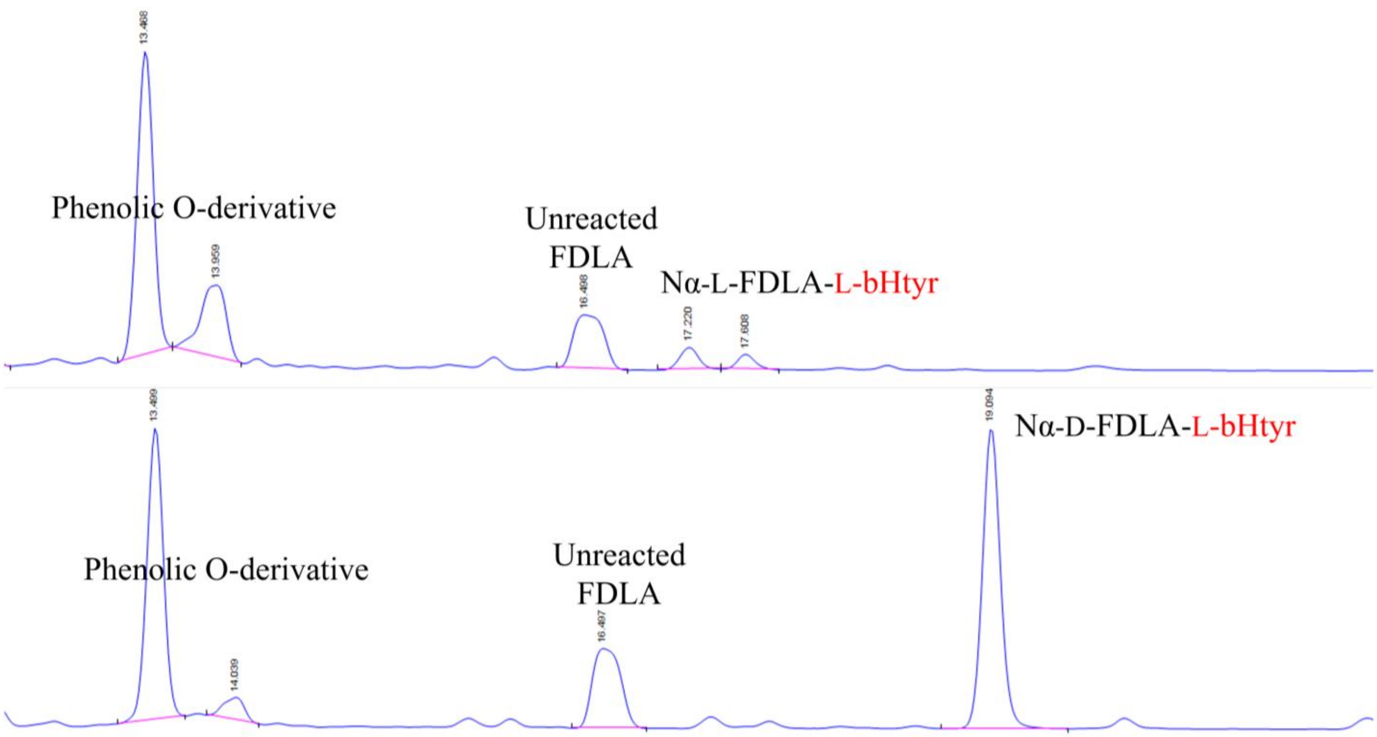


**Figure S28.** LC-MS analysis of the hydrolysate of **1** reacted with L-FDLA (top panel) and D-FDLA (bottom panel) to determine the configuration of the bHtyr in **1**.

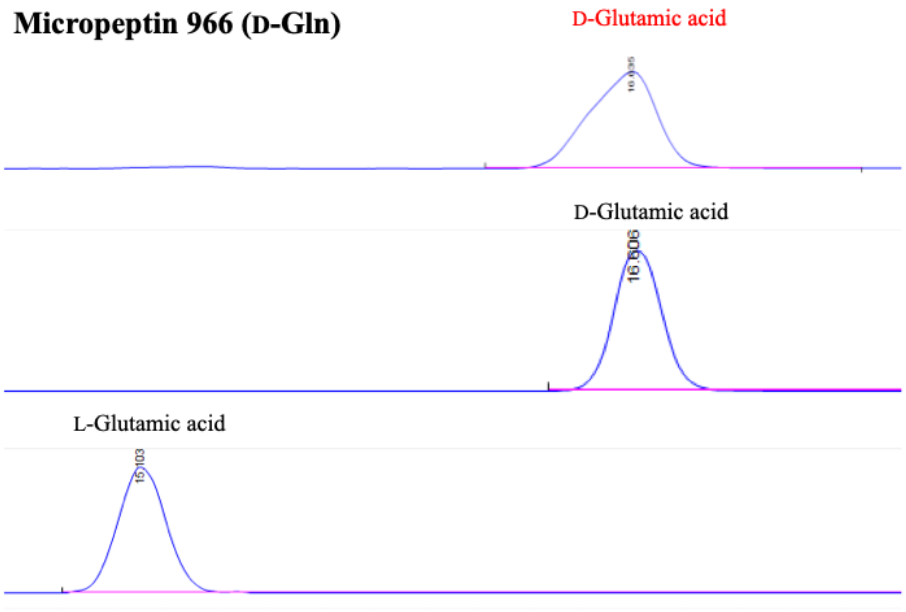


**Figure S29.** LC-MS analysis of the hydrolysate of **2** (top panel) and L- and D-Glutamic acid (middle and bottom panel, respectively) derivatized with L-FDVA.

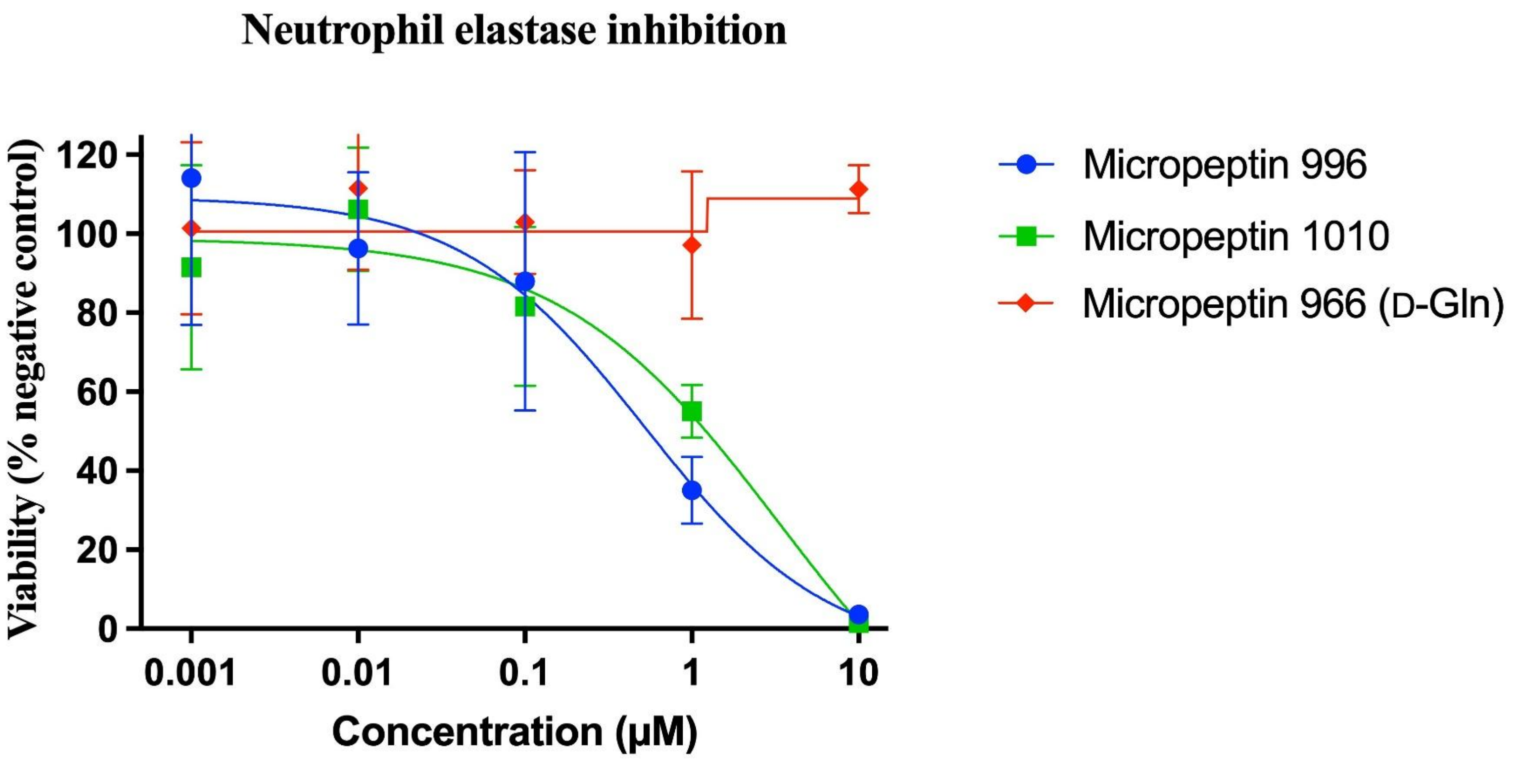


**Figure S30.** Activity of micropeptin 996 (L-Gln), micropeptin 1010 (**1**), and micropeptin 996 (D-Gln) (**2**) against human neutrophil elastase.

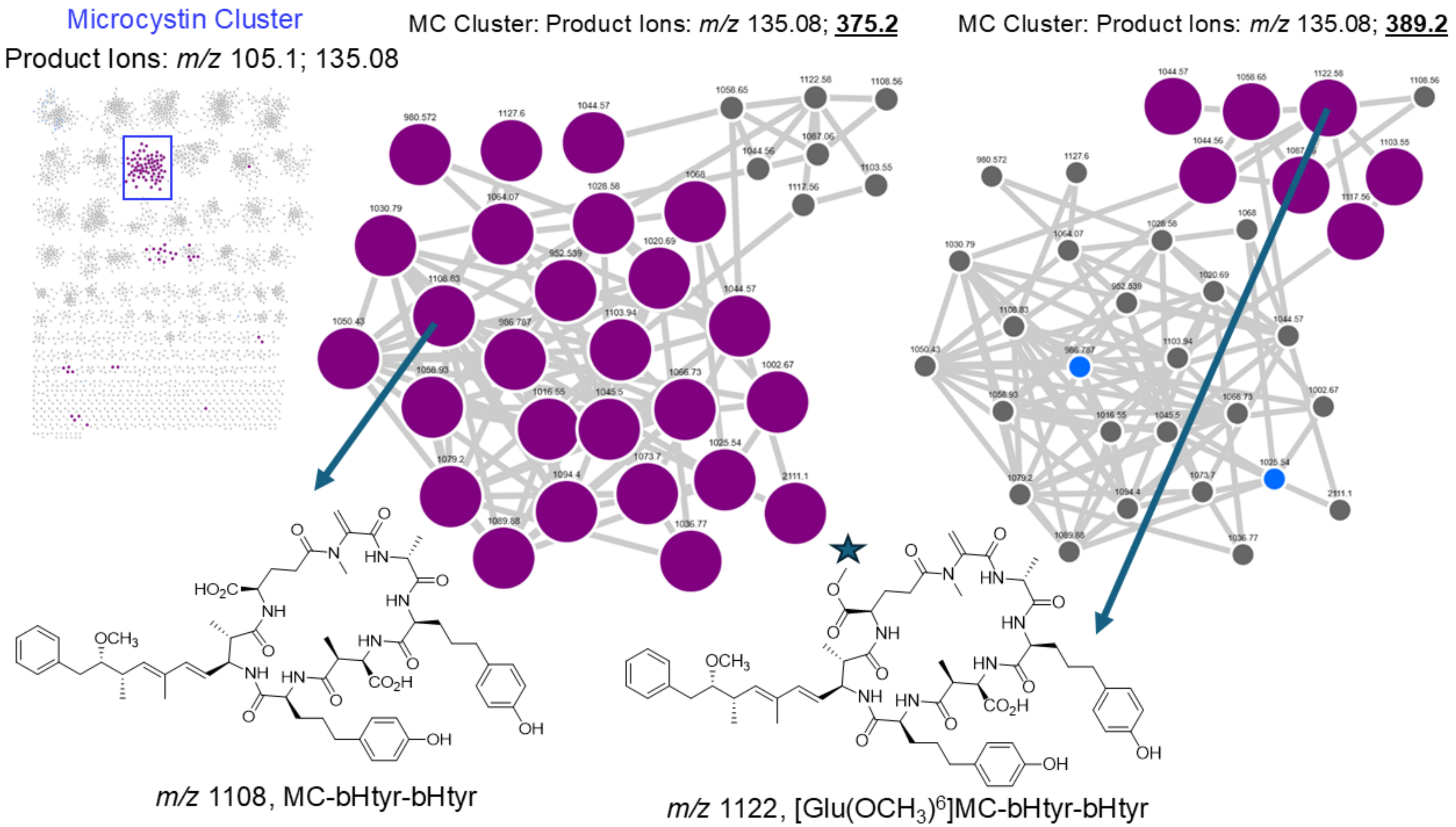


**Figure S31.** Microcystin cluster in MS/MS molecular network subjected to two different product ion searches: *m/z* 135.08 and *m/z* 375.2 and then *m/z* 135.08 and *m/z* 389.02 to illustrated microcystins with likely [Glu($OCH_3$)]$^6$ modifications, which is also supported by the annotation of library microcystins shown.

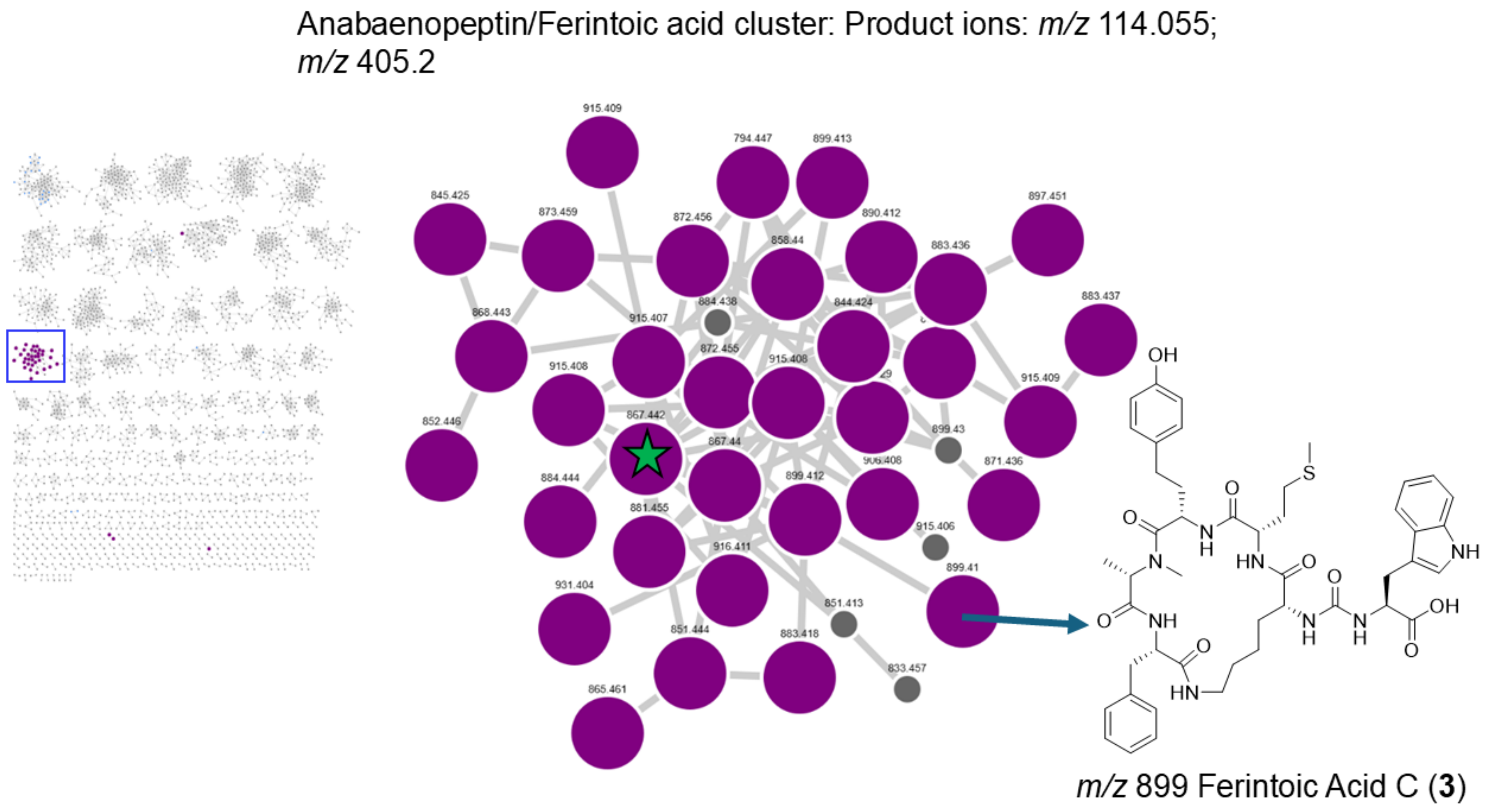


**Figure S32.** MS/MS cluster of anabaenopeptins/ferintoic acids annotated via product ion searching. Ferintoic acid A (green star) and ferintoic acid C (**3**) were validated using our standard library.

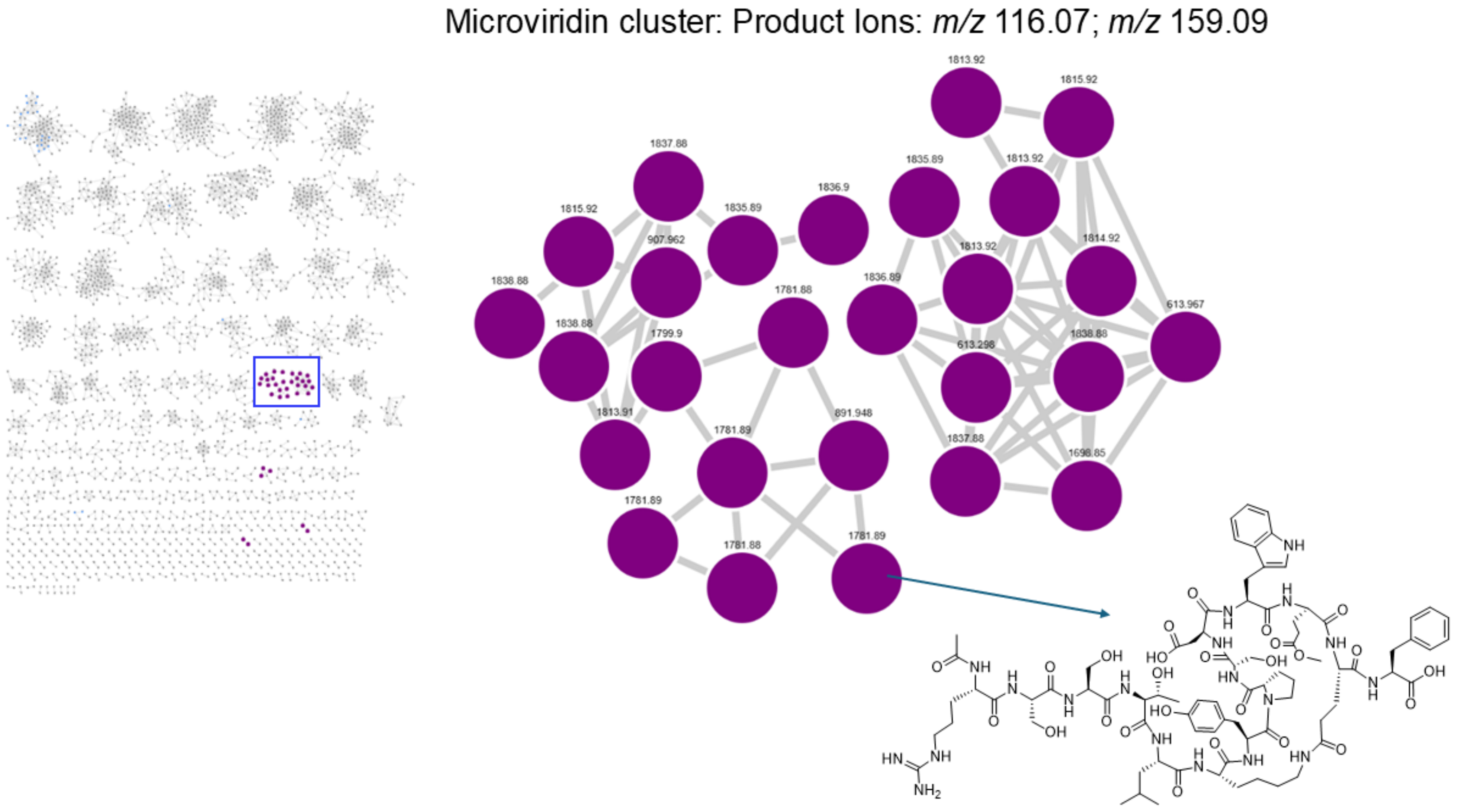


Microviridin 1781

**Figure S33.** Microviridin cluster annotated using product ion searching in MS/MS network. Standard compound microviridin 1781 was used for validation.

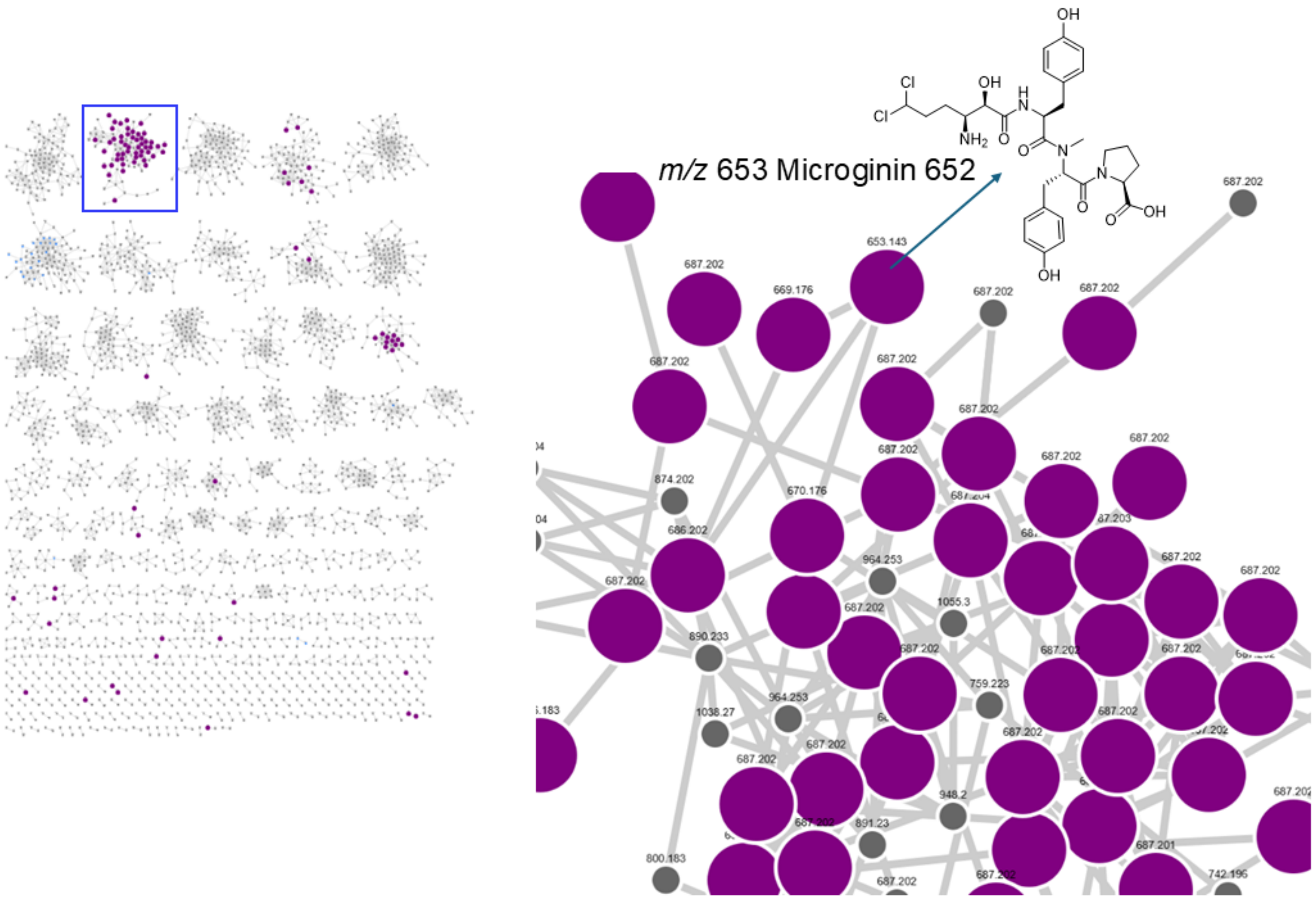


**Figure S34.** Microginin cluster annotated using product ion searching in MS/MS networks (see main text for product ion search details). Standard compound microginin 652 was used for validation.

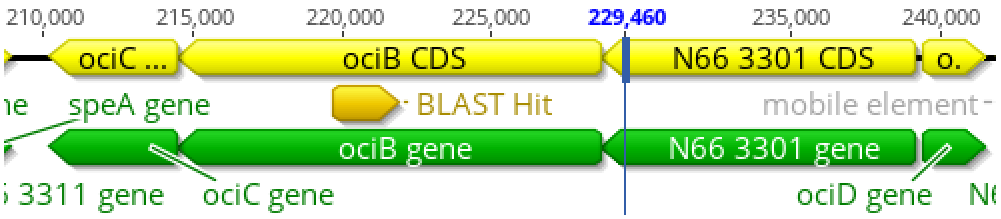


**Figure S35.** Blast hit to the *ociB* gene (cyanopeptolin biosynthetic pathway) in the metagenomic sequence data from Lake Erie (Miller Road Park).

REFERENCES

1) Fujii, K.; Sivonen, K.; Kashiwagi, T.; Hirayama, K.; Harada, K. -I. Nostophycin, a Novel Cyclic Peptide from the Toxic Cyanobacterium *Nostoc* sp. 152. *J. Org. Chem*. **1999**, *64* (16), 5777–5782.
